# Multi-state kinetics of the syringe-like injection mechanism of Tc toxins

**DOI:** 10.1101/2024.01.16.575634

**Authors:** Peter Njenga Ng’ang’a, Julian Folz, Svetlana Kucher, Daniel Roderer, Ying Xu, Oleg Sitsel, Alexander Belyy, Daniel Prumbaum, Ralf Kühnemuth, Tufa E. Assafa, Min Dong, Claus A. M. Seidel, Enrica Bordignon, Stefan Raunser

**Affiliations:** Department of Structural Biochemistry, Max Planck Institute of Molecular Physiology, Otto-Hahn-Str. 11, 44227 Dortmund, Germany; Chair of Molecular Physical Chemistry, Heinrich-Heine University of Düsseldorf, Universitätsstr. 1, 40225 Düsseldorf, Germany; Department of Physical Chemistry, University of Geneva, Quai Ernest Ansermet 30, 1211 Geneva, Switzerland; Faculty of Chemistry and Biochemistry, Ruhr University Bochum, Universitätsstr. 150, 44801 Bochum, Germany; Leibniz-Forschungsinstitut für Molekulare Pharmakologie, Robert-Roessle-Str. 10, 13125 Berlin, Germany; Department of Urology, Boston Children’s Hospital, Boston, MA, USA; Department of Surgery and Department of Microbiology, Harvard Medical School, Boston, MA, USA

## Abstract

Tc toxins are virulence factors of many insects and human pathogenic bacteria. They attach as soluble prepores to receptors on host cells and following acidification in the late endosome, perforate the cell membrane like a syringe to translocate toxic enzymes into the host cell through their pore-forming channel. Although this complex transformation has been structurally well studied, the functional aspects of this large-scale rearrangement, such as the reaction pathway with possible intermediate states and the resulting temporal evolution have remained elusive. Here, we used an integrated biophysical approach to monitor the prepore-to-pore transition and found that it takes ∼28 h when induced by high pH in the absence of other factors. In the presence of liposomes, an increasingly high pH or receptors, such as heparin or Vsg, the probability to transform prepores to pores increases by a factor of up to 4. This effect can also be mimicked by biotinylation or site-directed mutagenesis of the shell, demonstrating that shell destabilization is a crucial step in prepore-to-pore transition. We show that shell opening is a heterogeneous process with transition times ranging from 60 ms to 1.6 s and resolve three sequential intermediate states: an initial transient intermediate during shell destabilization, a first stable intermediate where the receptor-binding domains on the shell rearrange and a second stable intermediate with an open shell. In contrast, the ejection of the pore-forming channel from the open shell is highly cooperative with a transition time of < 60 ms. This detailed knowledge of the Tc toxin mechanism of action, even in the absence of receptors, is important for the future application of Tc toxins as biomedical devices or biopesticides.

## Introduction

Toxin complexes (Tc) are pathogenicity factors in various bacteria that infect insects or mammals^1–3^. While Tcs from insect pathogenic bacteria, such as *Photorhabdus luminescens*^4^ or *Xenorhabdus nematophila*^5^, show potential for biotechnological applications^6^, Tcs of human pathogenic bacteria such as *Yersinia pseudotuberculosis*^7^ or *Yersinia enterocolitica*^8^ may also play an important role in infectious diseases.

Tc toxins use a special syringe-like mechanism to perforate the host cell membrane and inject a deadly enzyme into the host cytosol. They are composed of three components: TcA, TcB and TcC^9,10^. TcA is the key component for receptor binding, pore formation and translocation of the toxin^11,12^. It forms a bell-shaped homopentamer of ∼1.4 MDa made up of a central, α-helical channel surrounded by a shell, to which it is connected by a stretched linker^13^. The receptor-binding domains (RBDs) at the surface of the shell are structurally variable^14^, allowing the association of various Tc toxins to different receptors, such as glycans^12,15–17^ or highly glycosylated protein receptors such as Visgun (Vsg)^18^.

TcB and TcC together form a hollow cocoon of ∼250 kDa that encapsulates the ∼30 kDa C-terminal domain of TcC^19,20^, also known as the hypervariable region (HVR). The HVR is the actual toxic enzyme that is autoproteolytically cleaved inside the cocoon. In the case of TccC3 from *P. luminescens,* the HVR is an ADP-ribosyltransferase, that modifies F-actin at threonine-148^21–23^ leading to disorganization of the target cell’s cytoskeleton. The TcA pentamer and TcB-TcC heterodimer spontaneously assemble with sub-nanomolar affinity to form the ∼1.7 MDa ABC holotoxin^24^.

In the prepore form of the holotoxin, the shell of TcA is closed. A pH change to either acidic^23^ or basic^11,25^ conditions results in the opening of the shell. Subsequently, the stretched linkers between the shell and the channel condense rubber-band-like from an extended, unfolded conformation to a compact, partially folded conformation. As a consequence, the channel is ejected and enters the target cell membrane similar to a syringe. The hydrophobic environment of the membrane triggers the opening of the tip of the channel. In this so-called pore state, the HVR is translocated through the channel into the target cell^11^.

The architectures of the prepore and the pore state of Tc toxins have been well examined, elucidating the starting and end point of the transition from prepore to pore in molecular detail^24,26^. However, functional aspects of this large-scale rearrangement of the pentamer, such as the reaction pathway and its energy landscape with possible intermediate states, synchronization of the protomers and the resulting temporal evolution are still unknown. This limits our understanding of Tc toxin action and makes it difficult to tailor Tc toxins for biotechnological applications. Here, we combine electron paramagnetic resonance (EPR) spectroscopy, confocal and total internal reflection (TIR) single-molecule fluorescence spectroscopy, single particle electron cryo microscopy (cryo-EM) and biochemistry to address these questions. We examine the Tc toxin from the *Photorhabdus luminescens* strain W14, comprising TcA (TcdA1), TcB (TcdB2) and TcC (TccC3) and demonstrate that the prepore-to-pore transition of this Tc toxin at high pH involves stable intermediate states, in which the RBDs of the shell rearrange and the shell opens. Additionally, we show that the transition from prepore to pore has a reaction time of ∼28 h *in vitro*, however, an individual channel ejection event proceeds within several milliseconds. This discrepancy originates from a low probability of the reaction to be initiated *in vitro*. In the presence of liposomes, an increasingly high pH or a receptor, such as heparin or Vsg, the probability increases by a factor of up to 4, an effect which can also be mimicked by biotinylation or site-directed mutagenesis, indicating that destabilization of the shell is essential for the quick transition from prepore to pore. Our results are a key contribution to a holistic understanding of the complex mechanism of action of molecular machines, in particular Tc toxins and provide an important basis for their adaptation to biotechnological applications.

## Results

### Receptor binding accelerates pH-dependent shell opening

We and others have previously shown that TcA binds to target cells via glycans or glycosylated surface receptors and transitions from prepore to pore, before delivering its toxic payload into the cytoplasm^11,12,15,17,18,27^. During prepore-to-pore transition, TcA undergoes two major conformational changes: shell opening and channel ejection (Figure 1a). However, there are no intermediate states known that explain whether these steps proceed simultaneously or sequentially and no data are available on the kinetics of the reaction. We therefore set out to address the individual reaction kinetics of shell opening and channel ejection by EPR spectroscopy. In this regard, we designed TcA variants with the point mutations Q914C, D1193C and S2365C that allowed 3-(2-iodoacetamido)-proxyl (IAP)-labelling to measure spin label distance changes that occur during shell opening and channel ejection in a time-resolved manner (Figure 1b-c, Supplementary Figure 1). Both IAP-labelled toxins were still active against HEK 293T cells (Supplementary Figure 1e), showing that site-directed mutagenesis and labelling did not interfere with holotoxin assembly, cell binding, prepore-to-pore transition and toxin translocation.

**Figure 1:**
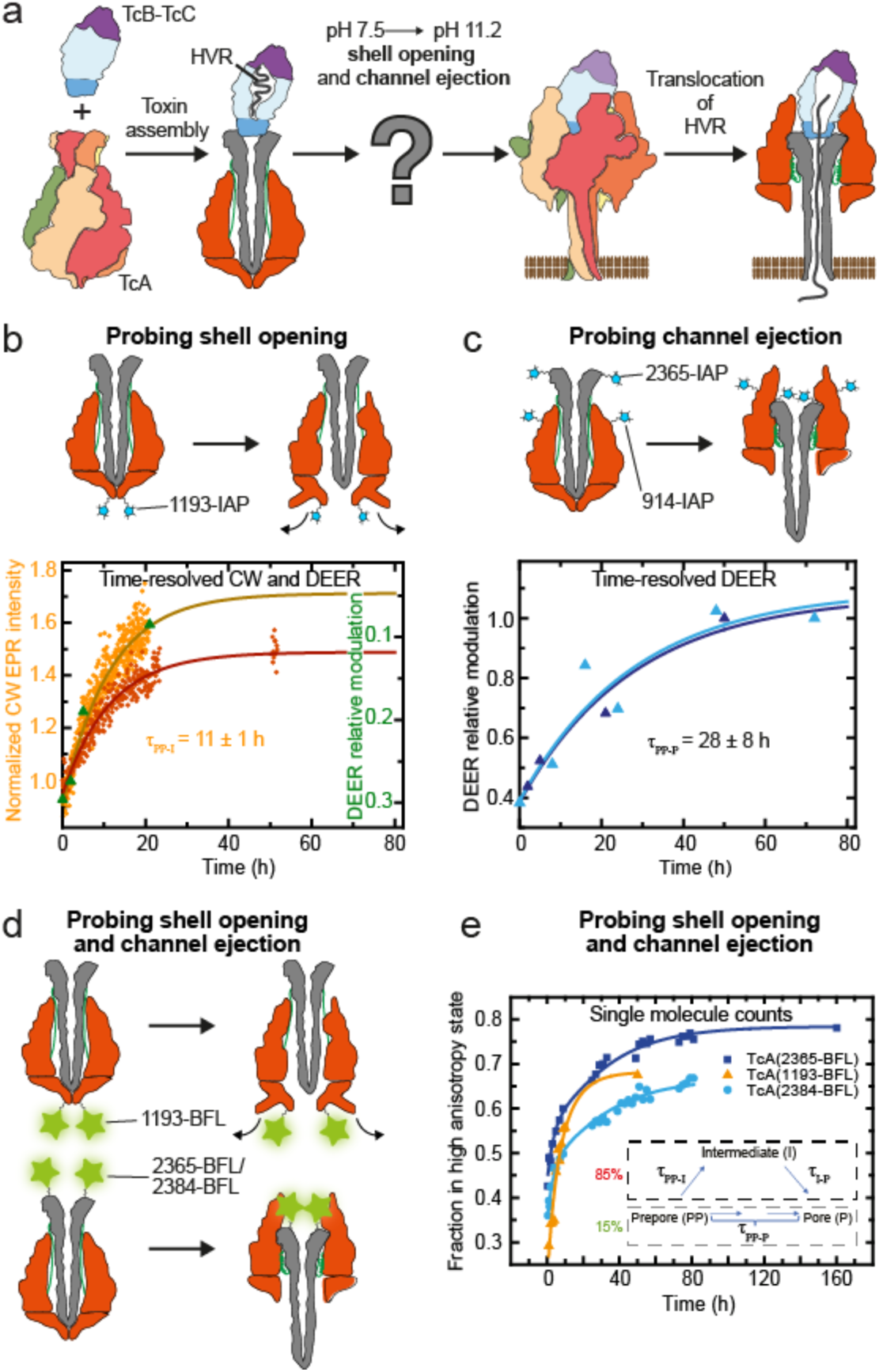
Kinetics of the individual steps of prepore-to-pore transition of TcA. **a:** Schematic overview of the functional mechanism of action of Tc toxins. The toxins are shown as both surface cartoon representations (left) and vertical cut-throughs (right). TcB, TcC and the protomers of TcA are differently colored. **b: Top panel**: design of a TcA variant to probe opening of the shell via EPR spectroscopy. TcA(1193Cys), was labeled with 3-(2-iodoacetamido)-proxyl IAP, resulting in TcA(1193-IAP). **Bottom panel**: CW EPR kinetics at pH 11.2 showing a first order reaction obtained from a global fit of two biological replicates (orange and red curves) and DEER relative modulation kinetics (green data points). **c: Top panel**: design of a TcA variant to probe channel ejection via EPR spectroscopy: TcA(914Cys/2365Cys) was labeled with IAP at both cysteines, resulting in TcA(914-IAP/2365-IAP). **Bottom panel**: EPR DEER kinetics at pH 11.2 showing a first order reaction obtained from a global fit of two biological replicates (light and dark blue data points). **d:** Design of TcA variants to probe shell opening and channel ejection via single molecule fluorescence anisotropy. TcA(1193Cys), TcA(2365Cys) and TcA(2384Cys) were labeled with Bodipy-FL-iodacetamide (BFL), resulting in TcA(1193-BFL), TcA(2365-BFL), and TcA(2384-BFL), respectively. **e:** Confocal single molecule fluorescence anisotropy experiments at pH 11.2. The time dependence for shell opening was fitted using a monoexponential function (orange curve). For channel ejection, a complex model (see inset and justification see Supplementary Note 1 and Figure SN1) with a consecutive path including one intermediate state and a parallel direct path were needed to describe all three data sets with global rate constants (blue and cyan curves for TcA(2365-BFL) and TcA(2384-BFL), respectively).

We first focused on the opening of the shell using TcA-1193IAP. We induced the prepore-to-pore transition by a pH shift to 11.2 and monitored the kinetics of the conformational changes via continuous wave (CW) EPR at room temperature and the associated changes in the inter-protomer distances via double electron electron resonance (DEER) measurements on samples snap frozen after defined incubation times. Time-resolved CW EPR kinetics revealed that shell opening correlates with increasing label mobility and proceeds as a first order reaction with a reaction time of 11 ± 1 h (Figure 1b). The expected increase in interspin distances due to shell opening (Figure 1b, Supplementary Figure 2) consistently followed the first order kinetics obtained with CW EPR. EPR experiments performed after a shift to pH 4 confirmed the absence of shell opening at acidic pH *in vitro* (Supplementary Figure 2a). Interestingly, slight basic pH changes had a significant impact on the kinetics and decreased the time of shell opening by an order of magnitude between pH 11 and pH 11.8 (Supplementary Figure 3b,c).

Expectedly, holotoxin assembly, that is the binding of the TcB-TcC cocoon onto the TcA pentamer, had no major influence on the transition kinetics (Supplementary Figure 3c,f). In contrast, the shell opening was accelerated by an order of magnitude when liposomes were added instead of detergents (Supplementary Figure 3d-f). This is surprising, as membrane insertion and thus direct contact with the membrane occurs at a later stage and indicates that the interaction of the toxin with lipids destabilizes the shell and accelerates its opening. A similar acceleration through interaction of Tc toxins with the host membrane *in vivo* is therefore conceivable.

Tc toxins can potentially penetrate host cell membranes in the midgut directly, where shell opening is probably triggered by the high pH in this part of the insect digestive tract^28^. However, the more common route of Tc toxin intoxication is via the endosomal pathway, where pore formation happens after acidification of the late endosome^29^. So far, we could only reproduce the former *in vitro*, whereas the latter seems to depend on additional factors, such as binding to receptors. In order to find out whether binding of heparin, Lewis X glycans, or Vsg, which are known receptors of TcdA1 from *P. luminescens*^12,15,17,18^, has an influence on the pH dependence of shell opening, we incubated ABC holotoxins with the receptors at either pH 4 or pH 11.2 and monitored pore formation by negative stain EM (Figure 2a, Supplementary Figure 4). At pH 4, none of the receptors showed an effect compared to the control (Supplementary Figure 4c,e,g). At pH 11.2, Lewis X had no effect (Figure 2a, Supplementary Figure 4d), but interaction with Vsg and heparin caused a 3-times and 4-times increase in pore formation rates, respectively (Figure 2a, Supplementary Figure 4a,f). This indicates that heparin and Vsg play an important role when Tc toxins penetrate directly into host cell membranes and suggests that still other factors, such as for example yet unknown protein receptors, or a combination of different glycans and protein receptors are needed for Tc toxins acting through the endosomal pathway.

**Figure 2:**
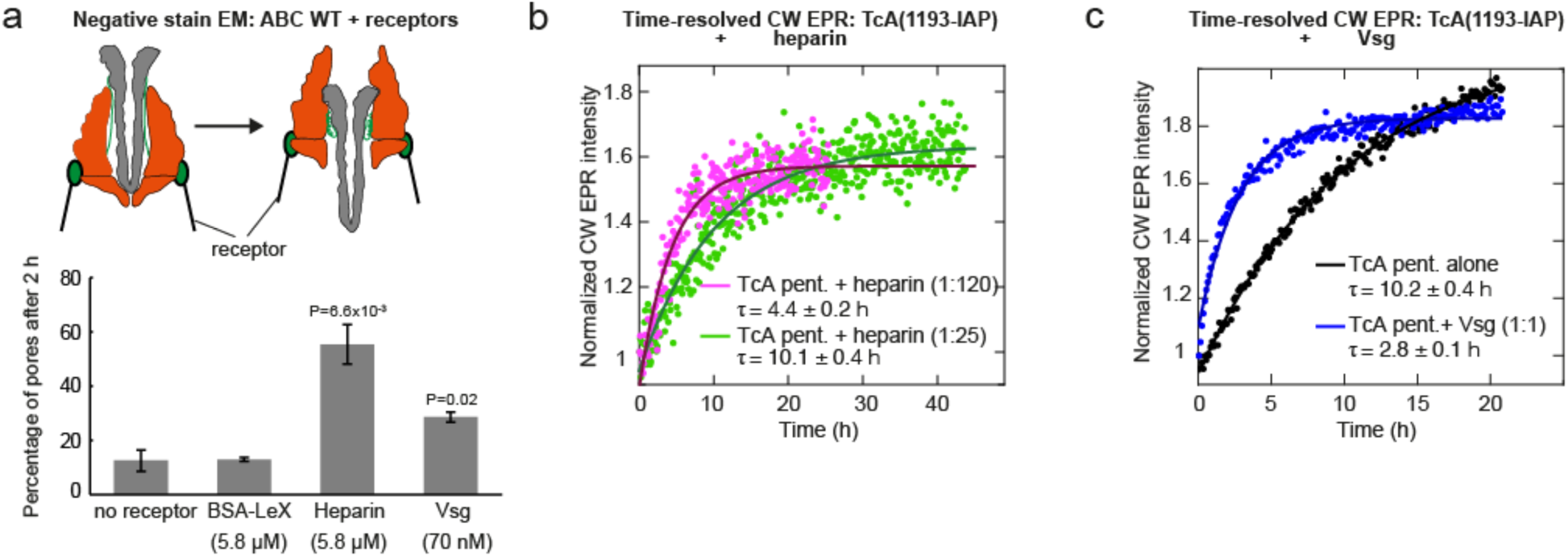
Effect of receptors on prepore-to-pore transition kinetics. **a:** Negative stain EM analysis of pore formation of ABC WT at 70.2 nM pentamer concentration after incubation for 2 h at pH 11.2 and 21 °C, in the presence or absence of the receptors Lewis X (BSA-LeX), heparin and Vsg at the indicated concentrations. The data are presented as mean values and the error bars represent the standard error of the mean (SEM) of three independent experiments. **b**: CW EPR kinetics of TcA(1193-IAP) at 2.8 µM TcA pentamer concentration, pH 11.2 and 21 °C in the presence of 71 and 333 µM of heparin. The TcA pentamer to heparin molar ratios are indicated. **c**: CW EPR kinetics of TcA(1193-IAP) at 3 µM TcA pentamer concentration at pH 11.2 and 21 °C in the presence of 3 µM Vsg. The TcA pentamer to Vsg molar ratios are indicated.

To find out if heparin binding influences shell opening or other steps in the prepore-to-pore transition, we preincubated TcA(1193-IAP) with heparin at different concentrations and then analysed the kinetics by CW EPR. A 120-fold molar excess of heparin to the TcA pentamer resulted in a 2.3-fold decrease in the opening time of the toxin, indicating that heparin indeed accelerates shell opening (Figure 2b, Supplementary Figure 4h,i). The effect of the protein receptor Vsg is higher, since preincubation of TcA(1193-IAP) with Vsg at a molar ratio of 1:1 to the TcA pentamer resulted in a 3.6-fold decrease in the opening time of the toxin (Figure 2c). Thus, binding of Vsg or heparin to TcA is not only involved in docking onto target cells, but it also potentiates the toxin for pH-induced shell opening at high pH. Taken together, we conclude that the slow shell opening of TcA *in vitro* can be accelerated by increasingly high pH, liposomes or receptor binding.

### Prepore-to-pore transition proceeds via intermediate states

To study the kinetics of channel ejection in the prepore-to-pore (PP-P) transition, we performed DEER studies with TcA(914-IAP/2365-IAP) at different incubation times. However, this experiment is unable to probe shell opening and the measured reaction times therefore correspond to the entire transition from prepore to pore (named τPP*-P; PP* in the following will denote the prepore state after the pH shift). The shift from pH 7.5 to pH 11.2 resulted in the expected decrease in interspin distance (Supplementary Figure 5) and the DEER kinetics showed that prepore-to-pore transition proceeds with an approximate reaction time τPP*-P of 28 ± 8 h (Figure 1c). Intriguingly, the pore formation kinetics obtained with negative stain EM data on both TcA(1193-IAP) and TcA(914-IAP/2365-IAP) showed a τPP*-P of 10 ± 2 h and 11 ± 2 h, respectively (Supplementary Figure 6), consistent with the kinetics of the shell opening detected in solution. This indicates that the pore formation process of the fraction of toxins trapped on the grid follows the first-order kinetics of the shell opening detected in solution. Therefore, in solution, the transition from prepore to pore is slower (> 20 h) than the opening of the shell (11 ± 1 h), indicating that there is at least one stable intermediate state (I) between shell opening and channel ejection, which is not captured by negative stain EM.

To further corroborate these experiments in solution and at room temperature, we used TcA variants with the fluorescence label Bodipy-FL (BFL) and performed single-molecule Förster resonance energy transfer (FRET) between fluorophores of a single type (homoFRET), probing fluorescence anisotropy. We monitored shell opening ((TcA(1193-BFL) – same labelling position as in EPR) and channel ejection ((TcA(2365-BFL) and TcA(2384-BFL) – different labelling positions as in EPR) (Figure 1d,e, Supplementary Figure 7). The kinetics of shell opening followed a simple monoexponential path, in agreement with the EPR data. However, the single-molecule anisotropies of TcA(2365-BFL) and TcA(2384-BFL) showed kinetics of pore formation (see Methods) which cannot be fitted by a simple single exponential and thereby confirmed the presence of at least one stable prepore-to-pore transition intermediate (I). Integration of data from all three variants using a simple consecutive model with an intermediate I (Figure 1e) gave a reaction time for the prepore-to-intermediate transition (τPP*-I) of 7 ± 1 h and a second reaction time for the intermediate-to-pore transition (τI-P) of 27 ± 5 h (Figure 1e). Small deviations at short times indicate the presence of a small fraction of TcAs (15%) with a direct fast transition from prepore to pore of τPP*-P = 3.7 ± 0.1 h, as outlined in Supplementary Note 1. Thus, in agreement with the EPR data, the single-molecule data showed faster kinetics for shell opening than for the overall pore formation. In addition, they revealed that the conformational transitions in TcA are complex and kinetically heterogeneous.

### Slow shell opening precedes fast channel ejection

Next, we set out to measure how fast individual Tc toxins open their shell and eject their channels by single-molecule FRET in a TIRF setup. We designed Tc variants that probe shell opening and channel ejection by a change in FRET efficiency. First, TcA labelled with Atto647N-iodacetamide at position C1279 (TcA(1279-At647N)) was used to probe shell opening by homoFRET. Secondly, we used a combination of a BFL-labelled TcA (TcA(914-BFL)) and an Atto647-labelled TcB-TcC (TcB(1041-At647N)-TcC) to trace channel ejection by heteroFRET (Figure 3a, Supplementary Figure 8). To immobilize both Tc toxins on neutravidin-coated cover slips, we biotinylated them and monitored the transition times after a pH shift to pH 11.2 by FRET (Figure 3b). Markedly, the reaction times of the biotinylated TcA variants were about four orders of magnitude faster than the times observed initially (Supplementary Figure 9), indicating that biotin labelling has a destabilizing effect on the shell at pH 11.2 and accelerates shell opening. Yet, the kinetics of the overall pore formation were slower than shell opening, indicating that at least one intermediate was still present.

**Figure 3:**
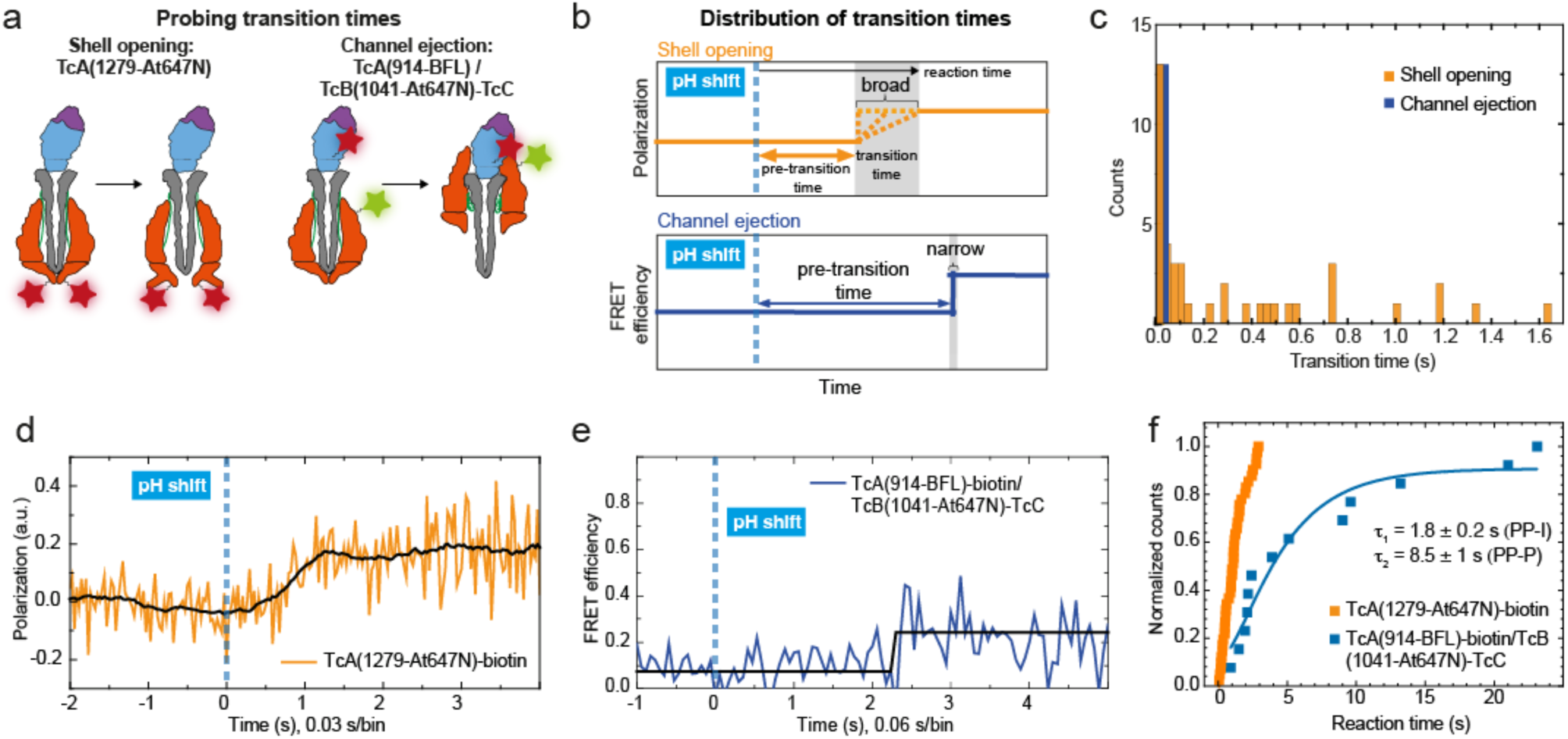
Single-molecule fluorescence studies of shell opening and channel ejection of surface immobilized toxins. **a**: Design of TcA and TcB-TcC variants to probe single shell opening and channel ejection events. **b:** Schematics of fluorescence traces probing shell opening (orange) and channel ejection (blue). After a pH jump from 7.4 to 11.2 (dashed blue line, time =0) the toxin undergoes a transition from the prepore to pore state (shaded in gray) after a specific pre-transition time (double-sided arrows). **c**: Single molecule fluorescence homoFRET experiments using biotinylated TcA(1279-At647N) to assess shell opening and channel ejection. Shell opening experiments revealed a distribution of the recorded transition times over 1.6 s (orange bars), whereas those of channel ejection showed transition times within a single frame, below 60 ms (blue bar). **d:** A representative shell opening trace using biotinylated TcA(1279-At647N) which reveals transition times longer than the bin width of 29.44 ms and a change in polarization caused by a difference in homoFRET and reduction of dye mobility due to the moving RBDs. a.u. = arbitrary units. **e:** A representative trace of the channel ejection heteroFRET assay using ABC with a donor dye at TcA(914-BFL) and acceptor dye at TcB(1041-At647N)-TcC. The displayed trace shows a fast transition of the signal from the prepore to pore state within one bin (bin width=58.88 ms). **f**: Reaction times of immobilized, biotinylated TcA(1279-At647N) (orange) and ABC composed of TcA(914-BFL)-biotin and TcB(1041-At647N)-TcC (blue). PP = prepore, I = intermediate, P = pore.

We recorded 42 shell opening traces of single Tc toxins and measured transition times from 60 ms to 1.6 s after an initial lag-phase (pre-transition time) following pH change (Figure 3b,c,d, Supplementary Figure 10-11, Methods). The distribution of transition times was quite broad, indicating that shell destabilization and opening did not happen abruptly, but involved small intermediate steps instead. Notably the pre-transition and reaction times are correlated in duration (Supplementary Figure SN5a-c) so that both processes (Figure 3f) influence the overall reaction time.

For channel ejection, we monitored 13 single reactions, all of which showed a transition time within the temporal resolution of the experimental setup (60 ms; Figure 3b,c,e, Supplementary Figure 10-11). This demonstrates that channel ejection is a spontaneous reaction, resulting from a concerted compaction of all five linkers. The pre-transition phase after pH change was longer for channel ejection than for shell opening (Figure 3b), indicating that the channel is only released after the shell has opened. Thus, the overall reaction time is essentially determined by the pre-transition time (Figure 3f).

### Shell destabilization accelerates prepore-to-pore transition

To understand the accelerating effect of biotinylation on prepore-to-pore transition, we produced mimics of biotinylation by site-directed mutagenesis. To this end, we specifically focused on electrostatic interactions that involve surface-exposed lysines (NHS-biotin targets) near protomer-protomer interfaces of the TcA pentamer. We chose promising lysines at three different positions of the protein complex: K567 on the alpha-helical domain of the shell, K1179 on the neuraminidase-like domain near the tip of the channel and K2008 on the linker (Supplementary Figure 12a). Although none of these residues is conserved in the Tc toxin family, they are involved in polar and electrostatic interactions potentially stabilizing the conformation of the prepore state of TcdA1 from *P. luminescens* (Supplementary Figure 12b). Since biotinylating these lysines would neutralize their charges, we mutated the chosen lysines to tryptophan, which also mimics the size increase by biotinylation.

We first focused on the mutants K567W and K2008W and counted the number of pores formed after 2 h of incubation at pH 11.2 by negative stain EM. Indeed, both mutations had an accelerating effect, and the number of pores was three to four times higher compared with the wild type. The double-mutant K567W/K2008W did not have the expected amplifying effect and the number of pores were comparable to the K2008W mutant (Figure 4a, Supplementary Figure 12c-d). In line with the EM study, the K567W/K2008W double mutant showed three-times faster kinetics of shell opening in EPR measurements compared to the wild type (Figure 4b, Supplementary Figure 13c). Interestingly, while the mutation K567W had no effect on the speed of intoxication of HeLa cells, intoxication with K2008W was considerably faster (Supplementary Figure 12e). Together, our results demonstrate that the accelerating effect of biotinylation can indeed be mimicked *in vitro* and *in vivo* by point mutations that destabilize the shell at different positions.

**Figure 4:**
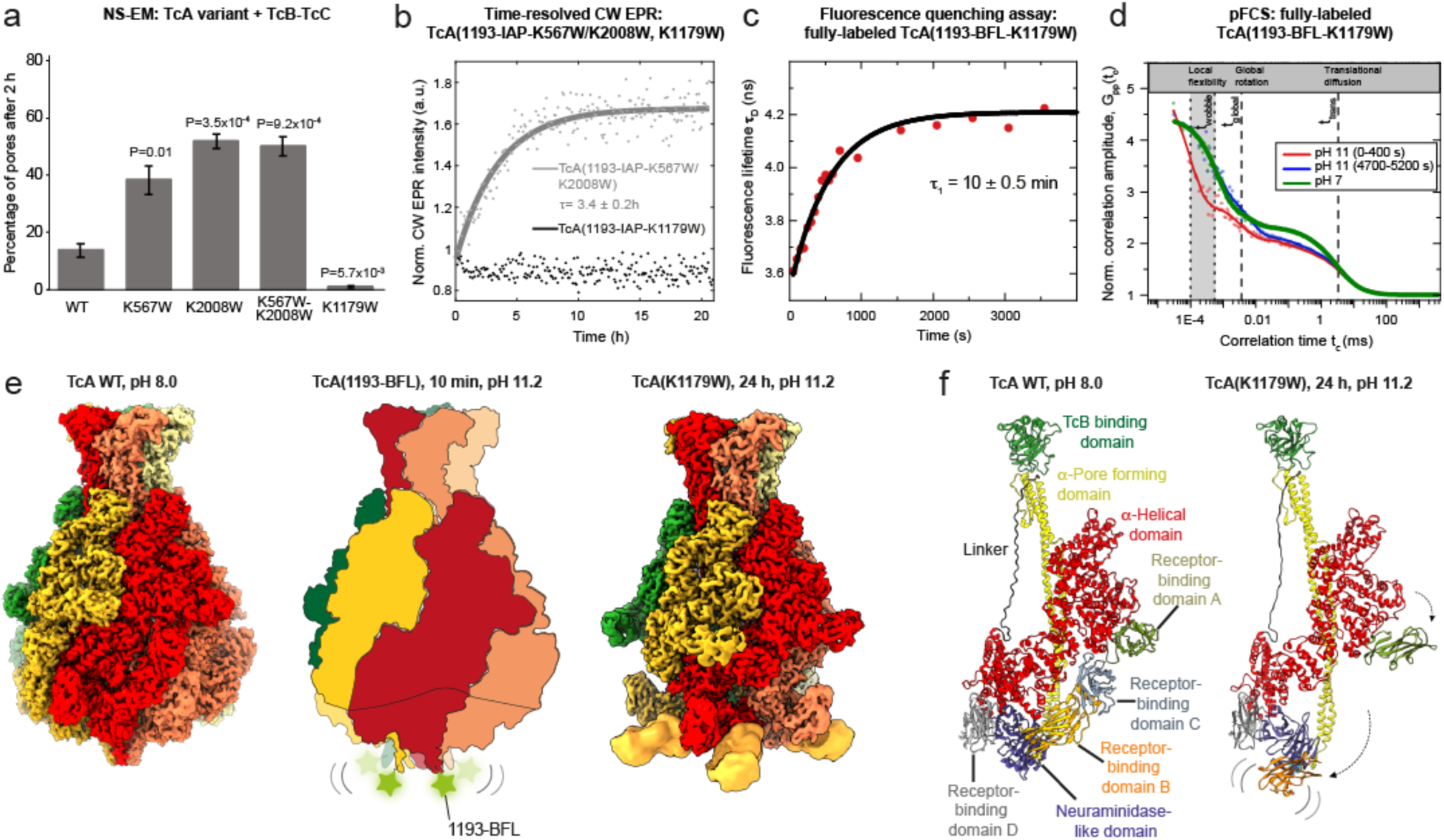
TcA prepore-to-pore stable intermediate kinetics and cryo-EM structure. **a**: Negative stain EM analysis of pore formation of the indicated K-to-W TcA variants with TcB-TcC after incubation for 2 h at pH 11.2 and 21 °C. The data are presented as mean values and the error bars represent the standard error of the mean (SEM) of three independent experiments. **b**: CW EPR kinetics of TcA(1193-IAP-K567W/K2008W) and TcA(1193-IAP-K1179W) at pH 11.2 showing that the increase in dynamics probing shell opening proceeds according to a first order reaction with a reaction time of 3.4 ± 0.2 h for TcA(1193-IAP-K567W/K2008W) while it does not occur for TcA(1193-IAP-K1179W). The latter shows only a small decrease in the label’s dynamics within the first hour. **c**-**d**: Multiparameter fluorescence detection (MFD) experiments analyzed by polarization-resolved fluorescence correlation spectroscopy (pFCS) and anisotropy measurements of TcA(1193-BFL-K1179W) at pH 11.2, long incubation (**c**) and shorter incubation (**d)**. **d**: pFCS resolves the fast-wobbling dynamics of TcA(1193-BFL-K1179W). Experimental data (dots) and fit curves (full lines, see Methods) for the autocorrelation amplitude G_pp_(t_c_) of parallel polarized fluorescence of TcA(1193-BFL-K1179W) at different incubation times and pH conditions: incubation time 0-400 s at pH 11 (red curve), incubation time 4700-5200 s at pH 11 (blue curve), and for comparison at pH 7 (green curve) (see Supplementary Note 2). **e**: **Left panel**: cryo-EM density map of TcA WT at pH 8.0 (EMDB 10033). The five protomers are colored individually**. Middle panel**: schematic representation of TcA(1193-BFL) incubated for 10 minutes at pH 11.2. **Right panel**: cryo-EM density map of TcA(K1179W) intermediate state (SI1) after 24 h of incubation at pH 11.2 and 21 °C. The five protomers are colored individually and the extra density observable at low binarization threshold (Supplementary Figure 14f) is shown in orange. **f**: Comparison of a protomer of the TcA WT prepore (**left panel**, PBD 6RW6) and of the TcA(K1179W) (**right panel**) prepore-to-pore intermediate.

We then analysed the mutant K1179W. Intriguingly, K1179W had the opposite effect to the other mutants and we almost did not find any pores after 2 h (Figure 4a, Supplementary Figure 12c) and time-resolved EPR measurements showed that the TcA shell did not open significantly (Figure 4b, Supplementary Figure 13e,f). In addition, this mutant was not able to intoxicate HeLa cells (Supplementary Figure 12e). Thus, K1179W stabilizes the closed state of the shell independent of the pH.

However, a fluorescence quenching assay (Figure 4c) and polarisation-resolved fluorescence correlation spectroscopy (pFCS)^30^ (Figure 4d) of the BFL-labelled K1179W variant (TcA(1193-BFL-K1179W)) revealed a near-instantaneous (10 ± 0.5 min) local conformational change of the labelled neuraminidase-like domain with a large angle of motion and an unique fast wobbling time t_wobble_ of 100 ns, which is approximately one order of magnitude faster than the global rotational diffusion time t_global_ ≈ 2.2 µs for the toxin particle (1.4 MDa TcA) (see Methods). The detection of the dequenching and this transient local flexibility is related to the pH-induced destabilization of the tip of the shell, which is essential for shell opening and subsequent channel release. This allowed us to assign an additional intermediate referred to as transient intermediate state, TS1. After 4700 s, this TcA species vanished and a more rigid state with t_wobble_ of 600 ns with a smaller angle of motion was formed, which was stable under our conditions (Figure 4c,d, Supplementary Figure 13g). The initial wobbling could explain the broader distance distribution observed by DEER on TcA(1193-IAP-K1179W) after snap freezing of the sample at pH 11.2 (Supplementary Figure 13f) and the subsequent stabilization is in line with the small decrease in the dynamics within the first hour of incubation detected by CW EPR (Figure 4b). Since this mutant does not form pores, it must reside in a stable intermediate state in which the shell is destabilized but not yet open. Therefore, we call this state stable intermediate 1, SI1. Notably, cryo-EM (see below), pFCS and EPR (Supplementary Figure 13f) found that the neuraminidase-like domain is in its prepore conformation in SI1 corroborating the notion that the initial flexibility in TI is indeed transient.

### Prepore-to-pore intermediate with displaced RBDs

To determine the structure of the stable intermediate state SI1, we performed a single particle cryo-EM analysis of the K1179W mutant after 24 h at pH 11.2. Expectedly, only 1% of the toxin was in the pore state, the rest of the particles resembled prepores but were more slender in appearance (Supplementary Figure 14a,b). Interestingly, 2D classes of these particles appeared to have protruding densities, implying conformational changes (Supplementary video 1). To elucidate the conformation of these particles, we determined their 3D structure at an average resolution of 2.9 Å (Figure 4e, Supplementary Figure 14c,f). The overall conformation is similar to the prepore state and the tip of the shell is closed below the channel (Supplementary Figure 13a). However, RBD A is flipped out by about 20° away from the shell and in line with the slimmer shape, the N-terminal 80 residues and densities corresponding to RBDs B and C were not resolved in the cryo-EM map, indicating that these domains are highly flexible (Figure 4e,f, Supplementary video 2). This was also supported by a variance analysis of the cryo-EM density map, that indicated the presence of disordered regions at the bottom of the outer shell. We therefore applied focused 3D classification and identified a major class with density below the bottom of the shell (Supplementary Figure 14f). The density corresponded to RBD B (Figure 4e,f), but in contrast to the prepore conformation, the domain is flipped by 180° away from the shell. RBD C which is directly connected to RBD B was not resolved, indicating a flexible hinge between the domains. Thus, a shift from pH 7.5 to pH 11.2 indeed induces a conformational change, resulting in a stable intermediate state in which the interaction between the RBDs A, B and C and the shell is destabilized and the RBDs are released from the shell. This stable intermediate would normally proceed further to the pore. However, the tryptophans at position 1179 form strong hydrophobic bonds with neighbouring phenylalanines at position 2145 and 2147 on the channel, which change their orientation from their wild-type conformation (facing down) to that in the mutant (facing up) (Supplementary Figure 13a,b) and thus block the opening of the neuraminidase-like domain for pore formation.

In order to demonstrate that this intermediate state with a destabilized shell and flipped-out RBDs is not only an effect of K1179W, we performed a single particle cryo-EM analysis of a TcA variant designed to probe shell destabilization at single-molecule level (TcA(1279-At647N)) after 6 h at pH 11.2. In this case, 65% of the toxin were in the pore state, but the rest of the particles resembled the intermediate state SI1 observed for the K1179W mutant (Supplementary Figure 15a) and the 3D reconstruction showed the same overall conformation including the flexible rearrangement of the RBDs. Thus, this structure represents indeed a stable intermediate state (Supplementary Figure 15d-f).

An additional cryo-EM structure of the K567W/K2008W double mutant at pH 11.2 also showed the same prepore-to-pore intermediate state, however, the incubation in this case had to be reduced to 2 h due to the faster kinetics of pore formation (Supplementary Figure 16). We thus conclude that there is an intermediate state SI1, in which the shell is destabilized and the RBDs are flipped out and flexible but the shell is still closed. Consequently, there must be also an additional intermediate state SI2 in which, as we have demonstrated by EPR distance changes, the shell is open but the channel is not yet released. Surprisingly, we were unable to trap this state in cryo-EM, suggesting that it is less stable under the plunge-freezing conditions used in cryo-EM. Therefore, to further corroborate the existence of two distinct stable intermediate states, we performed single-molecule homoFRET studies, which showed that the time-resolved anisotropies r(t) of the K1179W mutant and wildtype TcA with the label BFL differ significantly (Supplementary Figure 13g,h, Methods). While the K1179W mutant exhibited only small distance changes the wildtype showed large changes. Thus, we conclude that an intermediate SI2 is formed after the intermediate SI1 and before the channel ejects.

## Discussion

Our integrated biophysical approach combining experimental conditions in solution (fluorescence and CW EPR spectroscopy) and frozen (DEER and cryo-EM) sets a new standard in the understanding of molecular machines by linking structural and kinetic information. We addressed the challenge of getting local information in large multi-subunit protein complexes by specifically attaching probes in subunits to monitor reaction coordinates of interest (shell opening and channel ejection). In this way, local information such as flexibility and fluorescence quenching of the probes were used as a readout in CW EPR as well as fluorescence anisotropy, pFCS and fluorescence lifetime measurements. Moreover, distances between the probes in distinct subunits were monitored by DEER and homoFRET measurements giving information on conformational changes. Importantly, single-molecule fluorescence assays gave insights into the reaction pathways and complemented the ensemble measurements.

This multimodal approach enabled us to describe in detail three intermediate states (one transient and two stable) within the syringe-like injection mechanism of Tc toxins (Figure 5). At neutral pH, the shell of TcA is tightly closed, keeping the channel in its prepore conformation and the RBDs are attached to the alpha-helical domain of the shell. However, upon a pH increase, a transient intermediate (TI) is formed, where the neuraminidase-like domain becomes flexible for approximately 400 s. The flexibility of the domain is characterized by a wobbling time of 100 ns in pFCS, which is approximately one order of magnitude faster than the global rotational diffusion time of the toxin particle. This allowed us for the first time to measure flexibility in a large protein complex (1.4 MDa) in a time range that is to our knowledge experimentally inaccessible to other methods. The transient flexibility of the neuraminidase-like domain is potentially transmitted to the rest of the shell. This initial step is key for the release and flipping out of the RBDs by disrupting intra- and inter-protomer contacts, and leads to the stable intermediate 1 (SI1). Here, the shell is still closed (Figure 4e). To our knowledge, this rearrangement of the RBDs in an intermediate state is unique for Tc toxins. The RBDs might become accessible to receptors on the host membrane through this flipping movement. However, it is also conceivable that the movement of the RBDs bound to membrane-anchored receptors would result in a lever-like opening of the shell *in vivo*. In any case, we found direct evidence that activation in form of a pH increase is necessary to induce flexibility that is a prerequisite for subsequent conformational transitions.

**Figure 5:**
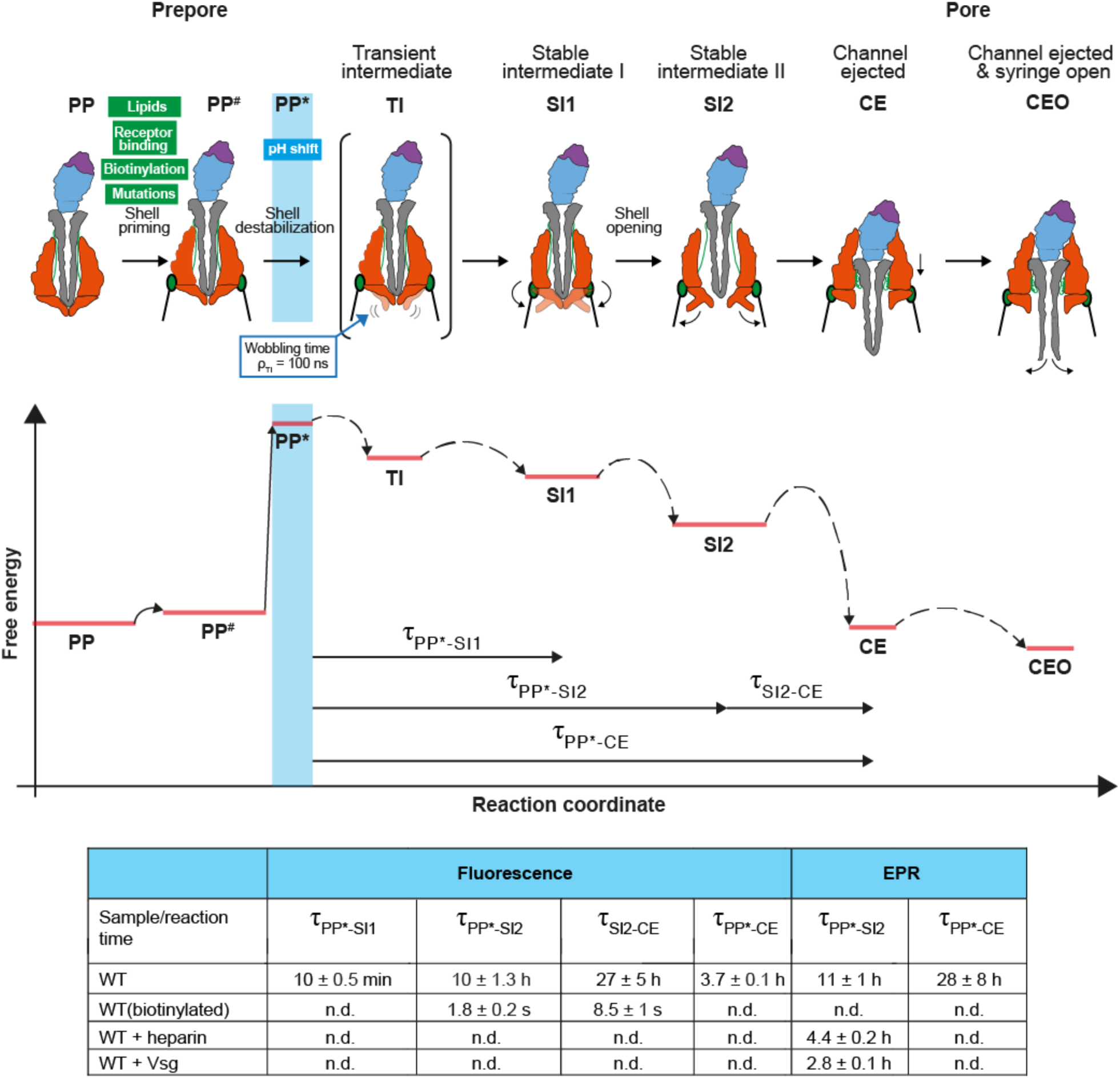
Consecutive model of shell destabilization, opening and channel ejection. Top panel: Scheme of the transition steps. **Middle panel**: A qualitative energy landscape depiction of prepore-to-pore transition and reaction times (τ) of the transition steps which are summarized in the table below the scheme. PP = prepore ground state, PP^#^ = primed prepore state, PP**\*** = primed prepore state after pH shift, TI = transition intermediate with flexibility at the neuraminidase domain, SI1 = stable intermediate 1 with flipped-out RBDs A, B and C, SI2 = stable intermediate 2 with flipped-out RBDs and an open shell, CE = channel ejected pore state with a closed channel-tip, CEO = channel ejected pore state with an open channel-tip. **Bottom panel**: Table showing reaction times of the indicated transition steps. In fluorescence, kinetic fit model 4 (Supplementary Note 1) was used for wt.

In the wild type, shell opening, transition from PP* to the stable intermediate 2 (SI2), has a reaction time of ∼10 h, indicating a low probability of the reaction to be initiated. We have demonstrated by smFRET and EPR that SI2, i.e., the toxin with an open shell but not yet ejected channel, exists in solution but cannot be visualized by cryo-EM. In sum, the transition from the prepore to the pore state (CE, channel ejected state) has a reaction time of ∼28 h *in vitro* with channel ejection being the rate-limiting step. Notably, the precision of the fluorescence measurements was high enough to reveal a small fraction of toxins (15%) with a faster overall reaction time of ∼3.7 h. This kinetic heterogeneity is also visible in single-molecule fluorescence traces monitoring the transitions of biotinylated wildtype molecules immobilized via neutravidin. After a short lag phase, the shells of individual toxins open rather continuously in a broad range of reaction times from 60 ms to 1.6 s, which confirms the presence of fast and slow toxins in this step (Figure 3d). We found a significant (50%) fraction of toxin with long transition times, which indicates limited cooperativity between the protomers (see correlation between times in Supplementary Figure SN5a-c). This kinetic heterogeneity is a hallmark for single protein molecules with many degrees of freedom for functionally relevant transitions^31^. On the contrary, the channel ejection has a long lag phase, but then this transition happens instantaneously in a single step within our time resolution of 60 ms. Interestingly, the channel ejection of biotinylated wildtype is rate limiting (reaction time of 8.5 s) and no intermediate can be detected (Figure 3f). This indicates that channel ejection is a spontaneous reaction, in which all five linkers must compact at the same time. We previously showed that once the channel ejects (CE state), it enters the host membrane where its tip undergoes a conformational change, inducing the opening of the channel (CEO state)^25^.

We have directly demonstrated that the high-pH-shift-induced destabilization and reorganization of the shell is accelerated by the interaction of the toxin with membranes and receptors *in vitro*. This acceleration is likely increased *in vivo* due to the combination of various lipids and receptors and their higher local concentration in the membrane. Interestingly, the accelerating effect of receptors can be mimicked by destabilizing the shell through biotinylation or site-directed mutagenesis of lysines at the protomer-protomer interface. Since the shells of these mutants or biotinylated toxins were closed at neutral pH, the lysines cannot be involved in an electrostatic lock. This is in line with our previous conclusion that TcAs, in general, do not have a classic pH switch^14^. Instead, the lysines are rather involved in a general stabilization through polar and electrostatic interactions either within domains, or between the shell domains of two protomers and their modification destabilizes the shell and accelerates its opening at high pH similar to the interaction with receptors. These mutants could be of use in obtaining better functioning and perhaps more applicable toxin varieties for clinical therapies and/or biological insect pest control.

In contrast, stabilizing modifications of specific protomer-protomer interactions, as in the case of the K1179W mutant, result in the blocked stable intermediate 1 (SI1) that cannot open even after destabilization of the shell at high or low pH and thus cannot intoxicate cells. Antibodies or nanobodies that would mimic this stabilizing mutant and prevent shell opening would be ideal antidotes against Tc toxins of human pathogenic bacteria.

## Acknowledgements

We gratefully thank K. Vogel-Bachmayr, O. Hofnagel and T. Wagner for technical support in the wet lab, EM central facility and statistical analysis, respectively. This work was supported by funds from the Max Planck Society (to S.R.) and the European Research Council under the European Union’s Horizon 2020 Programme (ERC-2019-SyG, grant no. 856118 to S.R), by the Deutsche Forschungsgemeinschaft (DFG, German Research Foundation) under Germany’s Excellence Strategy–EXC 2033–390677874–RESOLV (to E.B.) and by funds from the University of Geneva (to E.B.).

## Author contributions

S.R., E.B., and C.S. conceived and supervised the study. P.N.N., D.R., A.B. and D.P. collected and processed cryo-EM data and built the atomic models. P.N.N., D.R. and S.R. analysed the structures. J.F. and R.K. collected and processed smFRET data. J.F., R.K. and C.S. analysed smFRET data. T.E.A. collected preliminary EPR data. S.K. collected and processed all EPR data. S.K. and E.B. analysed EPR data. S.K., D.R., E.B. and S.R. critically discussed the EPR data. Y.X. and M.D. provided Vsg. O.S. prepared Vsg for EM and EPR. P.N.N., D.R., J.F., and S.K. prepared figures and wrote the supplementary material. P.N.N., E.B., C.S. and S.R. wrote the manuscript, with critical input from all authors.

## Additional Information

Correspondence and requests for materials should be addressed to S.R., E.B. or C.S.

## Data availability

The cryo-EM densities of TcA(1279-At647N), TcA(K1179W) and TcA(K567W/K2008W) have been deposited to the Electron Microscopy Data Bank (EMDB) under accession codes: EMD-16793, EMD-16791 and EMD-16792 respectively. These depositions include sharpened maps, unfiltered half-maps and the refinement masks. The atomic coordinates of the protein structures have been submitted to the Protein Data Bank (PDB) under accession codes: 8CQ2, 8CPZ and 8CQ0 respectively. We used the following previously published structure for modelling and comparisons: PDB ID: 6RW6. Raw data generated during the current study are available from the corresponding authors on request. Source data are provided with this paper.

## Methods

### Design of TcA variants

*P. luminescens* TcdA1 (TcA) WT contains ten cysteines per protomer, three of them being surface-exposed (C16, C20, C870) and thus expected to interfere with site-specific labelling for Electron Paramagnetic Resonance (EPR) spectroscopy and single-molecule Förster Resonance Energy Transfer (smFRET) spectroscopy experiments. In this regard, we mutated the three cysteines to serines via QuikChange-PCR. The labelling of this variant with 2-fold nitroxide spin labels under conditions described below did not result in a detectable CW EPR signal, indicating that the remaining cysteines are inaccessible. The obtained variant, TcA(C16S/C20S/C870S), therefore served as the basis for all TcA variants subsequently designed for labelling with iodacetamido-PROXYL (IAP), Bodipy-FL-iodacetamide (BFL), and Atto647-N-iodacetamide (At647N).

We next introduced a cysteine at position 1193 at the bottom of the shell to follow shell opening of TcA (Supplementary Figure 1a), or two cysteines at positions 914 and 2365 (upper part of the shell and funnel, respectively) to follow channel insertion (Supplementary Figure 1b), resulting in TcA(1193Cys) and TcA(914Cys/2365Cys). These TcA variants served for subsequent labelling with IAP for EPR and BFL for single-molecule FRET.

To label TcA with BFL and Atto647N for single-molecule FRET and fluorescence polarization experiments, we created TcA(1279Cys) and TcA(914Cys) analogously. In addition, we introduced a cysteine at position 1041 of *P. luminescens* TcdB2-TccC3 (TcB-TcC), resulting in TcB-TcC(1041Cys) to attach Atto647N.

To mimic biotinylation of TcA at different surface-exposed lysines, we first assessed the protein for all surface-exposed lysines, particularly focusing on those near protomer-protomer interfaces. For this, we used the H++ server (newbiophysics.cs.vt.edu/H++)^32^ to predict the electrostatic landscape of TcA at pH 11. Both the WT model (PDB ID: 6RW6) and models where a chosen lysine was changed to a tryptophan were used for the prediction. Since biotinylating the exposed lysines would neutralize them, we mutated the chosen lysines via QuikChange-PCR to tryptophans. Tryptophan was chosen because it comes closest in size to biotin. Of those observed lysines, three were selected that are near protomer-protomer interfaces; K567 at a lysine cluster on the shell, K1179 on the neuraminidase-like domain near the tip of the channel and K2008 on the linker (Supplementary Figure 12a,b). Mutation resulted in the TcA(K567W), TcA(K1179W) and TcA(K2008W) variants used for negative stain EM and cryo-EM, as well as cell cytotoxicity. Furthermore, a double mutant with K567W and K2008W, TcA(K567W/K2008W), was further mutated to add 1193Cys, or 914Cys/2365Cys for EPR measurements. Similarly, TcA(K1179W) was mutated to add either 1193Cys for EPR and single-molecule FRET measurements, or 2365Cys for single-molecule FRET measurements.

For single particle cryo-EM analyses of prepore-to-pore transition intermediates, some key TcA variants were selected, namely: TcA(1279-At647N) which served as a WT/control for cryo-EM experiments as it showed WT-level prepore-to-pore transition kinetics in single-molecule experiments, TcA(K1179W) which showed almost no transition to the pore state in EM, EPR and smFRET experiments and ABC(K567W/K2008W) which showed a significantly increased rate of prepore-to-pore transition in the mentioned experiments. The TcA variants TcA(K1179W) and TcA(K567W/K2008W) in ABC(K567W/K2008W) were both based on the unmodified TcA WT and therefore lacked the modifications made for variants designed for labelling. All three datasets were collected on proteins incubated at pH 11.2 as described in the subsequent sections.

### Protein production

TcA (WT and all variants) and TcB-TcC (WT and TcB-TcC(1041Cys)) were expressed in BL21-CodonPlus(DE3)-RIPL in 10 L LB medium and purified as described previously^24^. All variants of TcA were expressed and purified like TcA WT with the exception that 0.5 mM TCEP was present in all purification buffers to keep all surface-exposed cysteines in reduced state. The extracellular, highly O-glycosylated domain of Vsg was produced and purified as previously described^18^, with an additional desalting step into 20 mM HEPES pH 7.5, 150 mM NaCl, 0.05% Tween-20 using a PD Spin-Trap G-25 desalting column (Cytiva).

### Spin-labelling of TcA for EPR studies and determination of labelling efficiency

Immediately before labelling, TcA variants were buffer exchanged to labelling buffer (10 mM Hepes-NaOH pH 7.5, 150 mM NaCl, 0.05% Tween-20) and concentrated to at least 5 µM (protomer concentration). We used IAP (Sigma-Aldrich, cat. no. 253421) for labelling because it sustains highly basic pH values necessary for this study without the drawbacks of commonly used *S*-(1-oxyl-2,2,5,5-tetramethyl-2,5-dihydro-1H-pyrrol-3-yl)methyl methanesulfonothioate (MTSL)^33^ or maleimide-functionalized labels^34^. For TcA(1193Cys), a two-fold molar amount of IAP per protomer was added, and for TcA(914Cys/2365Cys), a three-fold molar amount was added. Labeling was performed for 1 h at 22 °C, followed by overnight incubation at 4 °C. Subsequently unbound IAP was removed by diluting the samples three times 10-fold in labelling buffer and concentrating back to the original volume with an Amicon Ultra 100 kDa cutoff concentrator (Millipore). The protein concentrations for subsequent EPR experiments are given in Supplementary Table 1.

For determination of labelling efficiency, 20 µL of the concentrated samples were pipetted into glass capillaries with 0.7 mm inner diameter (Blaubrand). Next, continuous wave (CW) EPR spectra were either recorded on a Miniscope MS 5000 (Magnettech, Bruker Biospin) at X band (9.5 GHz), or on a Bruker X-band (9.8 GHz) E580 (Bruker Biospin) spectrometer equipped with Bruker Super high Q resonator at ambient temperature (21 °C). Measurement conditions for Miniscope (Supplementary Figure 1d) were: 10 mW power, 0.15 mT modulation amplitude, 100 kHz modulation frequency, 15 mT sweep width, 80 s sweep time. Measurement conditions for spectra detected with the Bruker spectrometer (red spectrum in Supplementary Figure 1d, 13) were the same as given below for the kinetic measurements.

Labelling efficiency was calculated as a percent ratio between the spin concentration and the concentration of introduced cysteines (1193Cys or 914Cys/2365Cys) in TcA protein (protein concentration determined by the absorption at 280 nm with extinction coefficient 400,750 M^-1^ cm^-1^ per protomer). Spin concentration was determined from double integration of X-band CW EPR spectra (Supplementary Figure 1d, 13) by comparison to a standard stock solution of 100 µM of 4-Hydroxy-2,2,6,6-tetramethylpiperidine 1-oxyl (TEMPOL, Sigma Aldrich) in water measured at the same conditions. The error in spin labelling efficiency taking into account uncertainties in protein concentration and baseline-noise-related errors in the double integral value is estimated to be ±10%^35^. All labelling efficiencies, as well as corresponding protein and spin concentrations are listed in Supplementary Table 1.

### Labelling of TcA and TcB-TcC for fluorescence polarization and FRET studies

TcA(1193Cys), TcA(1279Cys), and TcA(914Cys) were labelled with the green fluorescent dye BFL or the red fluorescent dye Atto647N for single-molecule FRET or fluorescence polarization experiments. For experiments that require one fluorophore per TcA protomer, a two-fold molar excess of label over the protomer concentration was added in labelling buffer (10 mM Hepes-NaOH pH 7.5, 150 mM NaCl, 0.05% Tween-20), followed by 1 h incubation at 22 °C and subsequently overnight at 4 °C. For experiments that require one fluorophore per pentamer, a 1.1-fold molar excess of label over the pentamer concentration was added and incubated as above. TcB-TcC(1041Cys) was labelled with a two-fold molar excess of Atto647N using the same conditions. After labelling, unreacted dye was removed using a Superose 6 increase column equilibrated in 10 mM Hepes-NaOH pH 7.5, 150 mM NaCl.

### Biotinylation of TcA variants

To immobilize the proteins for TIRF measurements, TcA(914Cys) and TcA(1279Cys) were biotinylated using EZ-link NHS-PEG12-Biotin (ThermoFisher, cat. no. A35389). Biotinylation was performed in labelling buffer simultaneous to the labelling reaction with BFL and Atto647N, respectively. 5.6 µM NHS-PEG12-Biotin were added to 2.8 µM TcA pentamer from a 500 µM stock solution in water-free DMSO, and the reactions were incubated for 20 min at 22 °C, followed by overnight incubation at 4 °C. Unreacted biotin was removed analogously to the removal of free IAP label.

### Simulation of spin label distances

The expected interspin distances between the chosen sites were simulated by rotamer library analysis with MMM 2018.2^36,37^ in Matlab R2017b (Mathworks). To speed up the rotamer library calculations, the amino acids of TcA located at >10 nm distance from the labelling sites were removed from the structure prior to the simulations. The implemented room temperature rotamer library for IAP was used to calculate the allowed rotamers attached at cysteine-substituted sites. TcA(1193-IAP) revealed 22 allowed rotamers per chain in the prepore state, which increased to 40-62 depending on the chain in the pore state. TcA(914-IAP/2365-IAP) showed 35/91 rotamers (prepore) and 29-34/72-77 (pore) for IAP at position 914 and 2365, respectively. For representation, single rotamers at each site are depicted in Supplementary Figure 1a,b.

The remaining native, not surface exposed cysteines at TcA positions 499, 756, 870, 1149, 1236, 1372, 2273 and 2445 revealed only 1-2 allowed IAP rotamers, which indeed was proven experimentally (no detectable CW EPR signal after labelling).

### Continuous wave (CW) EPR kinetic measurements

10 µL of labelled TcA in labelling buffer was mixed with 10 µL of measurement buffers (pH 4.0: 100 mM Na-acetate, 150 mM NaCl, 0.1% Tween-20; pH 7.5: labelling buffer; pH 11.2: 100 mM CAPS-NaOH, 150 mM NaCl, 0.1 % Tween-20), thereby resulting in two times lower final protein and spin concentration as compared to values given in Supplementary Table 1. CW EPR kinetic measurements were performed on a Bruker X-band (9.8 GHz) E580 (Bruker Biospin) spectrometer equipped with Bruker Super high Q resonator at ambient temperature. Kinetics were plotted as peak-to-peak intensity of the central spectral line over time, normalized to the intensity of the first spectrum (3 min dead time). Spectra were recorded with 0.75-2 mW power, 0.1-0.15 mT modulation amplitude, 100 kHz modulation frequency, 13 mT sweep width, 40 s sweep time, 4 or 8 averages per scan. CW EPR kinetics were successfully detected on TcA(1193-IAP), which showed increased dynamics of the shell over time within the sensitivity range of X-band EPR (30 ps to 30 ns). The CW EPR kinetics were fitted with a mono-exponential function. No changes in the dynamics of the spin labels during the transition from prepore to pore were observed for TcA(914-IAP/2365-IAP). The corresponding time constants of the fits (kinetics at different pH values and in presence of liposomes) are summarized in Supplementary Table 2.

### Double Electron Electron Resonance (DEER)

10 µL of the labelled TcA variants in labelling buffer were mixed with 10 µL of the different measurement buffers (see above) prepared with 50% v/v D_2_O and incubated for time intervals between 0 min and 72 h at 21 °C or at 4 °C as stated in the corresponding legends. After incubation, 20 µL of fully deuterated glycerol (glycerol-d8, Sigma-Aldrich) were added, thereby resulting in four times lower final protein and spin concentration as compared to values given in Supplementary Table 1. Finally, the samples were transferred into 3 mm quartz capillaries (Aachener Quarzglas Heinrich) and shock-frozen in liquid N_2_. The overall mixing/freezing time took 2 - 6 min. The final spin concentration in all DEER tubes did not exceed 8 µM, which minimizes the contribution of inter-molecular background and increases the fidelity of the extracted distances. For all pulsed EPR experiments a Bruker Q-band E580 spectrometer equipped with 150 W TWT amplifier was used together with a Bruker SpinJet-AWG for shaping the microwave pulses and a home-made microwave cavity suitable for 3 mm tubes^38,39^. The dead-time free 4-pulse DEER sequence^40^ with 16-step phase cycling^41^ and Gaussian pulses^42^ was used. Additionally, 8 steps of 16 ns cycling were used for ESEEM modulation averaging. Shot repetition time (SRT) was 3 - 4 ms. The interpulse delay d1 was 400 ns, d2 varied in µs range between the traces, 8 ns stepping Δt was used. All Gaussian pulses were optimized via nutation experiments and set to the same length of 32 ns (∼13.6 ns full width at half maximum) to achieve the same excitation bandwidth. The pump pulse was applied at the position of maximal microwave absorption of nitroxides. The pump pulse was set with + (90 - 100) MHz offset from the observer frequency, with both frequencies chosen symmetrically in the microwave dip. All pulsed measurements were performed on frozen samples at 50 K. The total acquisition time varied between the samples in the range 1 - 72 h (short - long traces, respectively).

### DEER data analysis and validation

DEER data were analyzed using Tikhonov regularization or NeuralNet tool^43^ with the generic set in DeerAnalysis2019^44^ according to the guidelines^45^. The zero time was determined experimentally and set to 120 ns. For Tikhonov regularization, the homogeneous background function with D = 3 was chosen for the data shown in Supplementary Figure 2a to correct for the minor background contribution. For TcA(1193-IAP), both Tikhonov regularization and neural network analysis was used for validation of DEER data in Supplementary Figure 2c, showing no significant discrepancies between the methods. Therefore, the supplementary DEER data in Supplementary Figure 3e, 5b,e, and 13f were analysed and validated using neural networks only.

The accuracy of the measured short distances was further validated by testing the absence of ghost frequencies in the multispin systems of TcA(1193-IAP) and TcA(914-IAP/2365-IAP) by reducing pump pulse with a power scaling factor λ = 0.5 (Supplementary Figure 2d, 5c)^46^.

### DEER “snapshot” kinetics

To obtain time-resolved DEER kinetics, samples were incubated for different times, snap-frozen in liquid nitrogen and measured. To resolve the small changes in modulation depths, short DEER traces were recorded to reach higher signal-to-noise ratios. The collected sets of primary DEER traces (Supplementary Figure 2b, 5d) revealed a clear trend of decreasing modulation depths for TcA(1193-IAP) due to the disappearance of the short distances and increasing modulation depths for TcA(914-IAP/2365-IAP) due to the appearance of the short distances upon prolonged incubation at pH 11.2. Thus, to estimate the time constant in both cases, the modulations at the first inflection point were plotted against incubation time. The modulation depth at this point should mainly represent the fraction of Tc proteins in the ensemble characterized by short distances. For the determination of the inflection point, the trace with the most pronounced high frequency oscillation and best signal-to-noise ratio was chosen and fitted with a polynomial function in DEER Analysis2019. The 1^st^ and 2^nd^ derivatives of the obtained polynomial fit were calculated to determine the 1^st^ inflection point in the time domain. The modulations extracted at this point after different incubation times were normalized to the maximal value for the two batches of TcA(914-IAP/2365-IAP).

### Confocal single-molecule fluorescence spectroscopy

Single-molecule measurements with Multiparameter Fluorescence Detection (MFD) were performed on a home built setup based on an Olympus IX70 inverted microscope as described by Widengren *et al*., 2006^47^. To prevent concentration drops due to unspecific adhesion of the molecule, the cover glass was passivated with BSA using 10µM stock solution, which was washed off after 2 minutes incubation time. An Olympus UPlanSAp 60x/1.2 objective was used. A linearly polarized, pulsed diode laser with a wavelength of 495 nm (LDH-D-C 495, PicoQuant) operated at 64 MHz was used. In case of PIE configuration^48^, an additional red laser with a wavelength of 635 nm was used, both operated at 32 MHz in an alternating order. For detection, the beam is split into parallel and perpendicular polarization and filtered by colour using ET 535/50 and HQ 730/140 (AHF, Analysentechnik) bandpass filters and finally detected by 8 detectors. Single photon counting was done with synchronized channels (HydraHarp 400, PicoQuant, Germany) operating in Time-Tagged Time-Resolved (TTTR) mode. Burst selection and data analysis were done using established procedures and in house software described in^49^, available upon request on the homepage of the Seidel group (https://www.mpc.hhu.de/software.html).

Single-molecule events were identified using a burst search algorithm according to Fries *et al*. (1998)^50^ using a Lee filter, a threshold of 0.2 ms and a minimum of 60 photons per burst. For anisotropy PDA, the whole trace including all bursts were analysed. For heteroFRET analysis, molecules with a stoichiometry between *S*=0.5 to *S*=0.8 were selected. The labelling geometry of 5 possible labelling spots for TcA(914-BFL)-Biotin:TcB(1041-At647N)-TcC and labelling scheme of 5:1 donor to acceptor led to a higher than usual value of the stoichiometry, where multiple donors were attached to the molecule with only one acceptor. However, only one of the label positions had a distance to the acceptor where FRET occurs, giving the possibility to qualitatively measure FRET efficiency derived distances.

In heteroFRET studies, static FRET lines as described by Kalinin *et al*. (2010)^51^ were used for consistency and quality control. A static FRET line relates the lifetime of the donor in the presence of an acceptor, 〈*τ_D_*_(*A*)_〉*_F_*, to the intensity-based FRET efficiency, *E*. In case of the Tc toxin, no dynamic exchange between conformational states was expected. This matches the experimental data, where the double labelled FRET population of the Tc toxin in prepore and pore states on the 〈*τ_D_*_(*A*)_〉*_F_* − *E* is located on the static FRET line (see Supplementary Figure 11).

Polarization-resolved Fluorescence correlation spectroscopy (pFCS)^30^ was performed by computing the auto-cross-correlating the two corresponding parallel and perpendicular polarized time traces of the detected photons using in house software^52^. Amplitudes and relaxation times of the observed bunching terms (translational diffusion, triplet and reaction kinetics) were extracted by fitting appropriate model functions to the data applying the same software. For details, see Supplementary Note 2

### Single-molecule TIRF microscopy

Imaging was performed on a home built Total Internal Reflection Fluorescence (TIRF) setup based on Olympus IX70 microscope and CCD camera (emCCD, DU-897D-BV, Andor). Using a special TIRF objective (Apo N 60x, 1.49 NA, Olympus) with very high numerical aperture leads to total internal reflection of the light beam on the surface of the glass. For homoFRET-studies a single colour excitation with a 488 nm continuous wave laser (Cobolt MLD) was used. In case of heteroFRET an additional 635nm cw-laser (Cobolt MLD) was added using alternating excitation (ALEX), where excitation was alternating, triggered with a home-built switch. Emission light passes a dichroic mirror and is then split by polarization (homoFRET) or by colours (heteroFRET) and projected as two spatially separated images (parallel/perpendicular or green/red, respectively) on the camera using an image splitter (OptoSplit II, Cairn Research Ltd). Splitting by colours was done using HC BS 580 Imaging (AHF Analysentechnik) and bandpass filters (green: HQ 535/50, red: HQ 680/60) and splitting by polarization was via polarizing beam-splitting cube. For every measurement, 2048 images with a pixel size of 512×512 pixels were taken with a single frame time of 29.44 ms. Spot selection and data analysis was done using the software iSMS^53^. For background subtraction, mean intensity of a background mask was used. Spot selection was done using an automatized intensity-based algorithm. To calibrate the overlay of the two individual images, a measurement with inhomogeneous areas was used. Polarization traces were smoothed using the Savitzky-Golay algorithm with varying number of points.

#### HeteroFRET assay for channel ejection

FRET Efficiency traces were averaged based on the time interval before and after the transition. The correction factors for the FRET efficiency were calculated based on the spectral properties of the dyes and applied globally for each molecule. The following correction factors were used:

| Correction factors (see Supplementary Methods) | HeteroFRET<br>Alexa488/Atto647N | homoFRET<br>Atto647N/Atto647N |
| --- | --- | --- |
| $\alpha$ | 0.016 | 0 |
| $\delta$ | 0.02 | 0 |
| $\frac{g_{R\vert A}}{g_{G\vert D}}$ | 1.25 | 1 |
| $\frac{eff \Phi_{F,A}}{eff \Phi_{F,D}}$ | $\frac{0.65}{0.8} = 0.81$ | 1 |
| $\gamma$ | 1.02 | 1 |

For heteroFRET efficiency, background corrected fluorescence *F* was used following equations described in section intensity-based MFD. Due the long necessary observation time of up to 40 seconds, most FRET pairs bleached during pre-transition time. Thus, could resolve only 13 single reactions with FRET efficiency change before photobleaching.

#### HomoFRET anisotropy assay for shell opening

Here, one has to take into account that the high numerical aperture of the TIRF objective influences the polarization of the laser beam, leading to a mixture of polarizations. Additionally, the laser beam enters the solution at an angle greater than a critical angle for total reflection. This effect was studied in detail and a shift to lower anisotropy values was observed^54,55^. Additionally, due to a high amount of different labelling sites of the biotin on the Tc toxin, the initial orientation and therefore polarization values of the toxin were distributed. However, in this assay the difference of polarization of the signal before and after the transition from prepore to pore was of interest. Therefore, we applied a polarization offset based on the mean polarization value before the pH change 〈*P*(*pH*7)〉 to every trace shifting the polarization value before the pH change to *P*=0. The polarization value resulted in 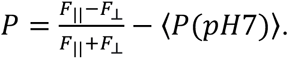.

In total, 765 homoFRET pairs were recorded where 48 of them showed a trackable transition, while the rest were either FRET inactive due to the labelling scheme described in the Supplementary Table 4 or the dyes were bleached. The time after pH change (t = 0) until the signal changes before the transition starts is defined as the pre-transition time (see Figure 3b). The transition time is defined as the time the signal needs until it is permanently changed. To determine these time points, a binned derivative of the trace was calculated as the change of between two signals over the interval, which was binned afterwards with a binning size of two frames.

### Immobilization of toxins for single-molecule TIRF measurements

Tc toxins were immobilized on the surface using biotinylated BSA. Cleaning of nunc chambers (Nunc Lab-Tek II, thickness No. 1.5H, ThermoFisher) was done by activating them in an oxygen plasma for 2 min (FEMTO Plasma Cleaner, Diener electronic). After cleaning, the surface was incubated for 10 min with biotinylated BSA (Sigma-Aldrich, 3 mg/ml in PBS), washed several times with PBS, incubated for 10 min with neutravidin (Invitrogen, 20µg/ml in water) and washed again with buffer. The biotinylated toxins were then added and incubated for up to 30 min. Finally, the chamber was washed again to remove diffusing toxins and dyes.

### Kinetics of shell opening and channel ejection monitored by single-molecule confocal fluorescence anisotropy measurements

Long confocal fluorescence measurement series were performed with several 10 h measurements, which were cut into 2h-intervals for data analysis. Ratio of parallel and perpendicular signal S_||_ and S_⊥_ of each burst of the whole trace was fitted in a Time Window (TW) based, anisotropy Photon Distribution Analysis (aPDA)^52^. More details are given in Supplementary Methods. The applied model fitted global values for the ratios/anisotropy with varying fractions. Typical data are shown in Supplementary Figure 7. Time dependent fractions were fitted by comparing four distinct kinetic models. See Supplementary Note 1 for all details.

### Simulation of inter-fluorophore distances by AV-Simulations

In order to design a label scheme with dyes positioned in the FRET sensitive range to measure the conformational change due to shell opening and pore ejection, a FRET position screening was applied following Dimura et al.^56^ and using in house software to find the most promising labelling positions for specified structural models. In short, it simulates for each position so called sterically Accessible Volumes (AVs)^57^ to a series of positions on the biomolecule while calculating the expected FRET-averaged inter-dye distance 〈*R_DA_*〉*_E_* (typical examples are shown in Supplementary Figure 8. The labelling positions were selected in two steps: (1) To reduce time and effort, the position for the donor dye was set to an already used position for other experiments and AV simulations were used to suggest positions for the acceptor in a FRET-sensitive range. (2) Suitable positions were checked for the presence of quenching amino acids in the surroundings^58^. Finally, the protein was labelled and tested using smMFD. The outcome was that the most sufficient label scheme in terms of good label ratio for shell opening was TcA(1279-At647N) and for pore ejection, TcA(914-BFL:TcB(1041-At647N)-TcC).

The relevant inter-dye distances and the expected FRET efficiencies of the studied Toxin constructs are compiled in Supplementary Table 4.

### Cell intoxication

HEK293T cells (ThermoFisher) were intoxicated with pre-formed holotoxin composed of TcB-TcC WT (unmodified) and TcA WT (unmodified), TcA(1193-IAP) or TcA(914-IAP/2365-IAP). 5 × 10^4^ cells in 400 μL DMEM/F12 medium (Pan Biotech) were grown overnight and subsequently intoxicated with 0.2, 0.5 or 5.0 nM of toxin. Incubation was performed for 16 h at 37°C before imaging. Experiments were performed in triplicate. Cells were not tested for Mycoplasma contamination.

To test the toxicity of TcA biotinylation mimics (K567W, K1179W and K2008W), HeLa cells were intoxicated with pre-formed holotoxins of each TcA mutant and TcB-TcC WT. 6 × 10^4^ cells in 1 mL DMEM medium (Pan Biotech) were grown for 48 h before addition of 5.8 nM of toxin. The cells were then incubated at 37°C with images acquired at 6, 7 and 8 h. Subsequently, the number of rounded cells was calculated for each mutant using ImageJ/Fiji^59^ as a quantification of cytotoxicity. Experiments were performed in triplicate. Cells were not tested for Mycoplasma contamination. Quantified data was analysed using an unpaired two-tailed t-test.

### Negative stain electron microscopy kinetics

Prior to pH-induced prepore-to-pore transition, TcA (WT, IAP-labelled variants and biotin mimics) was mixed with a 2-fold molar excess of TcB-TcC to form holotoxins (ABC WT, ABC(1193-IAP), ABC(914-IAP/2365-IAP), ABC(K567W), ABC(K1179W), ABC(K2008W) and ABC(K567W/K2008W)). The resulting ABC holotoxins were then separated from excess TcB-TcC by size exclusion chromatography on a Superose 6 increase 5/150 column (GE Life Sciences) equilibrated in labelling buffer. The prepore-to-pore formation of ABC was then started by mixing 0.1 - 0.2 mg/ml protein with 100 mM CAPS-NaOH pH 11.2, 150 mM NaCl, 0.1% Tween-20, or 100 mM sodium acetate pH 4.0, 150 mM NaCl, 0.1% Tween-20 in a 1:1 ratio. After the indicated time intervals of incubation (1 min – 72 h), the proteins were applied to glow-discharged copper grids with 8 nm amorphous carbon layer pre-treated with/without 0.1% polylysine, stained with 0.75% uranyl formate and imaged using a Tecnai Spirit transmission electron microscope (ThermoFisher) operated at 120 kV and equipped with a TVIPS TemCam F416 detector. Images were recorded at a pixel size of either 1.67 Å/pixel or 2.61 Å/pixel.

For every variant and incubation time, 36–64 micrographs were recorded to obtain at least 4000 particle images. To quantify prepores and pores, we trained two models in crYOLO^60^ to pick specifically ABC prepores and pores from the EM micrographs, respectively. Subsequently prepore and pore particles were picked using the prepore and pore models in crYOLO (Supplementary Figure 5,6,12) and the percentage of pore state as a function of time was used to determine the rate of prepore-to-pore transition. In addition, we picked prepores and pores together using a general picking model in crYOLO, and subjected the picked particles to 2D classification using ISAC^61^ in SPHIRE^62^. The numbers of particles in prepore and pore class averages were then quantified. To validate this quantification, we performed a Bayesian binomial regression following 2D classification to calculate the probability of obtaining pores using the mutant datasets as an example. The results mirrored the initial analysis (Figure 4a, Supplementary Figure 12d).

To test the influence of receptors on prepore-to-pore transition kinetics; 350.9 nM of either ABC WT or TcA WT (protomer concentration of TcA) was first mixed with its receptors. Then, the toxin-glycan mix was preincubated for 1 h at 21 °C in pH 7.5 buffer, before switching the pH to either pH 11.2, or pH 4 and incubating for a further 2 h to induce prepore-to-pore transition. For this, 16.5x (5.8 µM) molar excess of the glycan receptors heparin and Lewis X (conjugated to bovine serum albumin (BSA)), as well as 70 nM of Vsg (1:1 with the pentamer concentration of TcA) were used. Heparin, made from porcine intestinal mucosa was procured from Merck (Product number: 375054). BSA-Lewis X was prepared as previously described^12^.

Subsequently, the samples were negative stained, imaged and quantified as previously described. For prepore-to-pore transition with heparin at pH 4, performing the experiment in buffer led to a severe aggregation of the protein (Supplementary Figure 5b), presumably due to its strong negative charge, making quantification difficult. Therefore, to circumvent this, the experiment was performed directly on the negative stain grids. In this regard, the grids were first pre-incubated with 7 nM of ABC WT (holotoxin) for 1 min before blotting and addition of the glycan. After a 30 min incubation of the grids with heparin in humid conditions, the buffer was exchanged to pH 4 and then they were incubated for another 2 h before blotting, staining and imaging. Quantified data was analysed using an unpaired two-tailed t-test.

### Cryo-EM of TcA(1279-At647N) at pH 11.2

TcA(1279Cys) contained all the mutations introduced in variants that were designed for labelling as previously described, namely: C16S, C20S, C870S and T1279C. 0.25 mg/ml TcA(1279Cys), labelled with the fluorophore At647N in 10 mM HEPES-NaOH pH 7.4, 150 mM NaCl, 0.02% Tween-20 to form TcA(1279-At647N), was mixed 1:1 with 100 mM CAPS-NaOH, 150 mM NaCl, 0.05% Tween-20 and incubated for 6 h at 22 °C. Subsequently, 4 µl were loaded on a freshly glow discharged Quantifoil 2/1 300 mesh gold grid with 2 nm amorphous carbon support and incubated for 5 min on the grid. The sample was then vitrified in liquid ethane using a Vitrobot Mark IV (ThermoFisher) set to a blotting time of 3.0 s, blot force of −3, 100% humidity and 12 °C.

A cryo-EM data set was collected with a Cs-corrected Titan Krios transmission electron microscope (ThermoFisher) equipped with a X-FEG and a K3 direct electron detector with Bioquantum energy filter (Gatan) and a slit width of 15 eV. Images were recorded using the automated acquisition program EPU (ThermoFisher) in a super-resolution mode at a magnification of 81,000, corresponding to a pixel size of 0.44 Å/pixel on the specimen level. 13,059 movie-mode images were acquired in a defocus range of −1.2 to −3.0 μm. Each movie comprised 60 frames acquired over a total integration time of 3.0 s with a total cumulative dose of ∼ 78 e^-^/Å^2^.

### Cryo-EM image processing of TcA(1279-At647N)

Movie frames were aligned, dose-corrected, binned twice to 0.88 Å/pixel and averaged using MotionCor2^63^, implemented in TranSPHIRE^64^. CTF parameters were estimated using CTFFIND4^65^. 573,814 particles were auto-picked using crYOLO^60^ with a general particle model and extracted with a box size of 384 pixels.

Reference-free 2D classification of the dataset was performed with ISAC^61^ in SPHIRE^62^ with a pixel size of 4.40 Å/pixel on the particle level. The ‘Beautifier’ tool of SPHIRE^62^ was then applied to obtain refined and sharpened 2D class averages at the original pixel size, showing high-resolution features (Supplementary Figure 15a,b). 2D classification resulted in 80,710 particle images corresponding to the prepore and 150,465 particle images corresponding to the pore state of TcA. 3D-refinement of the 80,710 prepore particle images was performed with MERIDIEN in SPHIRE^62^ in C5 symmetry using the density map of TcdA1 WT (EMDB ID 10033), low-pass filtered to 20 Å, as initial reference. After initial 3D refinement, the particles were centred and re-extracted using the rotation/translation parameters obtained in MERIDIEN and 3D refinement was repeated using a mask that included the well-resolved part of the reconstruction, resulting in an average resolution of 3.8 Å according to FSC 0.143. Then, an additional CTF refinement, followed by 3D refinement in Relion 3.1^66^ resulted in a reconstruction with a final average resolution of 3.64 Å according to FSC 0.143. This reconstruction was later used for model building.

The obtained reconstruction lacked density for the receptor-binding domains B and C and 3D variance analysis revealed an additional slightly elongated density at the lower part of the shell. Therefore, to improve the density corresponding to the receptor-binding domains B and C, we applied 3D classification using Sort3D_DEPTH in SPHIRE with a mask focusing on the bottom of TcA (Supplementary Figure 15f). We obtained three 3D classes with 21,332, 20,992 and 16,447 particles, respectively. The largest 3D class showed an elongated additional density below RBD D, next to the neuraminidase-like domain. Local refinement of this 3D class with 1.875° initial angular sampling and subsequent post-processing with the PostRefiner tool improved this extra density, which became visible at low binarization threshold of the resulting map of 4.2 Å according to FSC 0.143 (Supplementary Figure 15f).

### Cryo-EM of TcA(K1179W) at pH 11.2

This variant was based on the unmodified TcA WT and therefore only included the K1179W mutation. 0.9 mg/ml of TcA(K1179W) in labelling buffer was mixed 1:1 with 100 mM CAPS-NaOH, 150 mM NaCl, 0.1% Tween-20 and incubated for 24 h at 22 °C. Subsequently, 4 µl were loaded on a freshly glow discharged Quantifoil 2/1 200 mesh gold grid with 2 nm amorphous carbon support and incubated for 1 min on the grid before blotting. A second round of sample application, incubation on the grid and blotting was performed and then the sample was vitrified in liquid ethane using a Vitrobot Mark IV (ThermoFisher) set to a blotting time of 3.0 s, blot force of 1, 0.5 s waiting time for round 1 and no waiting time for round 2, 100% humidity and 12 °C. A cryo-EM data set was collected with a Talos Arctica transmission electron microscope (ThermoFisher) equipped with a X-FEG and a Falcon III direct electron detector operated in linear mode. Images were recorded using the automated acquisition program EPU (ThermoFisher) at a magnification of 120,000, corresponding to a pixel size of 1.21 Å/pixel on the specimen level. 5494 movie-mode images were acquired in a defocus range of −1.5 to −2.75 μm. Each movie comprised 40 frames acquired over a total integration time of 3.0 s with a total cumulative dose of ∼ 56 e^-^/Å^2^.

### Cryo-EM image processing of TcA(K1179W)

Movie frames were aligned, dose-corrected and averaged using MotionCor2^63^, while CTF parameters were estimated using CTFFIND4^60^ both implemented in TranSPHIRE^64^. 390,444 particles were auto-picked using crYOLO^60^ with a general particle model and of those 356,439 were subsequently extracted using a box size of 352 pixels.

Reference-free 2D classification of the dataset was performed with ISAC^61^ in SPHIRE^62^ with a pixel size of 6.1 Å/pixel on the particle level. The ‘Beautifier’ tool of SPHIRE was then applied to obtain refined and sharpened 2D class averages at the original pixel size, showing high-resolution features (Supplementary Figure 14b). 2D classification resulted in 230,785 particle images corresponding to the prepore and 1921 particle images corresponding to the pore state of TcA.

3D-refinement of the prepore particle images was first performed with MERIDIEN in SPHIRE^62^ in C5 symmetry using the density map of TcdA1 WT (EMDB ID 10033), low-pass filtered to 20 Å, as initial reference. As previously observed, the resulting density map showed a prepore-to-pore intermediate state lacking densities for RBDs B and C. Lowering the binarization threshold revealed an additional density below RBD D, next to the neuraminidase-like domain, also observed previously (Supplementary Figure 14f). Further processing was performed in Relion 3.1^66^ including 3D refinement, particle polishing, CTF and local refinements. Subsequently, postprocessing of the final reconstruction using a mask that includes the well-resolved part of the previous reconstruction resulted in a final average resolution of 2.9 Å according to FSC 0.143, which was used for model building. Then, a focused 3D classification performed as previously described resulted in three classes of 160,851, 21,440 and 48,494 particles respectively. 3D refinement of the largest class produced a 3.3 Å map according to FSC 0.143 that shows the same elongated extra density below RBD D at low binarization threshold (Supplementary Figure 14f).

### Cryo-EM of ABC(K567W/K2008W) at pH 11.2

This variant was also based on the unmodified TcA WT and therefore only included the K567W and K2008W mutations. After holotoxin formation as previously described, 0.5 mg/ml of ABC(K567W/K2008W) in labelling buffer was mixed 1:1 with 100 mM CAPS-NaOH, 150 mM NaCl, 0.1% Tween-20 and incubated for 2 h at 22 °C. Subsequently, the buffer was neutralized by mixing 1:1 with labelling buffer before 4 µl were loaded on a freshly glow discharged Quantifoil 2/1 200 mesh gold grid with 2 nm amorphous carbon support and incubated for 5 min on the grid. The sample was then vitrified in liquid ethane using a Vitrobot Mark IV (ThermoFisher) set to a blotting time of 3.0 s, blot force of 2, 100% humidity and 12 °C.

Two cryo-EM data sets were collected with a Cs-corrected Titan Krios transmission electron microscope (ThermoFisher) equipped with a X-FEG and a K3 direct electron detector with Bioquantum energy filter (Gatan) and a slit width of 15 eV. Images were recorded using the automated acquisition program EPU (ThermoFisher) in super-resolution mode at a magnification of 81,000, corresponding to a pixel size of 0.44 Å/pixel on the specimen level. 13,290 and 11,321 movies were acquired respectively, in a defocus range of −1.5 to −3.0 μm. Each movie comprised 60 frames acquired over a total integration time of 3.0 s with a total cumulative dose of ∼ 57 e^-^/Å^2^ each.

### Cryo-EM image processing of ABC(K567W/K2008W)

Movie frames were aligned, dose-corrected, binned twice to 0.88 Å/pixel and averaged using MotionCor2^63^ and CTF parameters were estimated using CTFFIND4^60^ both implemented in TranSPHIRE^64^. 426,857 and 509,274 particles were auto-picked respectively, using crYOLO^60^ with a general particle model and then extracted with a box size of 560 pixels.

Several rounds of 2D classification were performed with ISAC^61^ in SPHIRE^62^ with a pixel size of 6.1 Å/pixel on the particle level, before and after particle centring, re-extraction and combination of the datasets. In total, the two datasets yielded 61,817 and 29,942 prepore and pore particles of TcA respectively. The 61,817 prepore particle images were further processed iteratively with 2D classifications combined with 3D refinements to obtain centred, high-quality holotoxin classes. Here, the rotation/translation parameters obtained from MERIDIEN 3D refinements were used to improve 2D classification in ISAC. After the final ISAC run, the ‘Beautifier’ tool of SPHIRE^62^ was applied to obtain refined and sharpened 2D class averages at the original pixel size, showing high-resolution features (Supplementary Figure 16a,b). Ultimately, 13,356 particles were taken for 3D refinement with Relion 3.1^66^ using the density map of TcdA1 WT (EMDB ID 10033), low-pass filtered to 20 Å, as initial reference and a mask that includes the well-resolved part of the reconstruction. This resulted in an average resolution of 3.3 Å according to FSC 0.143 (Supplementary Figure 16c).

As observed previously with TcA(1279-At647N) and TcA(K1179W), the obtained reconstruction lacked density for the receptor-binding domains B and C. However, at low binarization threshold, this map also revealed an additional density below RBD D, next to the neuraminidase-like domain. Subsequently, we performed 3D classification in Relion 3.1^66^ with a mask focusing on the region surrounding the additional density (Supplementary Figure 16f). The resulting 3D classes were relatively homogenous and could therefore all be taken for additional CTF and local refinement. Since the presence of the TcB-TcC cocoon did not show any effect on prepore-to-pore kinetics (Supplementary Figure 3c), the final post-processing step was performed with a mask on the TcA pentamer only to yield a 3.2 Å map according to FSC 0.143 (Supplementary Figure 16f).

### Atomic model building and refinement

For TcA(1279-At647N) at pH 11.2, we first rigid-body fitted the model of the TcA-WT prepore (PDB 6RW6) into the obtained density map of the intermediate using UCSF Chimera^67^ and truncated residues 21 - 93 (N-terminus) and 1309 - 1580 (RBDs B, C), for which no density was present. At the region corresponding to RBD A (residues 298 – 432), the reconstruction showed a density turned outward away from the main density at an angle of 20°. To determine if this density indeed belonged to RBD A, we first predicted the model of RBD A using the protein prediction software Alphafold 2^68^. Subsequently, the resultant best model was rigid-body-fitted into the density of the cryo-EM map and joined with the rest of the model after manual adjustment in Coot^69^. Then, the combined model was refined using ISOLDE^70^, followed by the real space refinement tool of PHENIX^71^.

Next, since the observed extra density lacked high resolution features that would allow tight fitting and model building, we performed map subtraction of the best 3D class of TcA(1279-At647N) showing this density from the global EM density map to obtain the difference map of the elongated density. Then, we low-pass-filtered this difference map to 10 Å and aligned it to the reconstruction and fitted model of TcA(1279-At647N). This elongated density fit to the space below the region corresponding to RBD B suggesting the RBD B (and perhaps also RBD C) were flipped out. Therefore, we rigid-body fitted residues 1309-1361 and 1493-1576 corresponding to RBD B into the additional density of all the maps of the TcA intermediate state, but could not fit RBD C due to insufficient density. The difference map showing the additional density corresponding to RBD B was not included in the density maps submitted to EMDB.

Atomic models of ABC(K567W/K2008W) and TcA(K1179W) were built based on the TcA(1279-At647N) model following the same procedure. Briefly, the model of TcA(1279-At647N) was first rigid-body fitted into the obtained density maps of the prepore-to-pore intermediates using UCSF Chimera^67^. Here, the relevant mutations were added, before the models were manually adjusted in coot and subsequently refined in ISOLDE and PHENIX. The data statistics of all obtained models are summarized in Supplementary Table 6.

