## Supplementary Figures/Tables/Notes for "Multi-state kinetics of the syringe-like injection mechanism of Tc toxins"

**Supplementary information**

### Supplementary Figures

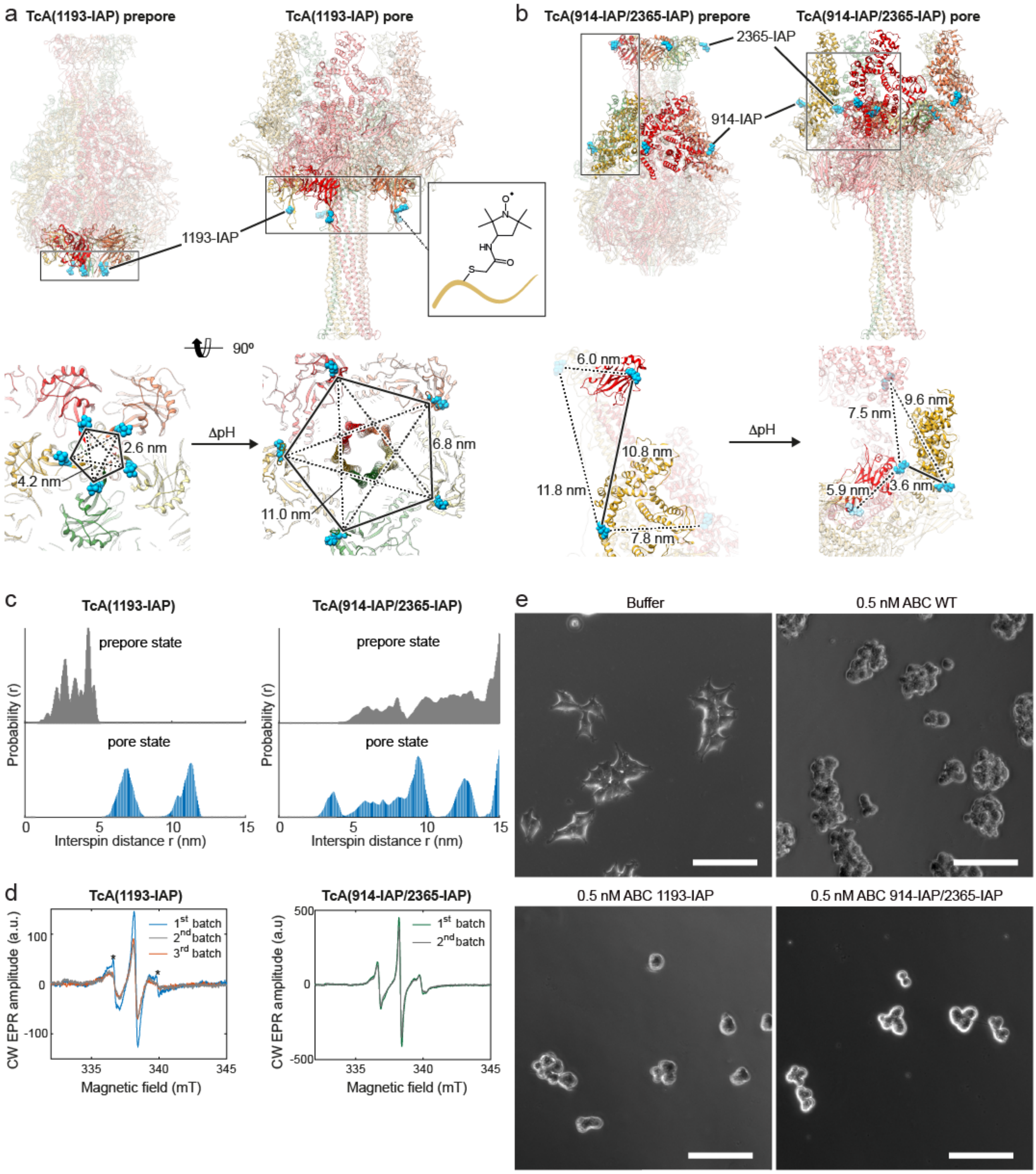

**Supplementary Figure 1: Design of TcA variants for EPR, labeling efficiency determination, distance distribution simulation and toxicity of IAP-labeled TcA variants to HEK293T cells. a: Top panel**: TcA(1193-IAP) designed to monitor the opening of the shell. **Inset**: IAP label attached to a cysteine. **Bottom panel**: bottom view of the TcA prepore (left) and pore (right) with the average mean distances between the 1193-IAP labels indicated (dashed and solid lines). **b: Top panel**: TcA(914-IAP/2365-IAP) designed to monitor channel ejection. **Bottom panel**: close-up view of the labeling sites of two adjacent protomers in the prepore (left) and the pore (right) with the distances between the IAP labels at 914Cys and 2365Cys indicated. **c:** Simulation of the distance distribution between the IAP labels in TcA(1193-IAP) (**left panel**) and TcA(914-IAP/2365-IAP) (**right panel**) in the prepore (upper panels, grey) and pore state (lower panels, blue), calculated using rotamer library approach with MMM[1,2]. **d**: X-band CW EPR spectra of different batches of TcA(1193-

IAP) (**left panel**) and TcA(914-IAP/2365-IAP) (**right panel**) variants as indicated. Peaks corresponding to remaining free label are indicated with *. Corresponding labeling efficiencies were obtained by spin counting of spectra and are summarized in Supplementary Table 1. **e:** Intoxication of HEK293T cells with 0.5 nM holotoxins formed by TcA WT, TcA(1193-IAP), or TcA(914-IAP/2365-IAP) and TcB-TcC WT. Intoxicated cells rounded up and detached from the surface. Scale bars, 100 µm.

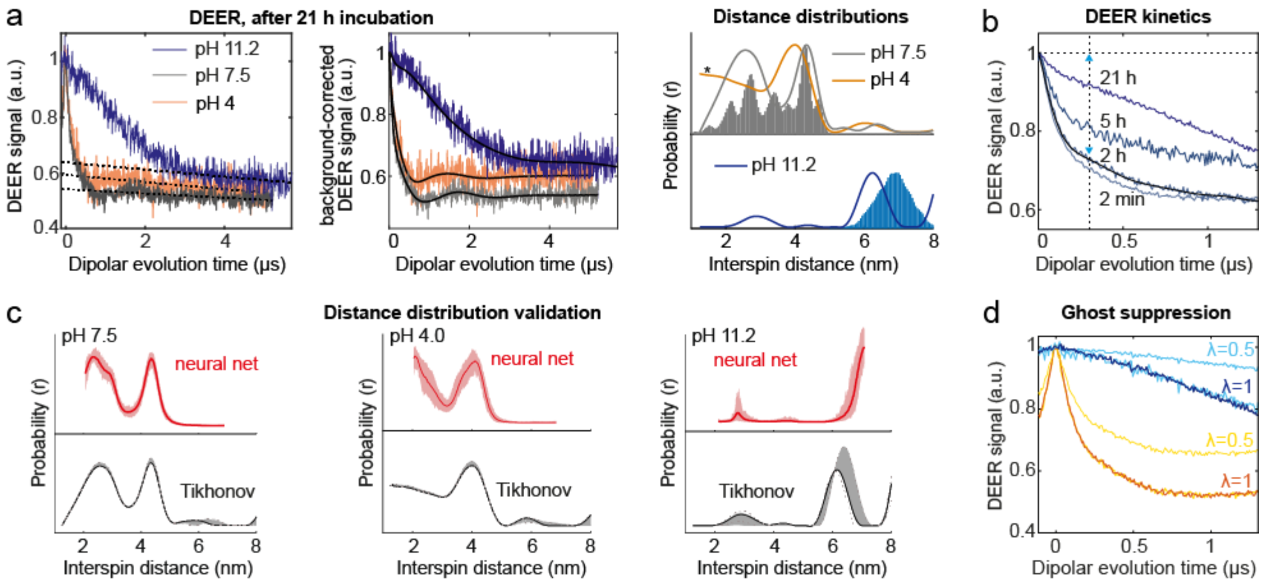

**Supplementary Figure 2: DEER data validation, DEER snapshot kinetic and ghost peak suppression for TcA(1193-IAP). a**: Primary DEER data, background corrected and corresponding distance distributions obtained by Tikhonov regularization in DeerAnalysis2019[3] for TcA(1193-IAP) after 21 h incubation at 21 ºC in pH 7.5, 4.0 and 11.2 buffers. **Left panel**: User-defined background correction, shown in a black dotted line. **Middle panel**: fits of background-corrected data shown in black solid lines. **Right panel**: normalized experimental data (solid lines) and simulations (shaded areas) of the distance distributions for TcA(1193-IAP). Data for pH 4.0 and pH 7.5 and simulations of the prepore (grey) are shown in the top part; data for pH 11.2 and simulations for the pore state (blue) are shown in the bottom part. **b**: DEER "snapshot" kinetic traces of TcA(1193-IAP) after different incubation times at pH 11.2. Polynomial fit is shown in black and used to calculate the derivatives to obtain 1$^{st}$ inflection time point (indicated as dotted line) for the determination of the modulation of short distances. **c**: Validated distance distributions for pH 7.5, 4.0 and 11.2 (**left, middle and right panels**, respectively) obtained with Tikhonov regularization (lower black curves, errors are shown as grey areas) are compared with analysis using a neural network in DeerAnalysis2019[3,4] (upper red curves, errors shown in black). **d**: Short primary DEER traces of pH 4.0 (orange and yellow curves) and pH 11.2 (light and dark blue curves) with π pump pulse (λ=1) and reduced pump pulse (λ=0.5, cyan and yellow) shown as detected and scaled λ = 1. The identical trace shapes for both λ values indicate that there are no ghost frequencies, artefacts.

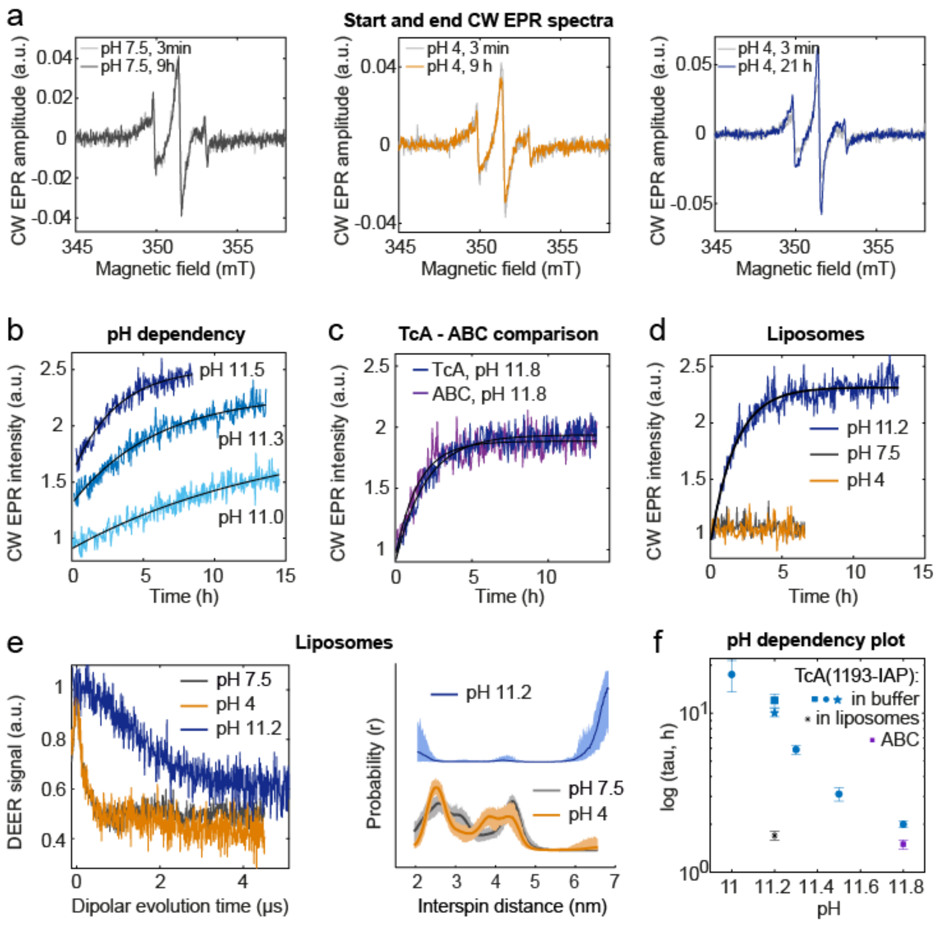

**Supplementary Figure 3: pH dependency and influence of liposomes on prepore-to-pore transition kinetics of TcA(1193-IAP) monitored by EPR. a**: X-band CW EPR spectra for TcA(1193-IAP) at the start (light grey) and end (colored) incubation period of 9 h at pH 7.5 (grey**, left panel**), pH 4.0 (orange, **middle panel**) and after 21 h at pH 11.2 (blue, **right panel**). **b**: CW EPR kinetics of TcA(1193-IAP) (batch 2 from Supplementary Figure 1d) at different basic pH values. Mono-exponential fits are shown as black solid lines with the reaction times given in Supplementary Table 2. **c**: Comparative CW EPR kinetics of holotoxin ABC(1193-IAP) and TcA(1193-IAP) at pH 11.8. The black solid lines represent mono-exponential fits with the time constants given in Supplementary Table 2. **d, e**: TcA(1193-IAP) in liposomes. CW EPR kinetics (**d**) and DEER (**e**; primary, **left panel**, and distance distributions, **right panel**) at the indicated pH values. DEER measurements were performed on samples after overnight incubation. **f**. Reaction times obtained from mono-exponential fits of CW kinetics, plotted logarithmically against pH values. The three different symbols for TcA in buffer represent the independent experiments performed with the three protein batches.

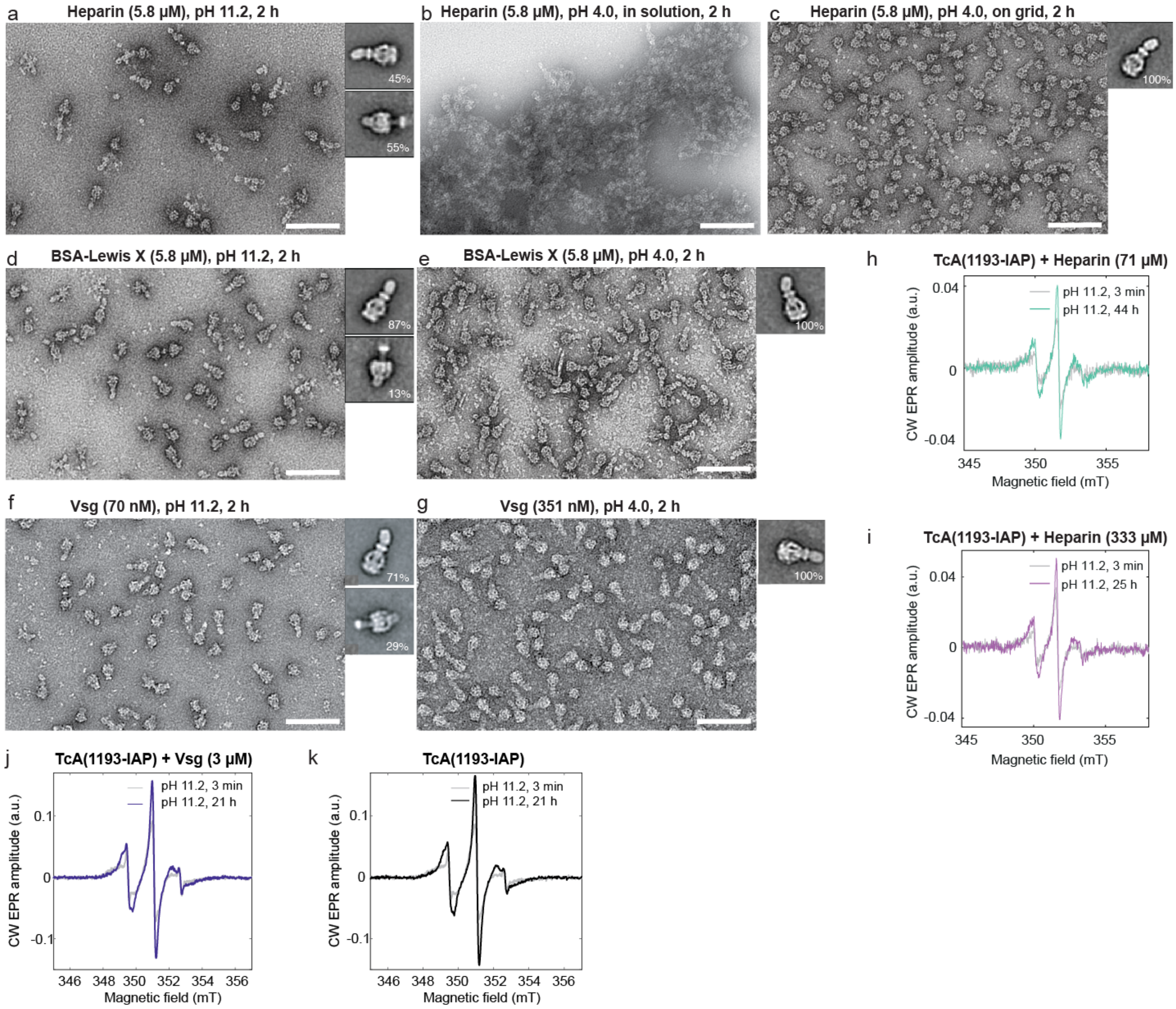

**Supplementary Figure 4: Influence of known receptors on prepore-to-pore transition kinetics of TcA. a-g:** Negative stain EM images and representative 2D class averages (right) of prepore-to-pore transition experiments of the ABC holotoxin (TcA WT and TcB-TcC) in the presence of its receptors heparin (**a-c**), BSA-Lewis X (**d**, **e**) and Vsg (**f**, **g**). The toxin was preincubated for 1 h at 21 °C with the receptors at the indicated concentrations, followed by 2 h incubation at either pH 11.2 or pH 4.0. Prepore-to-pore transition experiments at pH 11.2 were all performed in solution, while those at pH 4.0 were either in solution (**b, e, g**) or on a negative stain grid (**c**) to avoid precipitation. Scale bars in panels a and c-g are 100 nm and 140 nm for b. **h, i:** X-band CW EPR spectra of TcA(1193-IAP) + heparin at pH 11.2 after the kinetic (Figure 2b) start, 3 min (grey), and end (colored). The experiment was started after 1 h preincubation of 14.2 µM TcA (protomer) at 21 °C with 71 µM (**h**) or 333 µM (**i**) of heparin. **j, k**: X-band CW EPR spectra of TcA(1193-IAP) + Vsg (**j**) and TcA(1193-IAP) alone (**k**) at pH 11.2 after the kinetic (Figure 2c) start, 3 min (grey) and end (colored). The experiment was started after 1 h preincubation of 3 µM TcA (pentamer) 1:1 with Vsg at 21 °C.

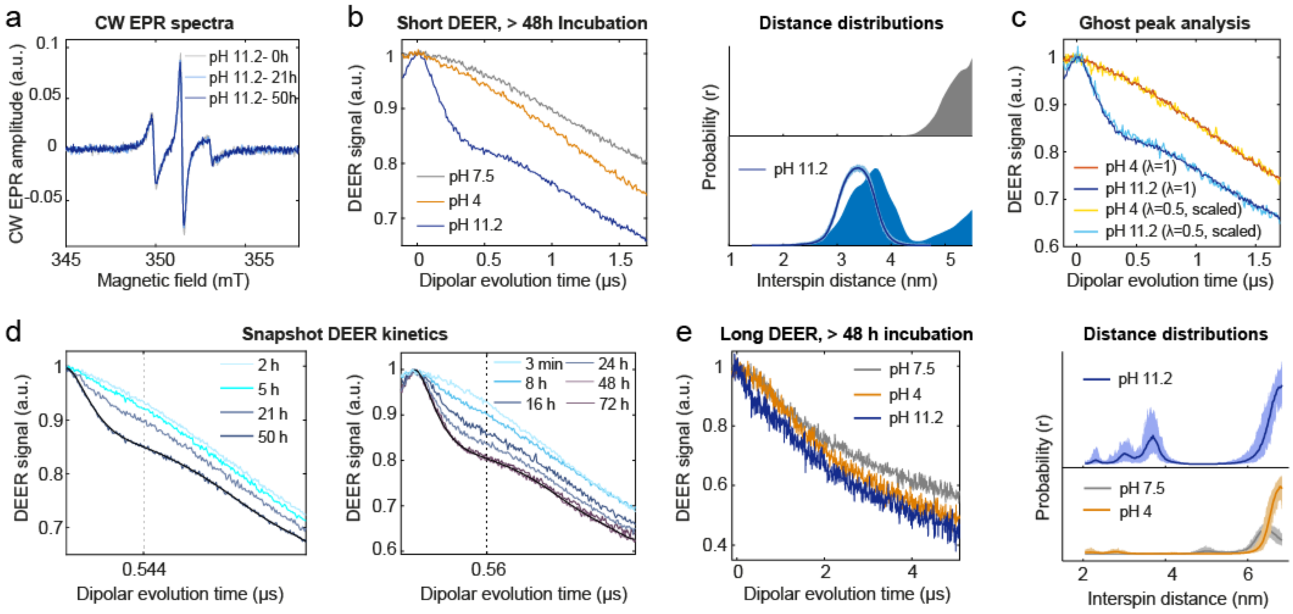

**Supplementary Figure 5: CW EPR spectra, DEER data analysis and ghost peak suppression for TcA(914-IAP/2365-IAP). a**: X-band CW EPR spectra of TcA(914-IAP/2365-IAP) at pH 11.2 after 0, 21 and 50 h incubation at 21 ºC. **b:** Short primary DEER traces (**left panel**) and corresponding distance distributions (**right panel**) of TcA(914/2365-IAP) after 48 h of incubation at 21 ºC, pH 11.2, and 72 h incubation at 4 ºC, pH 4 and pH 7.5. The corresponding simulations are shown as filled areas. **c**: Ghost peak analysis of short primary DEER traces performed with reduced pump pulse (λ = 0.5, yellow and light blue lines, shown scaled) indicating identical shapes to λ = 1 traces (orange, dark blue lines). **d**: DEER traces for $1^{st}$ (**left panel**) and $2^{nd}$ (**right panel**) TcA(914-IAP/2365-IAP) batches at pH 11.2 at 21 ºC used to reconstruct DEER "snapshot" kinetic in Figure 1c. The inflection point is marked on the x-axis. **e**: Long primary DEER traces (**left panel**) and corresponding distance distributions (**right panel**) of TcA(914/2365-IAP) at the indicated pH values.

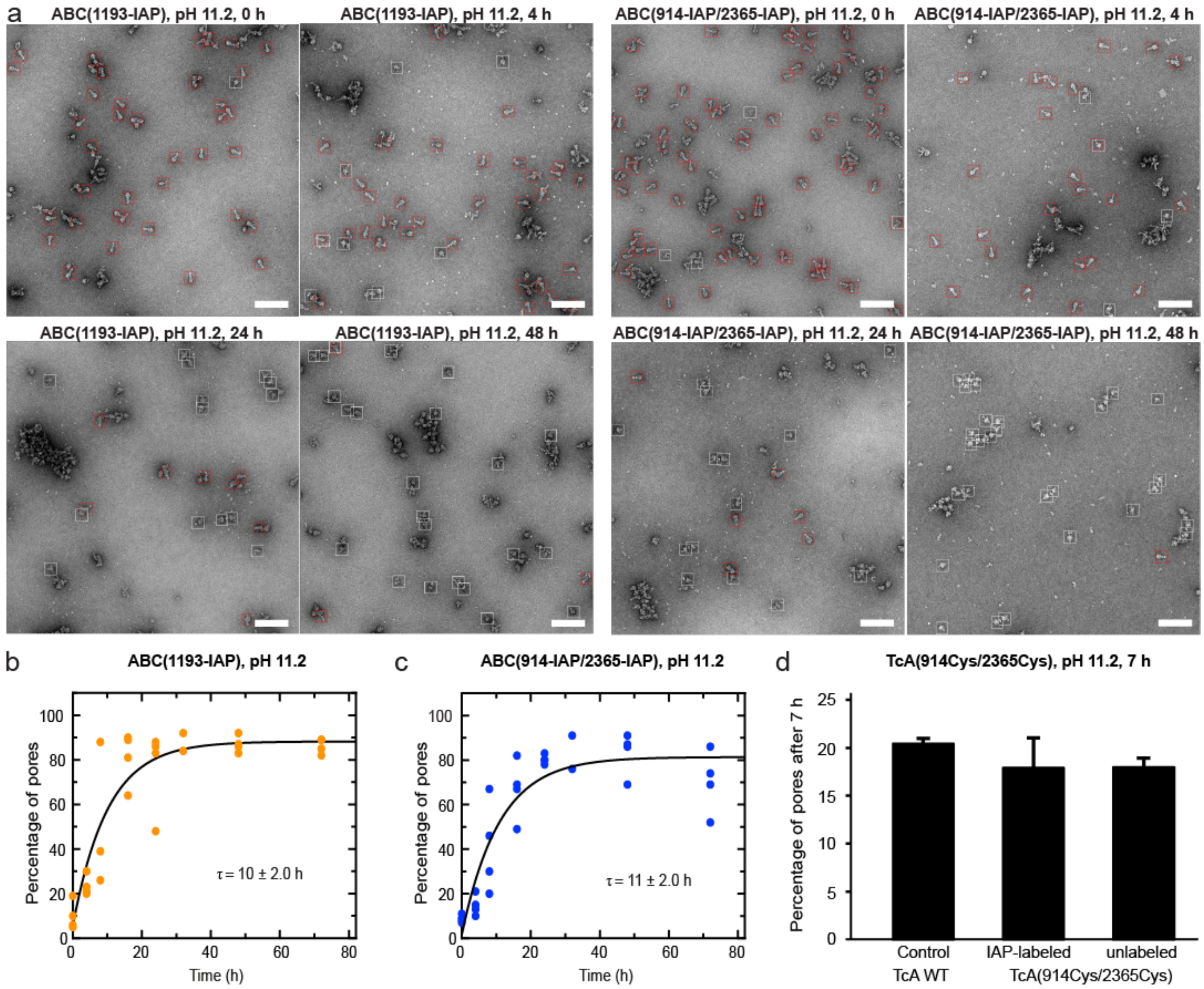

**Supplementary Figure 6: Pore formation quantification of ABC(1193-IAP), ABC(914-IAP/2365-IAP) and TcA(914Cys/2365Cys) at pH 11.2 by negative stain EM. a:** Representative negative stain micrographs of ABC(1193-IAP) and ABC(914-IAP/2365-IAP), respectively after 0 h, 4 h, 24 h or 48 h of incubation at pH 11.2 and 21 ºC. Scale bars are 100 nm. Prepores and pores were picked using crYOLO[5] with specifically trained models and are indicated with red and white boxes, respectively. **b-c**: Quantification of the number of pores from **a** after picking and subsequent 2D classification, ABC(1193-IAP) (**b**) and ABC(914-IAP/2365-IAP) (**c**). **d**: Quantification of the number of pores of labeled and unlabeled TcA(914Cys/2365Cys), compared to wild-type TcA (TcA WT) after incubation at pH 11.2 for 7 h. The data are presented as mean values and the error bars represent the standard error of the mean (SEM) of three independent experiments.

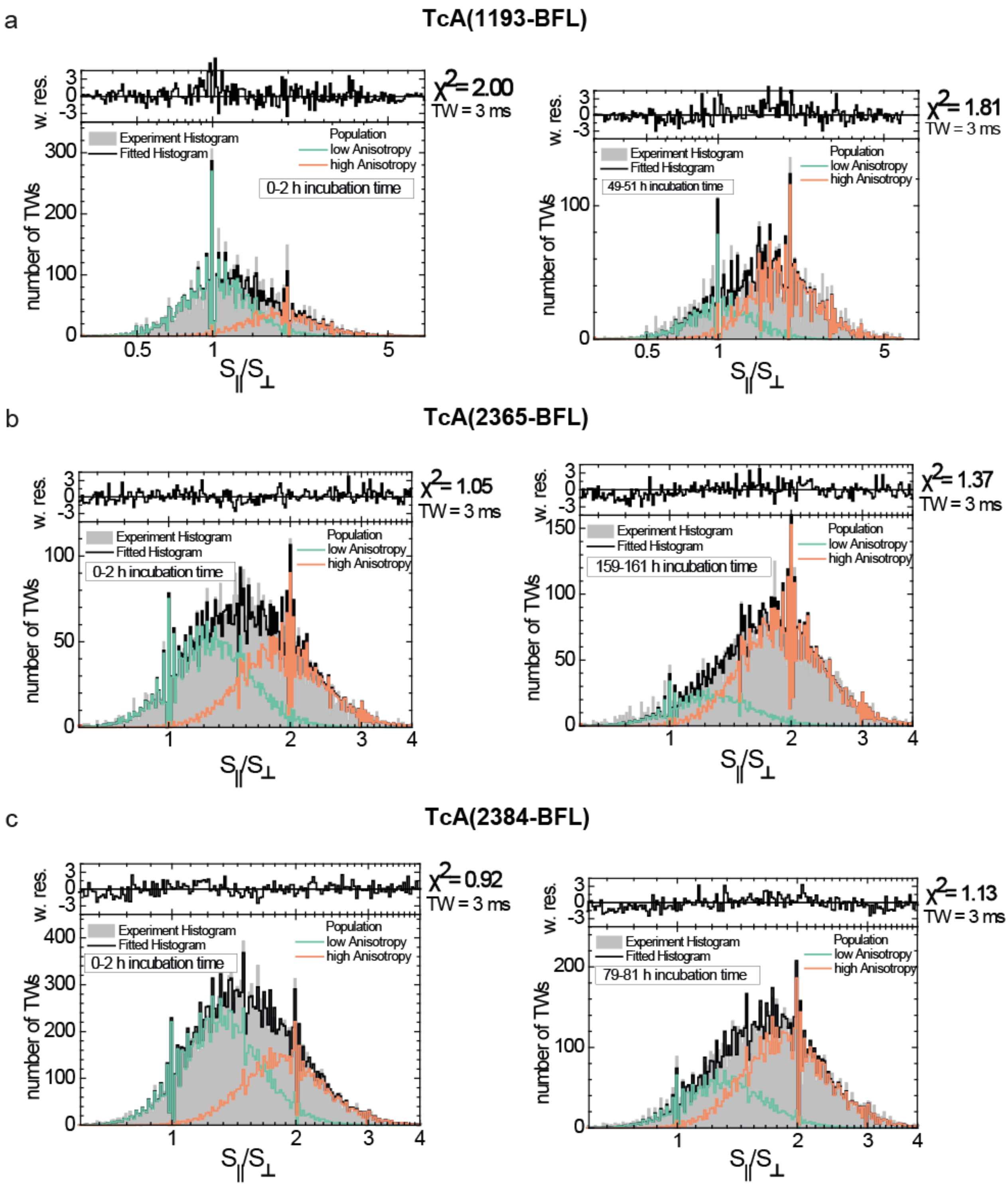

**Supplementary Figure 7: Anisotropy Photon Distribution analysis of freely diffusing single TcA subunits for different time segments.** The histograms of the ratios of parallel and perpendicular signals $S_{||}$ and $S_{\perp}$ of each burst with a time window TW = 3 ms were analysed as described in Supplementary Methods. Start and end point of long-time measurement series shown. The kinetic analysis of obtained fractions is shown in Figure 1e and is described in Supplementary Note 1 for TcA(1193-BFL) (**a**), monitoring shell opening, as well as TcA(2365-BFL) (**b**) and TcA(2384-BFL) (**c**), both monitoring channel ejection. The data are well described by a two-state model (black) with a superposition of a low anisotropy state (lr, green) and high anisotropy state (hr, orange). The quality of the fit was judged by weighted residuals (upper panels) and chi-squared analysis. The fit results are compiled in Supplementary Table 3. In the case of TcA(2365-BFL) and TcA(2384-BFL), high anisotropy state (hr = 0.29) is caused by environmental change from the surrounding opened shell/ejected pore and is therefore interpreted as pore state. For TcA(1193-BFL), high anisotropy state (r = 0.27) is caused by environment change and reduction of homoFRET, indicating an opening of the shell (see Supplementary Figure 11).

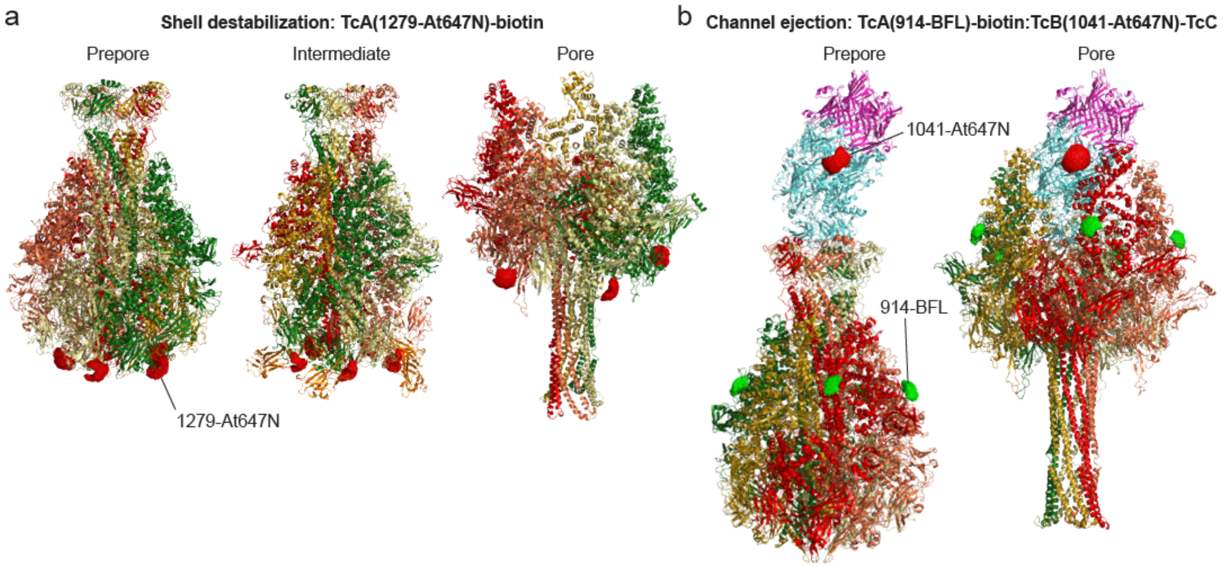

**Supplementary Figure 8: Representation of accessible volumes (AV) of the fluorescent labels as described in Methods a**: 3D structures of TcA prepore, intermediate and pore with AV of Atto647N attached to TcA(1279Cys). **b**: 3D structures with AV of BFL at TcA(914Cys) and Atto647N at TcB(1041Cys)-TcC, plotted on the ABC holotoxin.

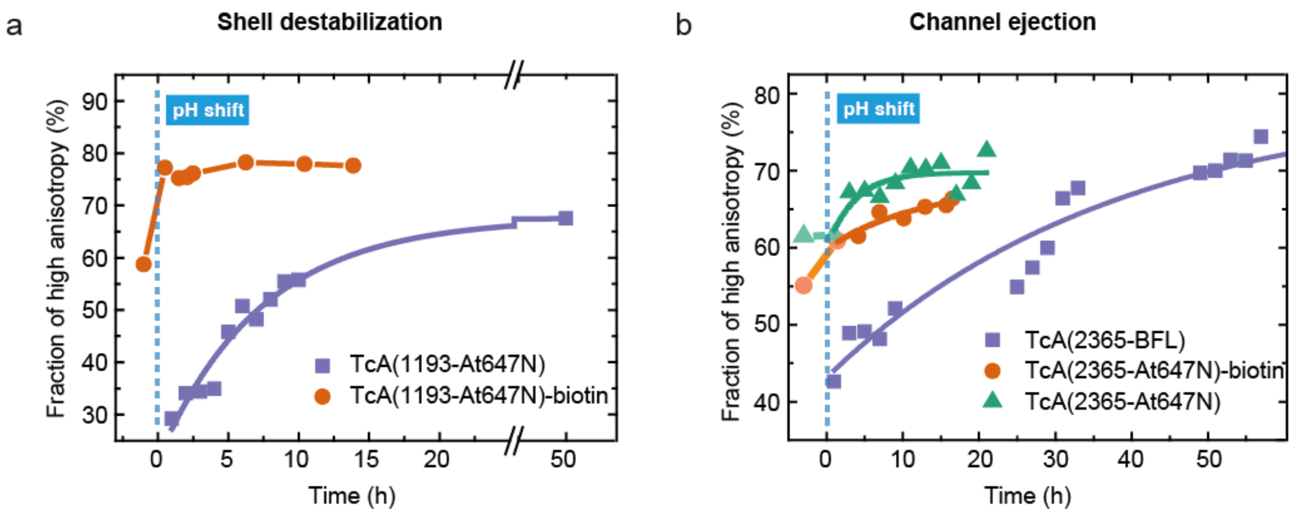

**Supplementary Figure 9: Prepore-to-pore transition kinetics of biotinylated TcA**. **a**: Shell destabilization. Consistent with the pore ejection kinetics, TcA(1193-At647N)-biotin shows an instant (less than several minutes) fraction change after pH shift, whereas the non-biotinylated sample shows a slow fraction change. **b**: Channel ejection, monitored using TcA(2365-BFL), biotinylated TcA(2365-At647N) and non-biotinylated TcA(2365-At647N). For TcA(2365-At647N)-biotin, most of the kinetics, i.e., change of fraction from prepore to pore, happen shortly after the pH shift (dashed blue line). Reaction times are compiled in Supplementary Table 5.

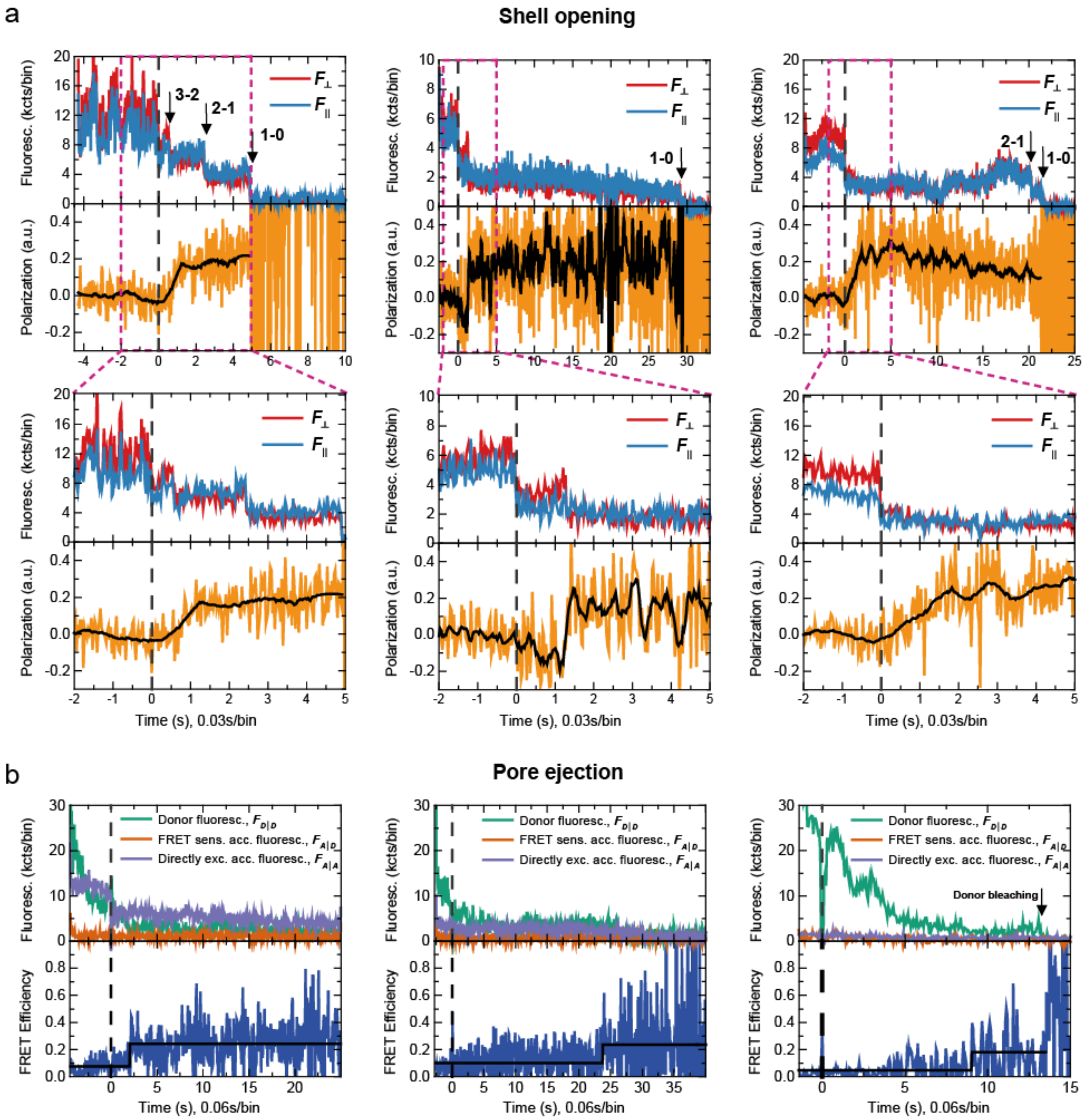

**Supplementary Figure 10: Example traces of pore ejection and shell opening. a:** Selected traces of the shell opening using an anisotropy assay that senses local environment and homoFRET, as shown in Figure 3d. TcA(1279-At647N)-biotin was labelled with a ratio of four dyes per pentamer. The top layer is displaying the background-corrected perpendicular polarized Fluorescence $F_{\perp}$, and the parallel polarized Fluorescence $F_{\|}$. Dashed magenta lines are indicating the zoom interval for the lower diagram. Arrows mark the intensity drop due to bleaching with the numbers of fluorophores attached to the toxin in bright state before and after the bleaching event. Bottom layer is showing the polarization value (orange) and the smoothed polarization trajectory using the Savitzky-Golay algorithm (black). **b**: Selected traces of the pore ejection, as shown in Figure 3e. TcA(914-BFL)-biotin:TcB(1041-At647N)-TcC was labeled with a ratio of one BFL per TcA-pentamer so that heteroFRET for a single dye pair was measured. Top layer is displaying the background, crosstalk and detection-ratio-corrected fluorescence of the donor ($F_{D|D}$), the FRET-sensitized acceptor ($F_{A|D}$) and of the directly excited acceptor ($F_{A|A}$). The black line in the bottom layer is showing the average efficiency value before and after the transition, approximately matching the expected efficiencies calculated using 3D structures of the prepore and pore states (Supplementary Table 4). The dashed vertical black line is indicating the pH change, offset corrected to t = 0 s. After a relaxation time, the jump is within a single bin (bin width = 58.88 ms). Before and after the transition, the FRET efficiency stays at a constant average value until bleaching occurs.

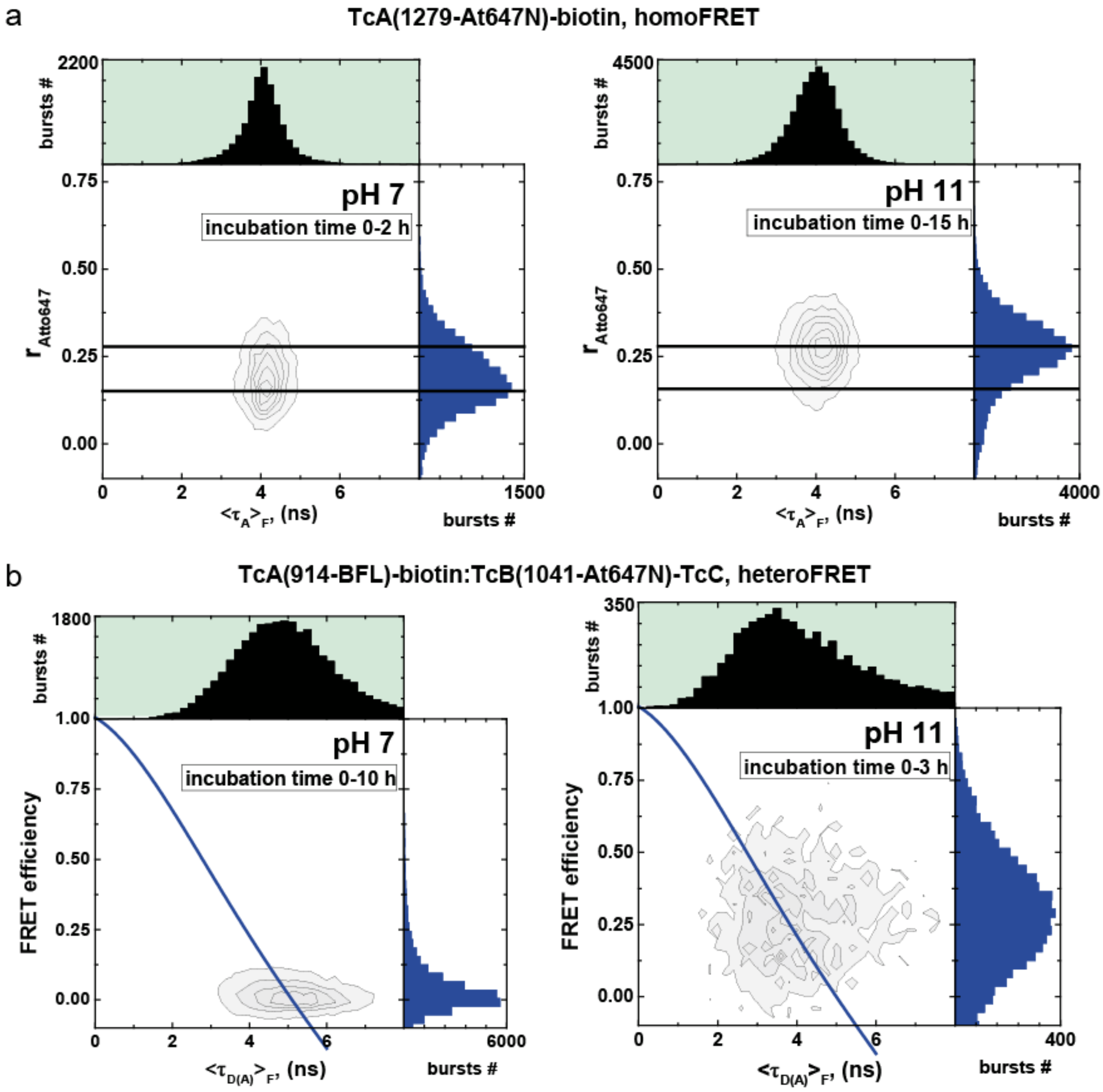

**Supplementary Figure 11: Multi-dimensional smFRET histograms of TcA(914-BFL)-biotin:TcB(1041-Atto647)-TcC and TcA(1279-At647N)-biotin**. **a:** Two-dimensional histogram of scatter-corrected anisotropy and lifetime $\langle\tau A\rangle f$ of TcA(1279-At647N)-biotin. Anisotropy values are corrected for background. The horizontal lines show the mean value of anisotropy under pH 7 environment (r=0.15) and pH 11 environment (r=0.27). **b**: Two-dimensional histogram FRET efficiency E vs lifetime of donor in the presence of acceptor $\langle\tau D(A)\rangle f$. One-dimensional histograms are the projected burst distributions over a single variable. Acceptor and donor only bursts were filtered out using stoichiometry and Alex 2CDE filter. FRET efficiency values are corrected for background, spectral crosstalk, direct acceptor excitation and detection efficiency ratio. Static FRET line (blue line) was calculated using [1] $E(\langle\tau D(A)\rangle f)=1-(((0.0030*\langle\tau D(A)\rangle f4)+(-0.0535*\langle\tau D(A)\rangle f3)+(0.3118*\langle\tau D(A)\rangle f2)+0.4093*\langle\tau D(A)\rangle f1-0.0270)/\langle\tau D(0)\rangle f)$.

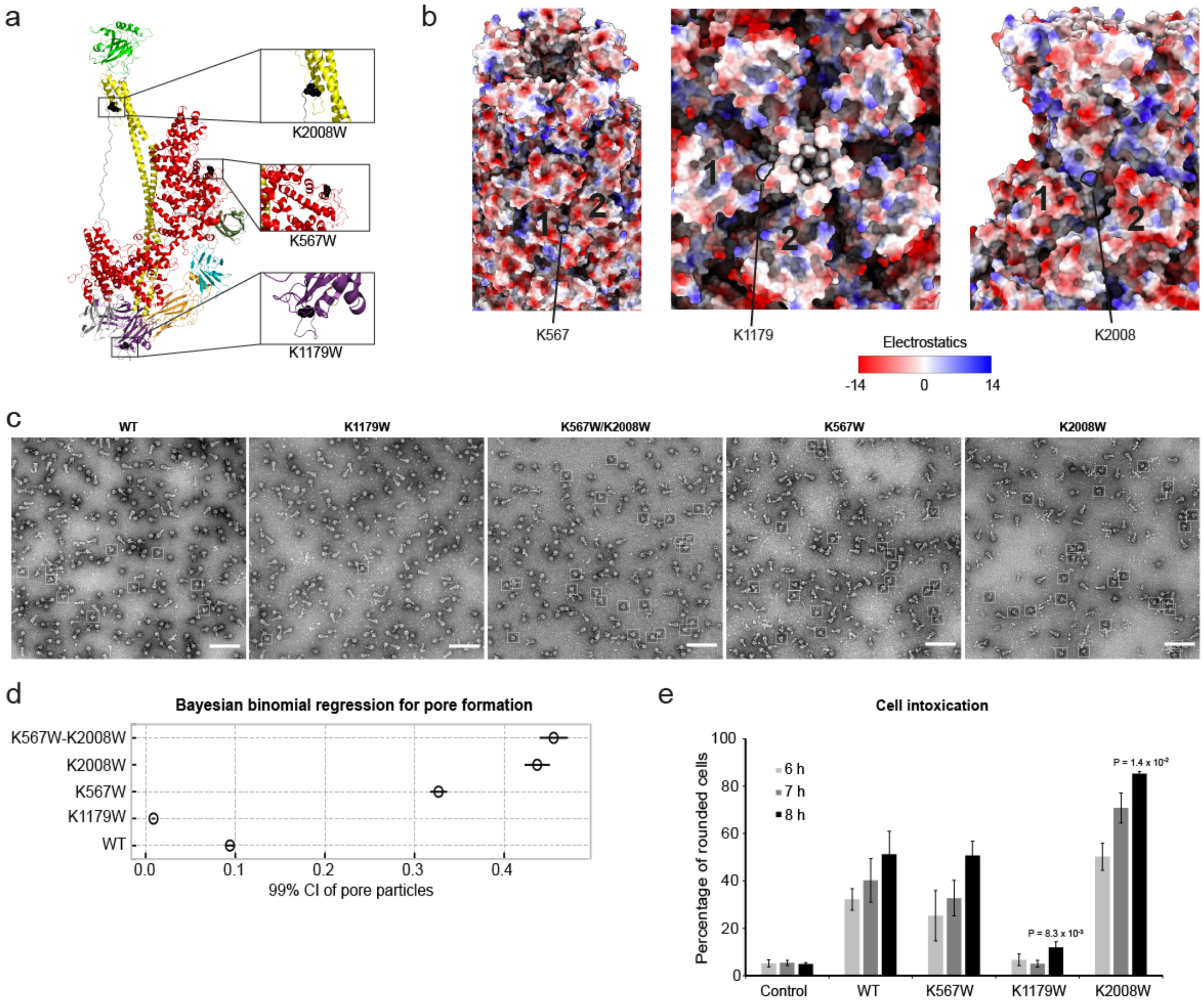

**Supplementary Figure 12: TcA mutants mimicking biotin-induced prepore-to-pore transition acceleration. a:** Structure model of a TcA WT monomer (PDB: 6RW6) showing the surface-exposed lysines selected for directed mutagenesis. **b**: Surface representations of TcA WT at pH 11 showing the target lysines to be mutated (black frames). The numbers show the two adjacent protomers near the residue of interest. The models are coloured according to their Coulomb potential at pH 11 following predictions using the H++ server. Positively charged (14 kcal/mol) and negatively charged (-14 kcal/mol) residues are colored in blue and red, respectively. **c**: Representative negative stain EM images of all mutants after 2 h of incubation at pH 11.2 and 21 °C. Scale bars are 100 nm. Prepores and pores were picked using a specifically trained model in crYOLO and the pores are indicated with white boxes. **d**: Bayesian binomial regression analysis of pore formation following 2D classification of all prepores and pores for each mutant. All particles were assigned a category (prepore or pore) according to the corresponding 2D class and subsequently counted. This enabled calculation of a 99% credibility interval (CI) for the probability of obtaining pores from each mutant. Priors were uninformative and normally distributed. **e**: Intoxication of Hela cells with holotoxins (5.28 nM) formed by the indicated TcA variants, TcA WT and TcB-TcC WT. Intoxicated cells rounded up and the number of rounded cells was quantified. The data are presented as mean values and the error bars represent standard error of the mean (SEM) of three independent experiments.

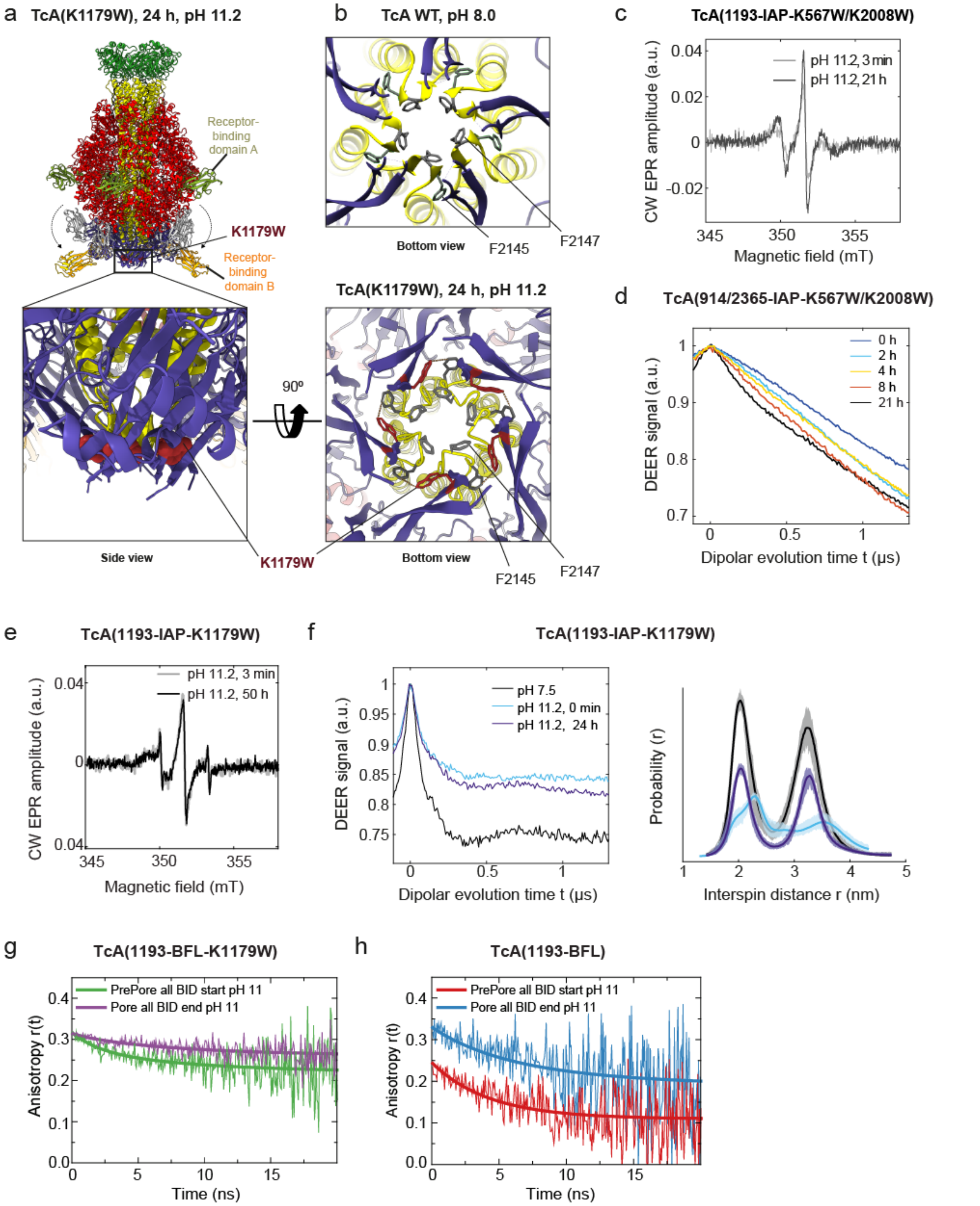

**Supplementary Figure 13: CW EPR and DEER spectra of TcA(914/2365-IAP-K567W/K2008W) and TcA(1193-IAP-K1179W) at pH 11.2. a:** Structure model of the TcA(K1179W) intermediate in cartoon side view showing the location of the biotin-mimicking K1179W mutation (represented as maroon spheres) and the flipped receptor-binding domains (RBDs) A (green) and B (orange). The RBD B facing the front has been omitted from the model to allow a better view of K1179W. **Bottom-left panel**: a close-up bottom view of the neuraminidase-domain showing the position of the K1179W mutation (maroon) presented as spheres. **Bottom-right panel**: bottom view of the location of the K1179W mutation (maroon) and the neighboring F2145 and F2147 (grey) on the channel to which it forms hydrophobic interactions. **b**: Bottom view of the TcA WT prepore model (PBD: 6RW6) showing the downward orientation of F2145 (grey, stick representation), which is in contrast to its upward orientation in the TcA(K1179W) intermediate (panel a (bottom-right)). **c**: X-band CW EPR spectra for TcA(1193-IAP-K567W-K2008W) at the start and end of a 21 h incubation period at pH 11.2 at 21ºC. The spectra are the start and end points of the kinetics in Figure 4b. **d**: Primary DEER data for TcA(914/2365-IAP-K567W/K2008W) after indicated incubation times at room temperature at pH 11.2. **e**: X-band CW EPR spectra for TcA(1193-IAP-K1179W) at the start (light grey) and end (black) incubation period of 50 h at pH 11.2. **f**: **Left panel**: primary DEER data for TcA(1193-IAP-K1179W) without incubation at room temperature at pH 7.5 and 11.2 and after 24h incubation at pH 11.2. **Right panel**: the corresponding distance distributions from the left panel. **g-h**: Fluorescence anisotropy measurements of TcA(1193-IAP-K1179W) (**g**) at pH 11.2 compared to the wild-type TcA(1193-IAP) (**h**).

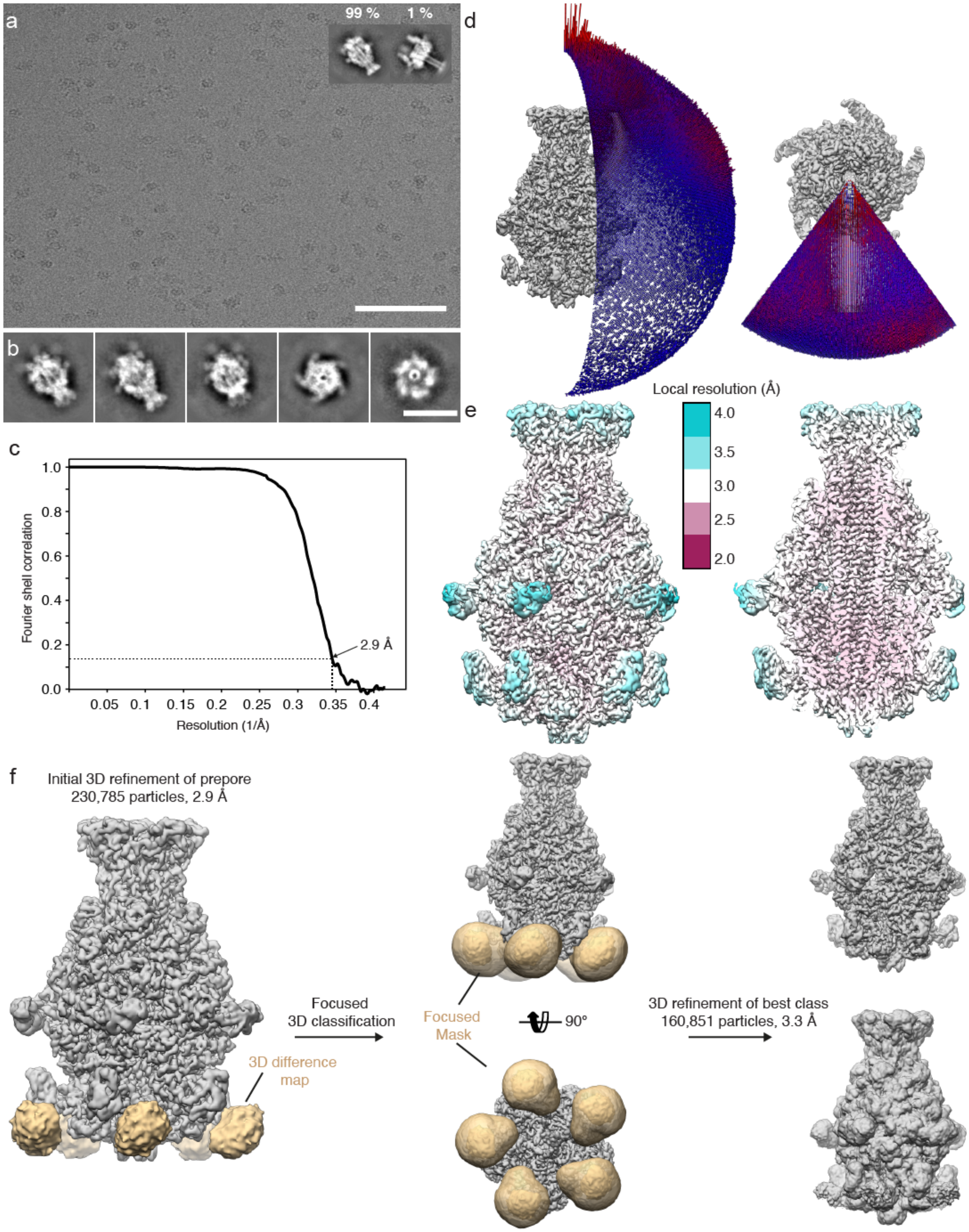

**Supplementary Figure 14: Cryo-EM of TcA(K1179W) after 24 h at pH 11.2. a**: Typical digital micrograph area of vitrified TcA(K1179W) at a defocus of 2 µm and a total dose of 56 $e^{-}$ $Å^{-2}$ acquired with a Falcon III direct electron detector. Scale bar is 100 nm. Inset: Representative 2D class averages, showing a prepore-shaped intermediate (99% of all particles) and a pore (1% of all particles). **b**: Representative reference-free 2D class averages obtained by ISAC and subsequently resampled to the original pixel size, refined and sharpened using the Beautifier tool implemented in the SPHIRE software package. Scale bar is 20 nm. **c**: Fourier shell correlation (FSC) of the masked cryo-EM map (black curve). The 0.143 FSC cut-off criterion indicates that the cryo-EM map has an average resolution of 2.9 Å. **d**: Angular distribution for the final round of the reconstruction. Each stick represents a projection view. Size and color of the stick is proportional to the number of particles. **e**: Surface and cross-section of the cryo-EM density map colored according to the local resolution. **f**: Workflow of focused 3D classification. The best 3D class was subjected to further local refinement, resulting in a final resolution of 3.3 Å. The final density map is shown at high binarization threshold (top-right) and low binarization threshold (bottom-right).

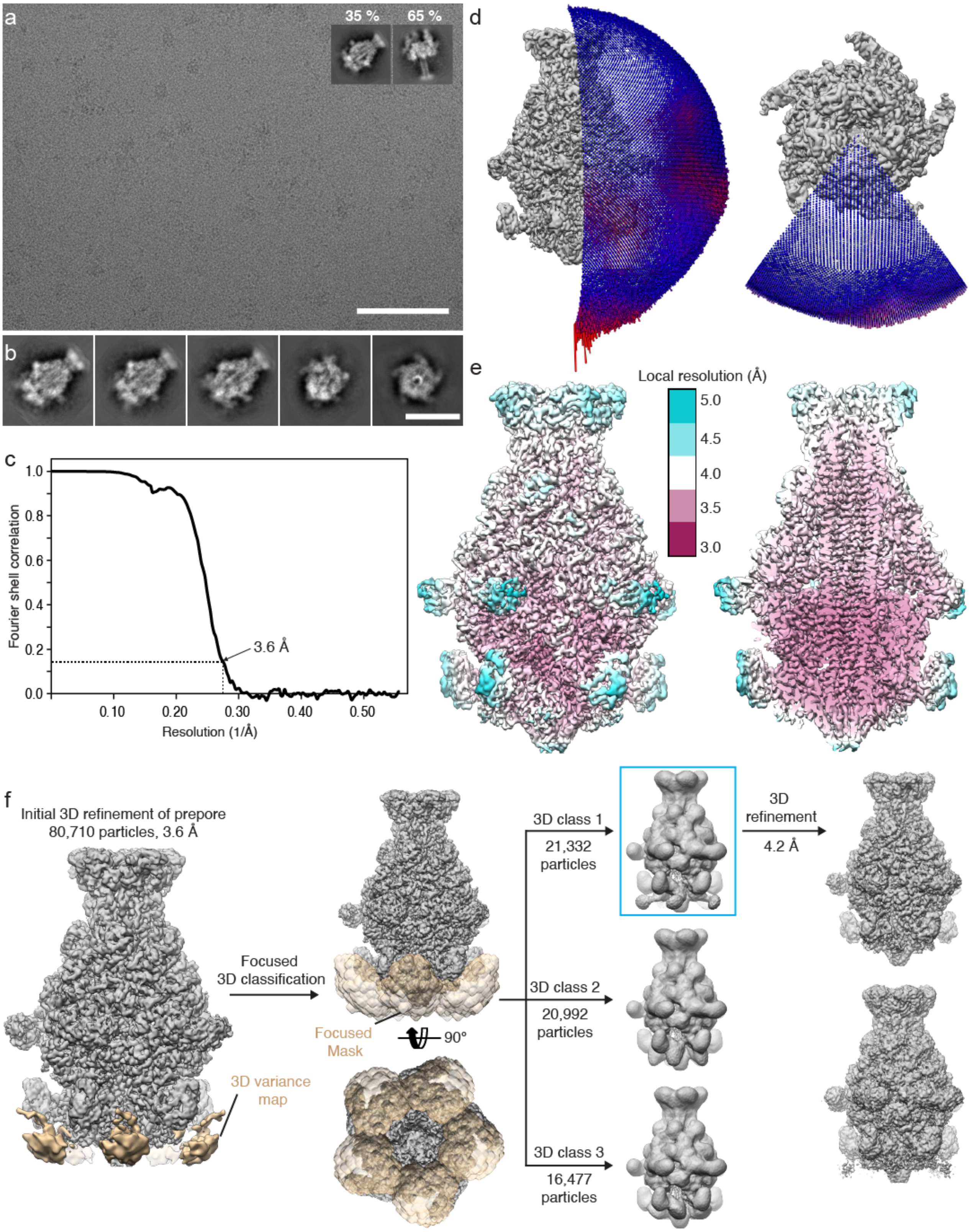

**Supplementary Figure 15: Cryo-EM of TcA(1279-At647N) after 6 h at pH 11.2. a**: Typical digital micrograph area of vitrified TcA(1279-At647N) at a defocus of 2 µm and a total dose of 78 $e^{-}$ $Å^{-2}$ acquired with a K3 direct electron detector. Scale bar, 100 nm. Inset: Representative 2D class averages, showing a prepore-shaped intermediate (35% of all particles) and a pore (65% of all particles). **b**: Representative reference-free 2D class averages obtained by ISAC and subsequently resampled to the original pixel size, refined and sharpened, using the Beautifier tool implemented in the SPHIRE software package. Scale bar, 20 nm. **c**: Fourier shell correlation (FSC) of the masked cryo-EM map (black curve). The 0.143 FSC cut-off criterion indicates that the cryo-EM map has an average resolution of 3.6 Å. **d**: Angular distribution for the final round of the reconstruction. Each stick represents a projection view. Size and color of the stick is proportional to the number of particles. **e**: Surface and cross-section of the cryo-EM density map colored according to the local resolution. **f**: Workflow of focused 3D classification. 3D class 1 (framed in box) was subjected to further local refinement, resulting in a final resolution of 4.2 Å. The final density map is shown at high binarization threshold (top-right) and low binarization threshold (bottom-right).

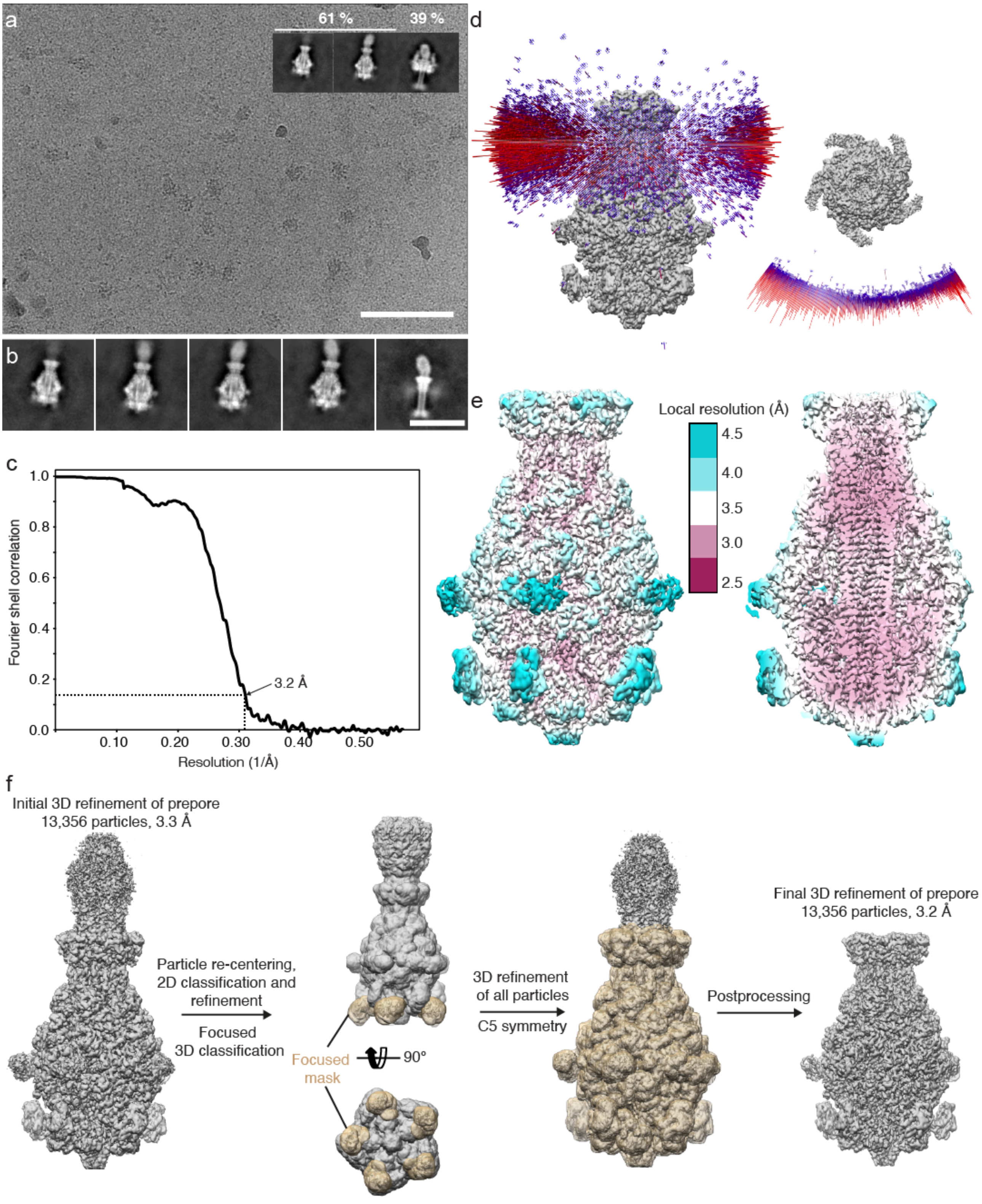

**Supplementary Figure 16: Cryo-EM of TcA(K567W/K2008W) after 2 h at pH 11.2.** **a**: Typical digital micrograph area of vitrified TcA(K567W/K2008W) at a defocus of 2 µm and a total dose of 57 $e^{-}$ $Å^{-2}$ acquired with a K3 direct electron detector. Scale bar, 100 nm. Inset: Representative 2D class averages, showing a prepore-shaped intermediate (61% of all particles) and a pore (39% of all particles). **b**: Representative reference-free 2D class averages obtained by ISAC and subsequently resampled to the original pixel size, refined and sharpened using the Beautifier tool implemented in the SPHIRE software package. Scale bar, 20 nm. **c**: Fourier shell correlation (FSC) of the masked cryo-EM map (black curve). The 0.143 FSC cut-off criterion indicates that the cryo-EM map has an average resolution of 3.2 Å. **d**: Angular distribution for the final round of the reconstruction. Each stick represents a projection view. Size and color of the stick is proportional to the number of particles. **e**: Surface and cross-section of the cryo-EM density map colored according to the local resolution. **f**: Workflow of focused 3D classification and post processing leading to the final 3.2 Å map used for model building.

### Supplementary Videos

#### Supplementary Video 1

A compilation of 2D classes of TcA(1279-At647N) showing protruding densities, thus implying conformational changes.

#### Supplementary Video 2

Transition of TcA from the prepore state (PP) upon pH change (PP*) to the stable intermediate state 1 (SI1). This video depicts the transition from PP* to SI1 as descried in Figure 5, where RBD A is flipped out by about 20º away from the shell, while RBDs B and C flip 180º. RBD C which is directly connected to RBD B is very flexible and was therefore not resolved, indicating a flexible hinge between the domains.

### Supplementary Tables

**Supplementary Table 1**: **Samples for EPR measurements**. Summary of starting protein, spin concentrations and labeling efficiencies of IAP-labelled TcA(1193Cys) and TcA(914Cys/2365Cys) (WT and K567W-K2008W variant) used in EPR studies. Spin concentrations were determined by double integration of CW EPR spectra (Supplementary Figure 1) in comparison with 100 µM Tempol standard. Labeling efficiencies represent percentage ratios of the corresponding spin concentration to introduced cysteine concentration. Errors in the labelling efficiency are estimated to be ±10%. Samples were diluted by a factor 2 for CW EPR and by a factor 4 for DEER.

| Mutant | Batch | TcA (protomer) concentration, µM | Spin total concentration, µM | Labelling efficiency (per Cys), % |
|---|---|---|---|---|
| TcA WT (1193-IAP) | 1 | 34 | 25 | 74 |
| TcA WT (1193-IAP) | 2 | 31 | 16 | 52 |
| TcA WT (1193-IAP) | 3 | 36 | 14 | 39 |
| TcA WT (914-IAP/2365-IAP) | 1 | 31 | 37 | 60 |
| TcA WT (914-IAP/2365-IAP) | 2 | 25 | 31 | 62 |
| TcA K567W/K2008W (1193-IAP) | 1 | 23 | 23 | 100 |

**Supplementary Table 2**: **Samples for EPR measurements**. Summary of reaction times of TcA(1193-IAP) (alone and as ABC holotoxin) obtained by mono-exponential fit $a*(1-\exp(-t/\tau))+b$ with no constrains of CW EPR kinetics at indicated pH values and conditions.

| **Sample** | **Batch** | **Conditions** | **pH** | **$\tau$, h** |
|---|---|---|---|---|
| TcA(1193-IAP) | 2 | buffer/DDM | 11.0 | 17.5±3.9 |
| TcA(1193-IAP) | 1 | buffer/DDM | 11.2 | 10.1±0.6 |
| TcA(1193-IAP) | 3 | buffer/Tween | 11.2 | 12.0±1.2 |
| TcA(1193-IAP) | 2 | buffer/DDM | 11.3 | 5.9±0.4 |
| TcA(1193-IAP) | 2 | buffer/DDM | 11.5 | 3.1±0.3 |
| TcA(1193-IAP) | 2 | buffer/DDM | 11.8 | 2.0±0.1 |
| ABC(1193-IAP) | 2 | buffer/DDM | 11.8 | 1.5±0.1 |
| TcA(1193-IAP) | 2 | liposomes | 11.2 | 1.7±0.1 |

**Supplementary Table 3: Anisotropy Photon Distribution Analysis**. Fit-results of the displayed measurements/analysis in Supplementary Figure 7.

| | High anisotropy | Low anisotropy | Incubation time | Fraction of high anisotropy state [%] | Fraction of low anisotropy state [%] |
|---|---|---|---|---|---|
| TcA(2365-BFL) | 0.29 | 0.13 | 1 | 48.8 | 51.2 |
| | | | 160 | 78.1 | 21.9 |
| TcA(2384-BFL) | 0.29 | 0.14 | 1 | 39.3 | 60.7 |
| | | | 80 | 66.8 | 33.2 |
| TcA(1193-BFL) | 0.27 | 0.03 | 1 | 29.2 | 70.8 |
| | | | 50 | 67.5 | 32.5 |

**Supplementary Table 4: Overview of distance and FRET efficiency predictions for pore ejection (a) and shell opening (b).** Based on 3D structures for the prepore and pore state of the toxin a mean distance of the fluorophores was calculated using FRET Position Screening (FPS)[6]. Distances between TcA pentamer and TcB-TcC were screened to find informative FRET pairs.

**a - Pore ejection**

**Dependence of $\langle R_{DA} \rangle_E$ on the donor site in the TcA pentamer. TcA(914-BFL):TcB-TcC(1041-At647N)**

| | | subunit | | | | |
|---|---|---|---|---|---|---|
| **state** | | **A** | **B** | **C** | **D** | **E** |
| **prepore** | $E_{FRET}$ | 0.00 | 0.00 | 0.00 | 0.00 | 0.00 |
| | $\langle R_{DA} \rangle_E$ | 163 Å | 179 Å | 202 Å | 201 Å | 180 Å |
| **Pore** | $E_{FRET}$ | 0.25 | 0.01 | 0.00 | 0.00 | 0.02 |
| | $\langle R_{DA} \rangle_E$ | 58 Å | 105 Å | 147 Å | 143 Å | 95 Å |

Förster Radius $R_0$=49 Å

**b - Shell opening**

**Dependence of $\langle R_{DA} \rangle_E$ on the donor/acceptor site in the TcA pentamer for TcA(1279-At647N)**

| state | | A- B | A- C | A- D | A-E |
|---|---|---|---|---|---|
| **prepore** | $E_{FRET}$ | 0.85 | 0.26 | 0.26 | 0.84 |
| | $\langle R_{DA} \rangle_E$ | 50 Å | 80 Å | 80 Å | 51 Å |
| **Pore** | $E_{FRET}$ | 0.19 | 0.01 | 0.01 | 0.19 |
| | $\langle R_{DA} \rangle_E$ | 86 Å | 139 Å | 138Å | 85 Å |

Förster Radius $R_0$=67 Å (see https://www.atto-tec.com/fileadmin/user_upload/Katalog_Flyer_Support/R_0_-Tabelle_2018_web.pdf)

**Supplementary Table 5: Kinetics of biotinylated samples**. Additional measurements testing the influence of Biotin on the kinetic behavior of the toxins. Different concentrations of biotin and different fluorophores were tested. The quality/measurability of the samples is expressed as active molecules, meaning the ratio of molecules that changed their state/polarization. Using biotinylated sample resulted in an instant (not resolvable) and fast (within one data point) change of prepore to pore state. Slower kinetics were fitted using a single exponential term and resulting time constant is given. Typical data are shown in Supplementary Figure 9.

| Sample | Dye | Biotin | Active molecules | Reaction time [h] |
|---|---|---|---|---|
| TcA-1193 | At647N | none | 2% | 6.5±6 |
| TcA-1193 | BFL | none | 23% | 3h |
| TcA-1193 | At647N | none | 6% | 5±1.5 |
| TcA-1193 | At647N | 0.5x | 5% | 7h±5 |
| TcA-1193 | At647N | 2x | 15% | fast |
| TcA-1279 | At647N | 2x | 37% | fast |
| TcA-1193 | At647N | 50x | 0% | |
| TcA-2365 | BFL | none | 29% | 20h±3 |
| TcA -2384 | BFL | none | 17% | 20h±3 |
| TcA-2365 | At647N | none | 11% | 6h±3 |
| TcA-2365 | At647N | 2x | 11% | fast,<br>3h±1.5 |

**Supplementary Table 6**: **Cryo-EM data collection, refinement and validation statistics for TcA(1279-At647N), ABC(K567W/K2008W) and TcA(K1179W) at pH 11.2**.

| **Data collection and processing** | TcA(1279-At647N) | ABC(K567W/K2008W)<br>Combined dataset | TcA(K1179W) |
|---|---|---|---|
| Camera | K3 | K3 | Falcon III |
| Magnification | 81,000 | 81,000 | 120,000 |
| Voltage (kV) | 300 | 300 | 200 |
| Exposure time (s) | 3.0 | 3.0 | 3.0 |
| Total electron exposure (e–/Å$^2$) | 78 | 57 | 56 |
| Defocus range (μm) | 1.2 – 3.0 | 1.5 – 3.0 | 1.5 – 2.75 |
| Pixel size (Å) | 0.88 | 0.88 | 1.21 |
| Symmetry imposed | C5 | C5 | C5 |
| Initial particle images (no.) | 573,814 | 426,857 and 509,274 | 390,444 |
| Final particle images (no.) | 80,710 | 13,356 | 230,785 |
| Map resolution (Å)<br>FSC threshold | 3.8<br>0.143 | 3.2<br>0.143 | 2.9<br>0.143 |
| **Refinement** | | | |
| Initial model used (PDB code) | 6RW6 | 6RW6 | 6RW6 |
| Model resolution (Å)<br>FSC threshold | 3.6<br>0.5 | 3.3<br>0.5 | 2.9<br>0.5 |
| Model composition<br>Non-hydrogen atoms<br>Protein residues<br>Ligands | <br>84,390<br>10,610<br>0 | <br>84,445<br>10,610<br>0 | <br>84,420<br>10,610<br>0 |
| Mean B factors (Å$^2$)<br>Protein | <br>71.08 | <br>81.9 | <br>68.3 |
| R.m.s. deviations<br>Bond lengths (Å)<br>Bond angles (°) | <br>0.009<br>0.891 | <br>0.003<br>0.669 | <br>0.003<br>0.705 |
| Validation<br>MolProbity score<br>Clashscore<br>Rotamer outliers (%) | <br>1.48<br>6.04<br>0 | <br>1.35<br>6.17<br>0 | <br>1.5<br>8.4<br>0 |
| Ramachandran plot<br>Favored (%)<br>Allowed (%)<br>Disallowed (%) | <br>97<br>3<br>0 | <br>98<br>2<br>0 | <br>98<br>2<br>0 |
| Regions of model not built | N-terminus (aa 1-90), RBD B (aa 1309-1361 and 1493–1576), aa 1917-1939 and RBD C (aa 1383-1490) | N-terminus (aa 1-90), RBD B (aa 1309-1361 and 1493–1576), aa 1914-1946 and RBD C (aa 1383-1490) | N-terminus (aa 1-91), RBD B (aa 1309-1361 and 1493–1576), aa 1914-1944 and RBD C (aa 1383-1490) |

### Supplementary Notes

**Supplementary Note 1: Kinetic model for Shell opening (Figure 1e)**

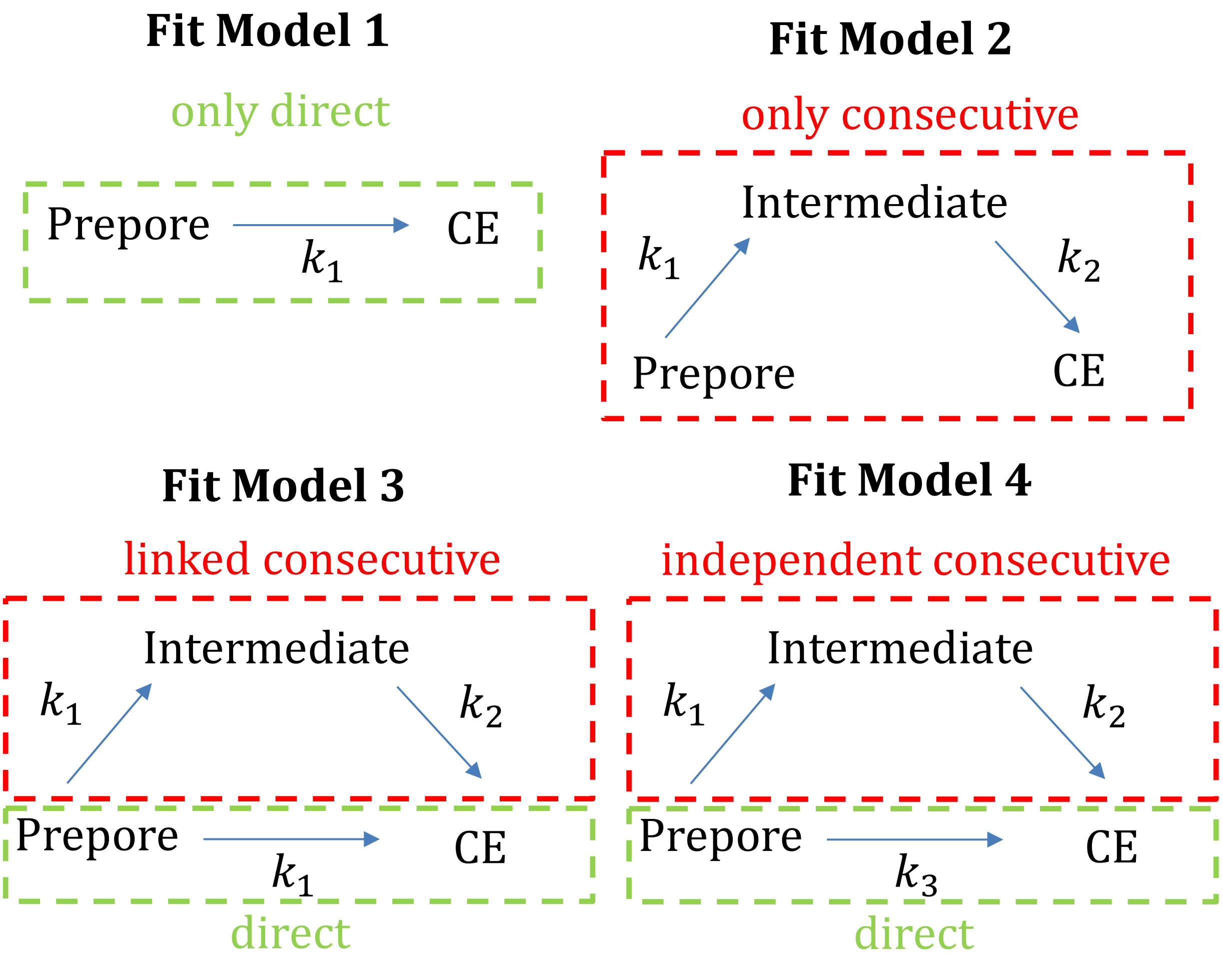

**Supplementary Figure SN1: Different fit models to resolve the kinetic of the prepore to channel ejected state.** Fit model 1 only assumes a simple exponential behavior in the kinetics of the prepore to channel ejected state. Fit model 2 is based on a consecutive step of the molecule to pass an intermediate state and then transitions to the channel ejected state. Fit model 3 allows a fraction of molecules to transition directly to the channel ejected state with the same rate they transition to an intermediate state. The rest of the molecules transition to the channel ejected state via the intermediate state. In fit model 4, the rate the molecule is allowed to directly transition to the CE state is independent of $k_1$. Data corresponding to each fit model is shown below.

Fit Model 1

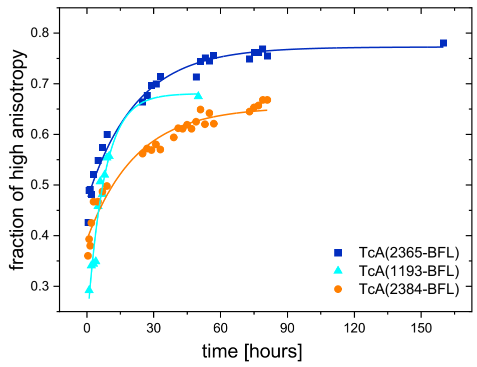

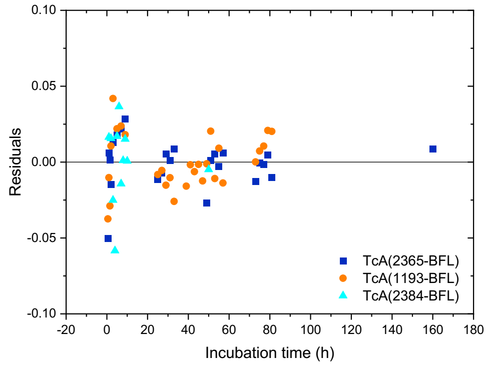

Fit Model 2

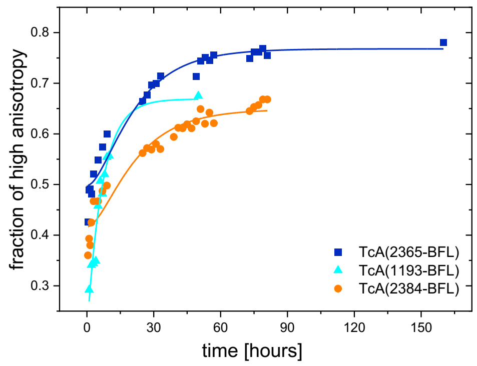

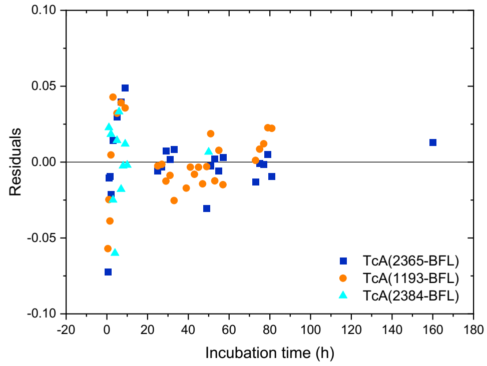

Fit Model 3

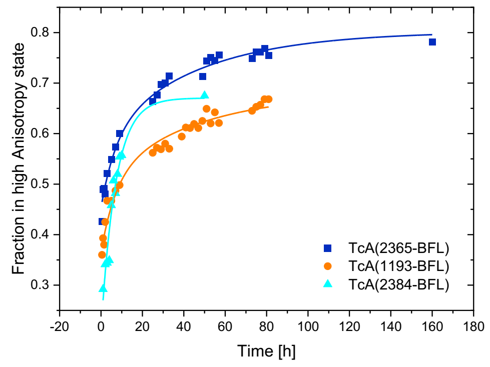

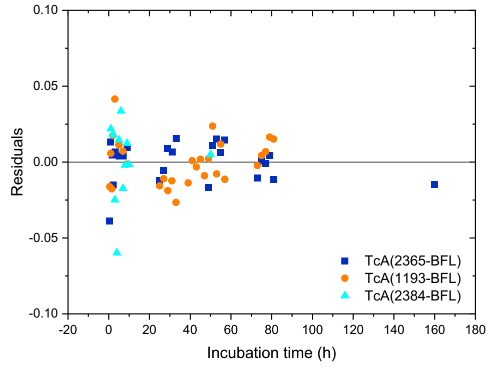

Fit Model 4

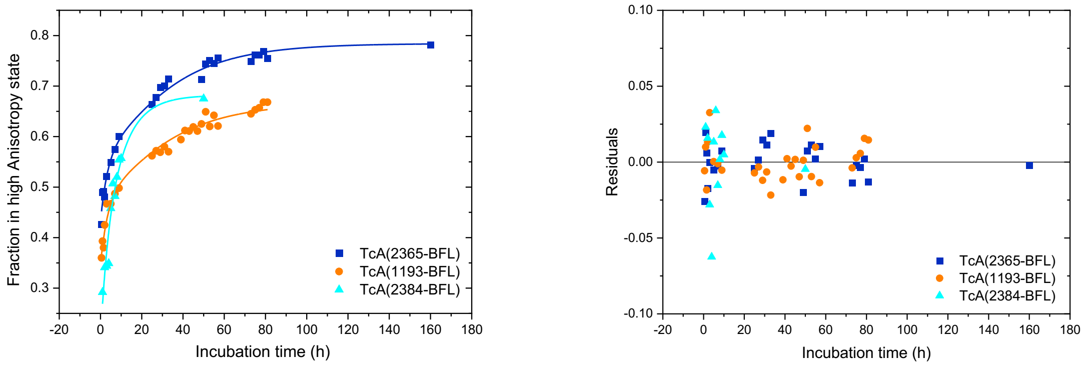

| Fit Model | reduced $\chi^2$ $[10^{-4}]$ shell opening | reduced $\chi^2$ $[10^{-4}]$ Pore injection | $k_{shell\ opening}$ [1/h] | $\sigma$ | $k_{pore}$ injection [1/h] | $\sigma$ |
|---|---|---|---|---|---|---|
| 1 | 3.134 | 8.295 | 0.04556 | 0.00375 | 0.13127 | 0.02158 |

| Fit Model | reduced $\chi^2$ $[10^{-4}]$ | $k_1$ [1/h] | $\sigma$ | $k_2$ [1/h] | $\sigma$ | $k_3$ [1/h] | $\sigma$ |
|---|---|---|---|---|---|---|---|
| 2 | 6.109 | 0.14355 | 0.02024 | 0.06004 | 0.00883 | - | - |
| 3 | 3.087 | 0.14176 | 0.01278 | 0.02182 | 0.0045 | - | - |
| 4 | 2.733 | 0.10061 | 0.01485 | 0.0365 | 0.00808 | 0.27339 | 0.07981 |

**Supplementary Note 2: Polarization-resolved Fluorescence Correlation Spectroscopy (pFCS)**

Polarization-resolved Fluorescence Correlation Spectroscopy (pFCS) was performed by correlating the parallel (p) and perpendicular (s) detected photons with each other and globally fitting the autocorrelation amplitude $G_{pp}(t_c)$ of parallel polarized fluorescence, the autocorrelation amplitude, $G_{ss}(t_c)$, of the perpendicular polarized fluorescence, and the crosscorrelation amplitude $G_{ps,sp}(t_c)$ of the parallel and perpendicular polarized fluorescence. We used a fit model with three bunching terms

$$G(t_c) = 1 + \frac{1}{N} \cdot \frac{1}{1 + \frac{t_c}{|t_d|}} \cdot \frac{1}{\sqrt{\left(1 + \frac{t_c}{(\frac{z_0}{\omega_0})^2 |t_d|}\right)}}$$

$$\cdot \left(1 - |C_{wb}| + |C_{wb}| \cdot e^{-\left(\frac{t_c}{|t_{wb}|}\right)} - |A_{gl}| + |A_{gl}| \cdot e^{-\left(\frac{t_c}{|t_{gl}|}\right)} - |B| + |B| \cdot e^{-\left(\frac{t_c}{|t_B|}\right)}\right)$$

with *N* the average number of molecules in the focus, $t_d$ the diffusion time, $\frac{z_0}{\omega_0}$ the gaussian shape, $C_{wb}$ the amplitude of the very fast dynamic wobbling term with the corresponding time constant, $t_{wb}$, $A_{gl}$ the amplitude of the triplet and global rotation of the molecule (prediction from HydroPro $t_A = [1 - 2.5]$ µs ) with corresponding time constant, $t_{gl}$, and B the amplitude with time constant $t_B$ of a third bunching term. The fitted parameter can be seen in Supplementary Tables SN1 - 4. The shown graph (Figure 4d) is an example plot for the parallel autocorrelation, which was fitted globally with the perpendicular- and cross-correlation. All data are shown in Supplementary Figures SN2 - 4 below.

**Supplementary Table SN1: Fit results for pFCS of TcA(1193-BFL-K1179W).** Values show the fit results for the autocorrelated signal of the parallel polarized fluorescence for different incubation times and pH environments of TcA(1193-BFL-K1179W). The first column shows the results for an incubation time of 0-400 s, the second one for 4700-5200 s. The last column shows the fit results for the measurement under pH 7 conditions.

| Sample | TcA(1193-BFL-K1179W) 0-400 s in pH 11 **$G_{pp}(t_c)$** | TcA(1193-BFL-K1179W) 4700-5200 s in pH 11 **$G_{pp}(t_c)$** | TcA(1193-BFL-K1179W) in pH 7 **$G_{pp}(t_c)$** |
|---|---|---|---|
| $\chi^2$ | 3.3158 | 4.4721 | 4.1057 |
| N | 0.2591 | 0.2905 | 0.2179 |
| $t_d$ [ms] | 4.3332 | 4.3878 | 3.3418 |
| $\frac{z_0}{\omega_0}$ | 2.0567 | 1.7059 | 1.7304 |
| $C_{wobble} = C_{wb}$ | 0.5786 | 0.3984 | 0.4890 |
| $t_{wb}$ [ms] | 0.0001 | 0.0006 | 0.0007 |
| $A_{global} = A_{gl}$ | 0.1655 | 0.2681 | 0.1264 |
| $t_{gl}$ [ms] | 0.0041 | 0.0068 | 0.0064 |
| B | 0.0383 | 0.0446 | 0.0249 |
| $t_B$ [ms] | 0.1683 | 0.1719 | 0.2390 |

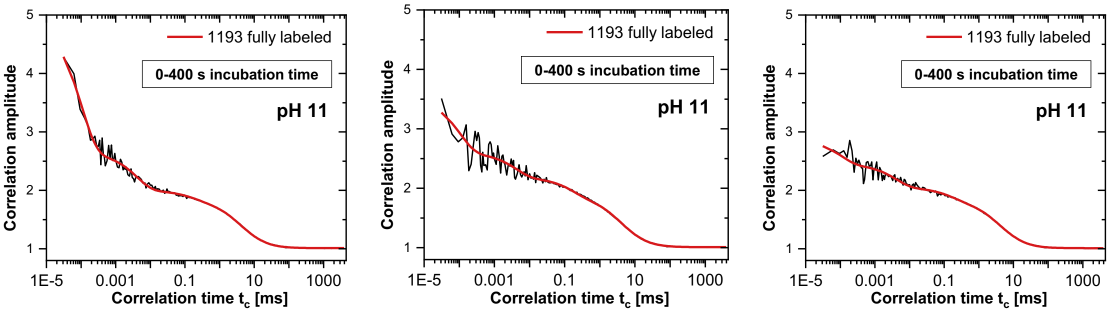

**Supplementary Figure SN2: pFCS of TcA(1193-BFL-K1179W) under pH 11 conditions for an incubation time of 0-400 s.** On the left side the correlation amplitude $G(t_c)$ of the autocorrelation of the parallel polarized fluorescence is shown, in the middle the correlation amplitude $G(t_c)$ of the autocorrelation of the perpendicular polarized fluorescence and on the right the correlation amplitude $G(t_c)$ of the cross correlation of the parallel and perpendicular polarized fluorescence.

**Supplementary Table SN2: Fit results of the pFCS of TcA(1193-BFL-K1179W) under pH 11 conditions for an incubation time of 0-400 s.** The first column shows the fit results of the correlation amplitude $G(t_c)$ of the autocorrelation of the parallel polarized fluorescence, the second column of the autocorrelation of the perpendicular polarized fluorescence and the third column of the cross correlation of the parallel and perpendicular polarized fluorescence.

| Sample | TcA(1193-BFL-K1179W) 0-400 s in pH 11 parallel autocorrelation **$G_{pp}(t_c)$** | TcA(1193-BFL-K1179W) 0-400 s in pH 11 perpendicular autocorrelation **$G_{ss}(t_c)$** | TcA(1193-BFL-K1179W) 0-400 s in pH 11 crosscorrelation **$G_{ps,sp}(t_c)$** |
|---|---|---|---|
| $\chi^2$ | 3.3158 | 4.0828 | 3.4052 |
| N | 0.2591 | 0.3988 | 0.5378 |
| $t_d$ [ms] | 4.3332 | 4.3332 | 4.3332 |
| $\frac{z_0}{\omega_0}$ | 2.0567 | 2.0567 | 2.0567 |
| $C_{wobble} = C_{wb}$ | 0.5786 | 0.3657 | 0.2272 |
| $t_{wb}$ [ms] | 0.0001 | 0.0001 | 0.0001 |
| $A_{global} = A_{gl}$ | 0.1655 | 0.1664 | 0.2137 |
| $t_{gl}$ [ms] | 0.0041 | 0.0041 | 0.0041 |
| B | 0.0383 | 0.1187 | 0.1093 |
| $t_B$ [ms] | 0.1683 | 0.1683 | 0.1683 |

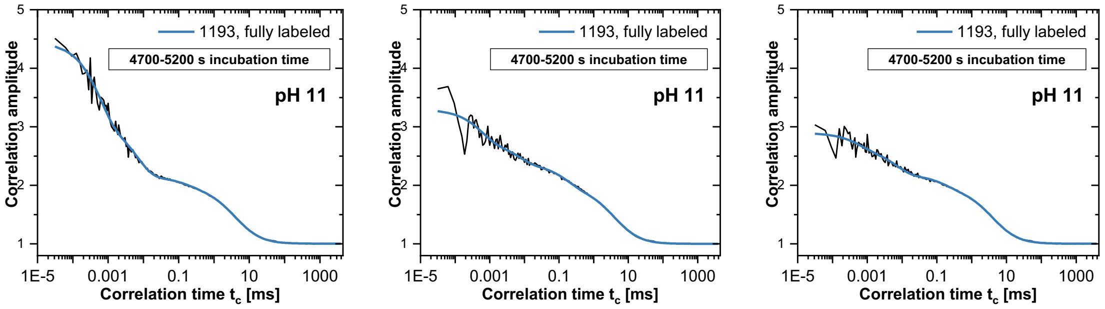

**Supplementary Figure SN3: pFCS of TcA(1193-BFL-K1179W) under pH 11 conditions for an incubation time of 4700-5200 s.** On the left side the correlation amplitude $G(t_c)$ of the autocorrelation of the parallel polarized fluorescence is shown, in the middle the correlation amplitude $G(t_c)$ of the autocorrelation of the perpendicular polarized fluorescence and on the right the correlation amplitude $G(t_c)$ of the cross correlation of the parallel and perpendicular polarized fluorescence.

**Supplementary Table SN3: Fit results of the pFCS of TcA(1193-BFL-K1179W) under pH 11 conditions for an incubation time of 4700-5200 s.** The first column shows the fit results of the correlation amplitude $G(t_c)$ of the autocorrelation of the parallel polarized fluorescence, the second column of the autocorrelation of the perpendicular polarized fluorescence and the third column of the cross correlation of the parallel and perpendicular polarized fluorescence.

| Sample | TcA(1193-BFL-K1179W) 4700-5200 s in pH 11 parallel autocorrelation **$G_{pp}(t_c)$** | TcA(1193-BFL-K1179W) 4700-5200 s in pH 11 perpendicular autocorrelation **$G_{ss}(t_c)$** | TcA(1193-BFL-K1179W) 4700-5200 s in pH 11 crosscorrelation **$G_{ps,sp}(t_c)$** |
|---|---|---|---|
| $\chi^2$ | 4.4721 | 4.3209 | 4.0583 |
| N | 0.2905 | 0.4351 | 0.5276 |
| $t_d$ [ms] | 4.3878 | 4.3878 | 4.3878 |
| $\frac{z_0}{\omega_0}$ | 1.7059 | 1.7059 | 1.7059 |
| $C_{wobble} = C_{wb}$ | 0.3984 | 0.2410 | 0.1397 |
| $t_{wb}$ [ms] | 0.0006 | 0.0006 | 0.0006 |
| $A_{global} = A_{gl}$ | 0.2681 | 0.1628 | 0.2353 |
| $t_{gl}$ [ms] | 0.0068 | 0.0068 | 0.0068 |
| B | 0.0446 | 0.1663 | 0.1127 |
| $t_B$ [ms] | 0.1719 | 0.1719 | 0.1719 |

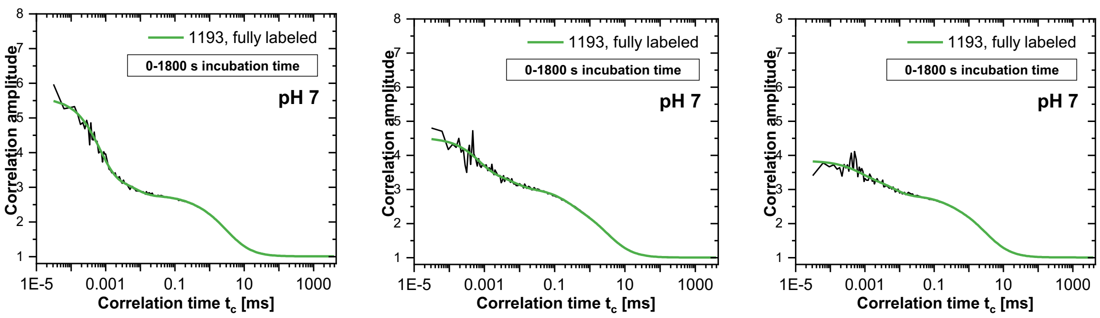

**Supplementary Figure SN4: pFCS of TcA(1193-BFL-K1179W) under pH 7.** On the left side the correlation amplitude $G(t_c)$ of the autocorrelation of the parallel polarized fluorescence is shown, in the middle the correlation amplitude $G(t_c)$ of the autocorrelation of the perpendicular polarized fluorescence and on the right the correlation amplitude $G(t_c)$ of the cross correlation of the parallel and perpendicular polarized fluorescence.

**Supplementary Table SN4: Fit results of the pFCS of TcA(1193-BFL-K1179W) under pH 7 conditions.** The first column shows the fit results of the correlation amplitude $G(t_c)$ of the autocorrelation of the parallel polarized fluorescence, $\boldsymbol{G_{pp}(t_c)}$, the second column of the autocorrelation of the perpendicular polarized fluorescence, $\boldsymbol{G_{ss}(t_c)}$**,** and the third column of the cross correlation of the parallel and perpendicular polarized fluorescence, $\boldsymbol{G_{ps,sp}(t_c)}$.

| Sample | TcA(1193-BFL-K1179W) in pH 7 parallel autocorrelation **$G_{pp}(t_c)$** | TcA(1193-BFL-K1179W) in pH 7 perpendicular autocorrelation **$G_{ss}(t_c)$** | TcA(1193-BFL-K1179W) in pH 7 crosscorrelation **$G_{ps,sp}(t_c)$** |
|---|---|---|---|
| $\chi^2$ | 4.1057 | 4.5714 | 2.5664 |
| N | 0.2179 | 0.2839 | 0.352 |
| $t_d$ [ms] | 3.3418 | 3.3418 | 3.3418 |
| $\frac{z_0}{\omega_0}$ | 1.7304 | 1.7304 | 1.7304 |
| $C_{wobble} = C_{wb}$ | 0.4890 | 0.2779 | 0.1571 |
| $t_{wb}$ [ms] | 0.0007 | 0.0007 | 0.0007 |
| $A_{global} = A_{gl}$ | 0.1264 | 0.1386 | 0.1975 |
| $t_{gl}$ [ms] | 0.0064 | 0.0064 | 0.0064 |
| B | 0.0249 | 0.1386 | 0.0851 |
| $t_B$ [ms] | 0.2390 | 0.2390 | 0.239 |

### Supplementary Note 3

The measurement of the trace of the transition of TcA(1279-At647N-Biotin) yielded different parameters: (*1*) the pre-transition time (see Supplementary Figure SN5a), defined as the time the signal stays constant after the pH change until a change in the signal occurred. The pH change was detected in the signal as an instant brightness change of the monitored dyes. Next (*2*), is the transition time (see Supplementary Figure SN5b), which is defined as the time the molecule needs to go from one state to the other. In terms of the signal, this is the time the signal needs to go from, in this case, a low anisotropy to a high anisotropy level. These parameters were read from the signal as shown in Supplementary Figure SN6, where the transition time was taken as the time the signal showed the maximum change in the trace based on the derivative of the averaged signal. The sum of the pre-transition time and transition time is defined as the total reaction time (*3*) (see Supplementary Figure SN5c).

The pre-transition and reaction times showed monoexponential behavior in their normalized distribution, which was fitted using (1)

$$[destabilized\ shell] = \left(1 - e^{-\frac{t}{\tau_1}}\right) + y$$

where *y* was used as an offset and τ is the relaxation time of the state.

**a**

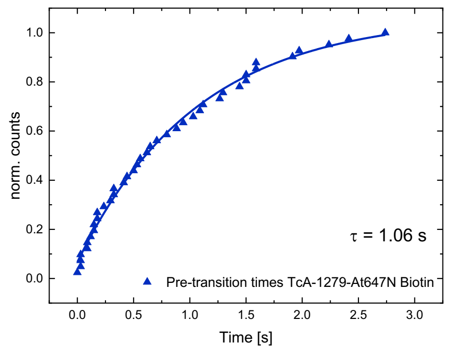

**b**

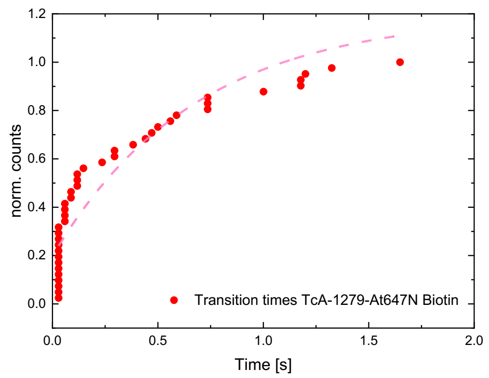

**c**

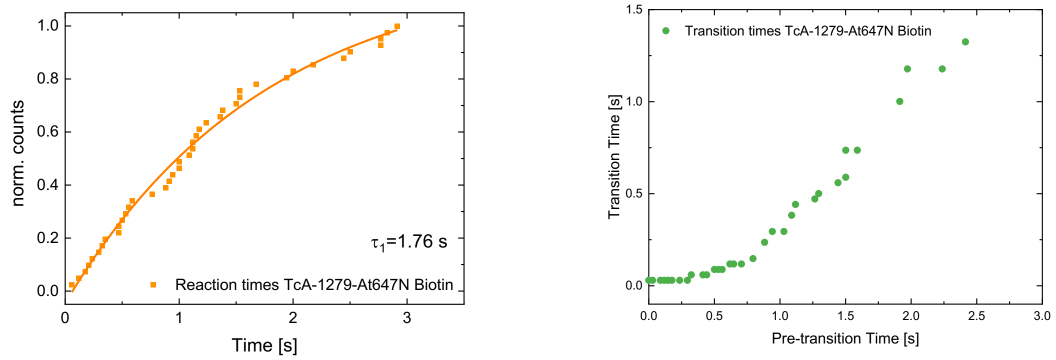

**Supplementary Figure SN2: Dynamic behavior of TcA(1279-At647N-Biotin) under pH change. a:** Normalized distribution of the pre-transition times defined as the time after pH change until the signal starts to permanently change. The monoexponential behavior was fitted using equation (1) **b** Normalized distribution of the transition times defined as the time the signal changed from low to high **c** Normalized distribution of the reaction times, defined as the sum of the pre-transition and transition times.

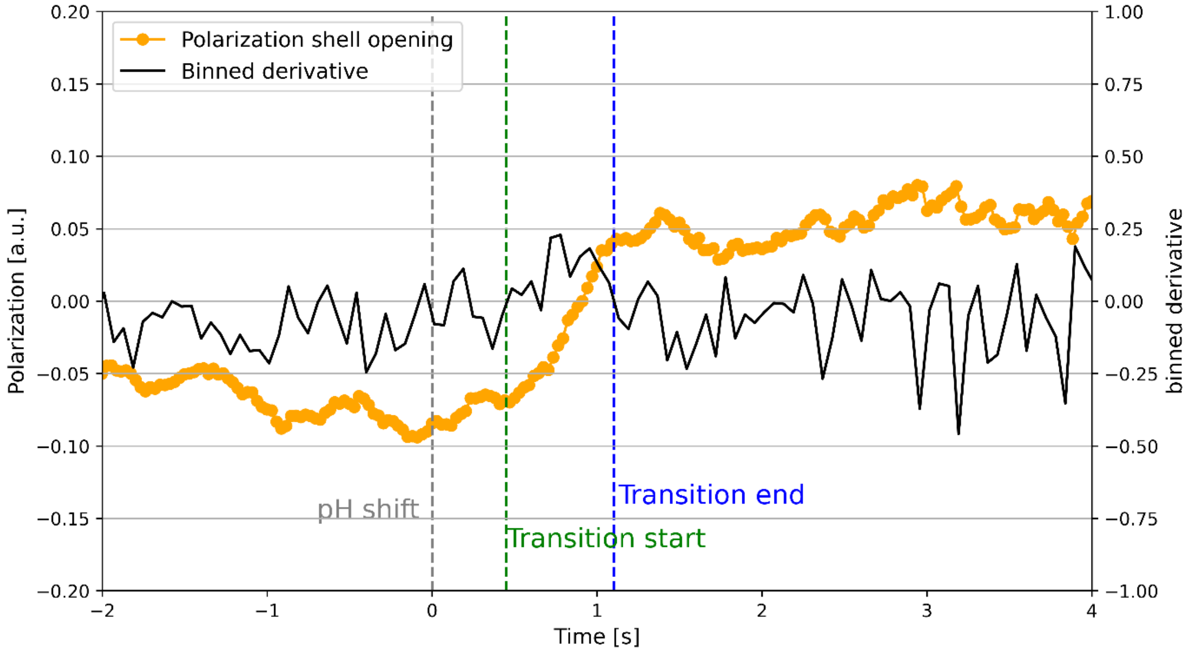

**Supplementary Figure SN3: Example of the detection of pre-transition and transition times.** The time after pH change (t = 0, grey dashed line) until the signal changes before the transition starts (green dashed line) is defined as the pre-transition time. The transition time is defined as the time the signal needs until it is permanently changed. Binned derivative was calculated as the change of between two signals over the interval, which was binned afterwards with a binning size of two (black line).

### Supplementary Methods

#### Single Molecule confocal MFD

Single-molecule events were identified using a burst search algorithm according to Fries *et al.*, 1998[7], a Lee filter, a threshold of 0.2 ms and a minimum of 60 photons per burst. For anisotropy Photon Distribution Analysis (PDA), the whole trace including all bursts was analyzed. For heteroFRET analysis, molecules with a stoichiometry between *S*=0.5 to *S*=0.8 were selected. The labeling geometry of five possible labeling spots for TcA(914-BFL)-Biotin:TcB(1041-Atto647N)-TcC and a labeling scheme of 5:1 donor to acceptor led to a higher than usual value of the stoichiometry, where multiple donors were attached to the molecule with only one acceptor. However, only one of the labeling positions had a distance to the acceptor where FRET occurs, giving the possibility to qualitatively measure FRET-efficiency-derived distances.

Static FRET lines as described by Kalinin *et al.*, 2012[6] were used for consistency and quality control. A static FRET line relates the lifetime of the donor in the presence of an acceptor, to the resulting FRET efficiency. For the Tc toxin, no dynamic exchange between conformational states was expected. This matched experimental data, where the double-labeled FRET population of the Tc toxin in prepore and pore states on the $\langle \tau_{D(A)} \rangle_F - E$ is located on the static FRET line (Supplementary Figure 11).

#### Correction factors of intensity based confocal MFD

Correction factors were estimated from the spectra of the used dyes obtained from the web/manufacturer (Corp., C.T. *Spectra Viewer*. 2021; Available from: https://www.chroma.com/spectra-viewer) and the spectra of the optical components based on Hellenkamp *et al.*, 2018[8]. The spectral donor crosstalk of the donor α is defined as the detection efficiency ratios $g$ of the red to green detection channel while donor excitation:

$$\alpha = \frac{g_{R|D}}{g_{G|D}}$$

The direct excitation of the acceptor with the donor laser δ is based on the ratio of the cross-sections of the acceptor excited with the donor laser $\sigma_{A|G}$ and with the acceptor laser $\sigma_{A|R}$ and the laser intensities:

$$\delta = \frac{\sigma_{A|G}}{\sigma_{A|R}} \frac{I_{Dex}}{I_{Aex}}$$

To monitor the labeling ratio of donor to acceptor of the molecule, the excitation flux was normalized using

$$\beta = \frac{\sigma_{A|R}}{\sigma_{D|G}} \frac{I_{Aex}}{I_{Dex}}$$

which follows that, molecules labeled with a 1:1 ratio have a stoichiometry value $S = 0.5$. Due to the complex labeling geometry of possible multilabeling, an additional ALEX-2CDE filter[9] was applied to remove further unwanted contributions of single-labeled molecules and reduce the effect of photobleaching. Differences in the quantum yields and the detection efficiencies of the different spectral ranges lead to $\gamma$ correction factor defined as:

$$\gamma = \frac{g_{R|A}}{g_{G|D}} \frac{{}^{eff}\Phi_{F,A}}{{}^{eff}\Phi_{F,D}}$$

with the dark-state-corrected effective quantum yields of the acceptor and donor.

**Intensity based spectroscopic parameters**

Raw intensities for green and red signal after donor and acceptor excitation were corrected from background taken from a buffer-only measurement:

$$\begin{aligned} {}^{ii}I_{\mathrm{Dem|Dex}} &= {}^{i}I_{\mathrm{Dem|Dex}} - I^{(BG)}_{\mathrm{Dem|Dex}} \\ {}^{ii}I_{\mathrm{Aem|Dex}} &= {}^{i}I_{\mathrm{Aem|Dex}} - I^{(BG)}_{\mathrm{Aem|Dex}} \\ {}^{ii}I_{\mathrm{Aem|Aex}} &= {}^{i}I_{\mathrm{Aem|Aex}} - I^{(BG)}_{\mathrm{Aem|Aex}} \end{aligned}$$

and additionally corrected from the calculated correction parameters to the fluorescence signal

$$\begin{aligned} F_{\mathrm{D|D}} &= \gamma\, {}^{ii}I_{\mathrm{Dem|Dex}} \\ F_{\mathrm{A|A}} &= \frac{1}{\beta}\, {}^{ii}I_{\mathrm{Aem|Aex}} \\ F_{\mathrm{A|D}} &= {}^{ii}I_{\mathrm{Aem|Dex}} - \alpha\, {}^{ii}I_{\mathrm{Dem|Dex}} - \delta\, {}^{ii}I_{\mathrm{Aem|Aex}} \end{aligned}$$

which could then be used to calculate FRET efficiency *E* and the stoichiometry *S*:

$$E = \frac{F_{\mathrm{A|D}}}{F_{\mathrm{D|D}} + F_{\mathrm{A|D}}}$$

$$S = \frac{F_{\mathrm{D|D}} + F_{\mathrm{A|D}}}{F_{\mathrm{D|D}} + F_{\mathrm{A|D}} + F_{\mathrm{A|A}}}$$

Förster Radius was calculated using the spectral overlap of the emission of the donor and the excitation of the acceptor dye $J$. For the dipole orientation factor $\kappa^2$ the isotropic average of 2/3 was assumed, for the refractive index a commonly used value of $n = 1.4$ was taken. Following this, the FRET efficiency averaged distance[10] $\langle R_{DA} \rangle_E$:

$$\langle R_{DA} \rangle_E = R_0 (E^{-1} - 1)^{1/6}$$

**FRET Position Screening (FPS)**

In order to design a label scheme with dyes positioned in the FRET-sensitive range to measure conformational change due to shell opening and pore ejection, a FRET position screening was applied following Dimura *et al.*, 2020[11], using in house software. In short, it applies position-wise so called Accessible Volumes (AVs)[6] to multiple positions on the molecule while calculating the expected distance $\langle R_{DA} \rangle_E$ (SI Figure 8). To do so, the first position for the donor dye was set to a position identified based on other experiments. Then, subsequent positions were selected, avoiding contact with surrounding amino acids which would quench the dyes. Subsequently, the labelled protein was tested using smMFD, resulting in TcA(1279-Atto647N) and TcA(914-BFL:TcB(1041-Atto647N)-TcC as the best choices in terms of good label ratio for measuring shell opening and pore ejection, respectively (SI Figure 8).

**Anisotropy Photon Distribution Analysis**

Anisotropy states and fractions were estimated via Photon Distribution Analysis (PDA). Anisotropy changed based on different phenomena. For TcA-2365-Bdp and TcA-2384-BdP, an environmental change due to prepore-to-pore transition restricted the dye's movement in pore state, leading to a higher anisotropy. In case of TcA-1193-Bdp and TcA-1279-Atto647N, the anisotropy in pore state was decreased due to a higher distance between the dyes and therefore the reduction of homoFRET efficiency. Based on previous studies[12,13], anisotropy PDA was performed. In this regard, the sum of the probability of every combination of the ratio $S_{||}/S_{\perp}$ was

$$P\left(\frac{S_{||}}{S_{\perp}}\right) = \sum_{(S_{||}/S_{\perp})_i} P(S_{||}, S_{\perp}, B_{||}, B_{\perp})$$

where $S_{||}$ is the parallel polarized signal and $S_{\perp}$ the perpendicular polarized signal, $B$ their corresponding background signal. Experimental data followed Poisson statistics and was fitted using a maximum entropy method described in the given references[8,9]. The scatter-corrected fluorescence followed:

$$r_s = \frac{G\left(S_{||} - \langle B_{||} \rangle\right) - \left(S_{\perp} - \langle B_{\perp} \rangle\right)}{G\left(S_{||} - \langle B_{||} \rangle\right)(1 - 3l_2) + \left(S_{\perp} - \langle B_{\perp} \rangle\right)(2 - 3l_1)}$$

with the correction factors $l_1$ and $l_2$ describing the mixing of the polarizations in the microscope objective and $G$ the ratio of detection in parallel and perpendicular channel, obtained from a free dye measurement. PDA analysis was performed globally over three different time steps, 1 ms, 2ms and 3 ms. The experimental numbers of time windows with a particular $r_s$ were fitted using a 2-state model with a high and low anisotropy state. Fitting involved minimization of reduced $\chi^2$-values. The resulting time-dependent fractions of high and low anisotropy states described the kinetic behavior of the toxin.

### Total Internal Reflection Fluorescence (TIRF)

Total Internal Reflection Fluorescence (TIRF) uses an evanescent field coming from the high refractive index difference of the TIRF objective and the sample. This results in a total reflection of an incoming laser beam in case it enters the medium with an angle higher than a critical angle based on the difference of the refractive indices following Snell's law. Thus, it makes the technique ideal for samples immobilized to the surface, because the laser beam only enters typically around 100 nm of the solution, leading to a high signal to noise ratio[14].

The correction factors were calculated based on the spectral properties of the dyes and applied globally for every molecule. The following correction factors were used:

| | **HeteroFRET Alexa488/Atto647N** | **homoFRET Atto647N/Atto647N** |
|---|---|---|
| $\alpha$ | 0.016 | 0 |
| $\delta$ | 0.02 | 0 |
| $\frac{g_{R\|A}}{g_{G\|D}}$ | 1.25 | 1 |
| $\frac{^{eff}\Phi_{F,A}}{^{eff}\Phi_{F,D}}$ | $\frac{0.65}{0.8} = 0.81$ | 1 |
| $\gamma$ | 1.02 | 1 |

For heteroFRET efficiency, background corrected fluorescence *F* was used following equations described in section intensity-based MFD.

In case of the homoFRET anisotropy assay, one has to take into account that the high numerical aperture of the TIRF objective influences the polarization of the laser beam, leading to a mixture of polarizations. Additionally, the laser beam enters the solution at an angle greater than a critical angle for total reflection. This effect was studied in detail and a shift to lower anisotropy values was observed (see examples[15,16]). Additionally, due to a high amount of different labeling sites of the biotin on the Tc toxin, the initial orientation and therefore polarization values of the toxin were distributed. However, in this assay the difference of polarization of the signal before and after the transition from prepore to pore was of interest. Therefore, we applied a polarization offset based on the mean polarization value before pH change $\langle P(ph7)\rangle$ to every trace, shifting the polarization value before the pH change to *P*=0. The polarization value resulted in:

$$P = \frac{F_{||} - F_{\perp}}{F_{||} + F_{\perp}} - \langle P(ph7)\rangle$$
